# Development of Machine Learning-based QSAR Models for the Designing of Novel Anti-cancer Therapeutics Against Malignant Glioma

**DOI:** 10.1101/2024.08.19.608549

**Authors:** Fareed Asaad, Mehreen Zaka, Serdar Durdağı

**Affiliations:** Computational Biology and Molecular Simulations Laboratory, Department of Biophysics, School of Medicine, Bahcesehir University, Istanbul, Türkiye; Department Of Natural and Applied Sciences, College of Arts and Sciences, American University of Iraq Baghdad (AUIB); Molecular Therapy Laboratory, Department of Pharmaceutical Chemistry, School of Pharmacy, Bahcesehir University, Istanbul, Türkiye; Lab for Innovative Drugs (Lab4IND), Computational Drug Design Center (HITMER), Bahcesehir University, Istanbul, Türkiye

**Author notes:** (SD). Equal contribution (Fareed Asaad, Mehreen Zaka).

**Keywords:** ML-based QSAR models, glioma, GBM, molecular simulations, IDH1 mutant inhibitors

## Abstract

In the early drug design and discovery phase, virtual screening of diverse small molecule libraries is crucial. Machine learning (ML)-based algorithms have made this process easier and faster. In this study, we have applied ML-based algorithms to generate the QSAR models for virtual screening. The aim of study is to design the statistically significant models for the screening of small molecule libraries to identify the novel hits against IDH1 mutant receptor crucial for glioblastoma multiforme (GBM). To construct the models, we have used both cell lines data (U87 and U251 cells) and the inhibitors of IDH1 mutant reported in the literature and used the pIC_50_ activity data to train our models. Furthermore, ligand-based 3D QSAR models and structure-based pharmacophore models were also constructed and validated.

## Introduction

Gliomas are brain tumors starting in the glial cells and they are classified as astrocytic, oligodendroglial, oligoastrocytic (mixed) or ependymal according to the morphological and activity characteristics of the cells under the microscope (Holland et al., 2000; Maher et al., 2001; Schwartzbaum et al., 2006; Agnihotri et al., 2013). According to World Health Organization (WHO) 2007 criteria, besides cell morphology, mitotic activity, microvascular proliferation, anaplasia, and necrotic activity are also evaluated and classical classification is done (Louis et al., 2007). In this classification, tumors are classified from low grade (slow growing) to high grade (fast growing), from WHO Grade I to Grade IV. Gliomas are common primary brain tumors of glial origin (representing 40-50% of all brain tumors) with highly aggressive and invasive properties. They are the most commonly occurring tumors of the central nervous system (CNS), which account for almost 80% of all malignant primary tumors of brain (Schwartzbaum et al., 2006; Agnihotri et al., 2013).

Gliomas constitute the second most common cause of death from intracranial disease after stroke (Viel et al., 2013) and they have dynamic heterogeneous tumor tissues consisting of tumor cells with uncontrolled proliferation inducing angiogenesis and escaping the host immune response. Their intratumoral heterogeneity and diversity of tumor-associated molecular defects lead to treatment resistance and tumor recurrence triggered by cancer stem-like cells (CSCs).

The present therapeutic approaches aim to target tumor drivers such as specific gene defects and oncogenic signaling pathways, but these approaches have low efficiency (Vasilogiannakopoulou et al., 2018). Gliomas show diverse anatomic and metabolic features that manifest differently across patients, ranging in aggressiveness from low-grades to high-grades. Around 50% of glioma patients are diagnosed with glioblastoma multiforme (GBM), the WHO grade IV glioma, which is the most aggressive brain tumor in adults (Louis et al., 2007). The difficulty in treating gliomas is primarily due to the invasion of tumor-containing cell populations. Despite extensive treatments including surgical resection, radiotherapy and chemotherapy, GBMs tend to be recurrent and fatal with a median survival of approximately one year (Burnet et al., 2007; Ohka et al., 2012; Thakkar et al., 2014). Studies showed that GBMs display the worst prognosis with only 10% of diagnosed patients surviving for 5 years.

Isocitrate dehydrogenase 1 (IDH1) is a critical metabolic enzyme involved in the tricarboxylic acid cycle. This enzyme catalyzes the oxidative decarboxylation of isocitrate acid to α-ketoglutaric (α- KG) in an NADP^+^-dependent manner by using divalent magnesium ion (Jiao et al., 2016). Somatic mutations of IDH1 have been frequently identified in many types of cancer, including approximately 80% of grade II-III gliomas, nearly 45% of GBM, and 33%-50% of adult primitive neuroectodermal tumors (Dang et al., 2009). IDH1 is also scored as the second attractive target for GBM after epidermal growth factor receptor (EGFR) according to the webserver Open Targets (https://www.opentargets.org/). Besides brain tumors, IDH1 mutations have also been detected in other cancers including acute myeloid leukemia (Parsons et al., 2008), colorectal cancer (Xu et al., 2011), and prostate cancer with low frequencies (Hartmann et al., 2009). Key amino acid residue Arg132 is the most common mutation (usually to histidine) in IDH1 which is located in the catalytic pocket (Dang et al., 2009). Specific mutations belong to heterozygous missense mutations and lead to a new form of IDH1 catalytic activity, which convert α-KG into an oncometabolite D2-hydroxyglutarate (D2-HG) (Dang et al., 2009). The oncometabolite (D2-HG) is associated with tumorigenesis, which impairs hematopoietic differentiation and promotes leukemia by inducing the hypermethylation of histone and chromatin and preventing cell differentiation (Figueroa et al., 2010; Xu et al., 2011). Due to the IDH1 mutation, high levels of D2-HG are formed that promote the occurrence and development of cancers.

Mutant IDH1 has become a very attractive therapeutic target in the field of antitumor drug discovery, and several pharmaceutical companies have attempted to develop novel small molecule inhibitors against mutant IDH1. So far, several small molecule inhibitors targeting mutant IDH1 enzymes have been developed (Rohle et al., 2013; Davis et al., 2014; Deng et al., 2015; Kim et al., 2015; Okoye-Okafor et al., 2015; Law et al., 2016; Chaturvedi et al., 2017; Xie et al., 2017; Nakagawa et al., 2019; Caravella et al., 2020; Konteatis et al., 2020). Some of these have been studied in various preclinical models, and some are currently being evaluated in phase I/II clinical studies for different tumor pathologies with IDH1 enzyme mutations. AG-120 (ivosidenib) as the only mutant IDH1 inhibitor in clinic approved by the FDA that has shown encouraging clinical benefits with a total overall response rate of 42% for advanced hematological malignancies (Foran et al., 2019).

In the current study, we have combined the machine learning (ML) algorithms and physics-based molecular simulations techniques (both ligand-based and target-driven based approaches) for the identification of novel hit compounds against GBM. For this aim, we have developed the novel models using ML-based pharmacophore modeling, classical pharmacophore modeling, and structure-based (i.e., e-Pharmacophore) hypothesis.

## Methods

In this study, two model construction methods were used to generate pharmacophore models with good predictive ability: Ligand-based and structure-based. In the ligand-based method, two approaches were used for model generation: (i) ML-based quantitative structure-activity relationships (QSAR) models by AutoQSAR module of the Maestro molecular modeling package which is a ML application to build, validate and deploy QSAR models; and (ii) classical pharmacophore model hypothesis building by PHASE, which is a highly flexible system for common pharmacophore identification and evaluation, generation of 3D QSAR models. For the structure-based approach, e-Pharmacophore module of the Maestro was used. This method uses the energy terms calculated by the Glide XP scoring function and combines the pharmacophore perception and database screening to rank the pharmacophore features.

### ML-based model construction

AutoQSAR is a ML-based algorithm that builds and applies QSAR models (Dixon et al., 2016). AutoQSAR uses the 2D information of compounds and experimental activity data (e.g., IC_50_) to generate the predictive models. Generally, the QSAR models show the linear or non-linear statistical correlation between the molecular descriptors and experimental activity. Statistical ML methods are exploited to compute the descriptors and fingerprints for predictive models’ development. The accuracy of the models is validated by using several parameters, including Q^2^, R^2^, root mean square error (RMSE), standard deviation (SD), and ranking score values (Dixon et al., 2016). Recent advancements in the application of ML for QSAR model development have demonstrated the effectiveness of AutoQSAR in predicting the activity of new compounds (Zeng et al., 2024).

### Ligand preparation

LigPrep module (Ligprep, Schrodinger LLC, 2018) with the OPLS3 forcefield was used to prepare ligands. An issue to be considered is the ionization of the ligand in physiological environments. At the pH of 7, the Epik module was used for potential ionization states.

### 3D QSAR Pharmacophore model construction

PHASE module from Maestro molecular modeling package (Dixon, 2006) was used to generate pharmacophore hypotheses and 3D QSAR analysis. A common pharmacophore hypothesis was detected from a set of chemical features resembling the spatial arrangement of the data set. These chemical features are categorized in PHASE as hydrogen bond acceptor (A), hydrogen bond donor (D), aromatic ring (R), hydrophobic groups (H), positively charged groups (P), and negatively charged groups (N). To generate the ligand-based 3D QSAR models, GBM inhibitors were collected from ChEMBL database (https://www.ebi.ac.uk/chembl/) with experimental activity (IC_50_) against U-87 and U-251 glioblastoma cell lines. The diverse dataset was created, ranging pIC_50_ values from 3 to 10. 169 compounds were used to construct QSAR hypothesis from U-87 cell line results and 87 compounds were used from U-251 cell line results. All the ligands were prepared using LigPrep, and conformers were generated using MacroModel. For each ligand, up to 1000 conformers were generated. Any compound which has pIC_50_ value ≥7 was set as active and any compound that has pIC_50_ value ≤4.5 was set as inactive. These settings gave 28 active, 24 inactive compounds for U-87 model, and 21 active molecules and 15 inactive molecules for U-251 cell line model. 3 PLS factors were used, and pharmacophore-based hypotheses were generated. Top-2 models were chosen, namely; ADHRRR, and AAAHH, based on the R^2^ and Q^2^ along with the statistics with the external test set.

### Protein Preparation

Protein Preparation module in Schrödinger 2018 (Schrödinger, LLC, New York, NY, 2018) was used for protein preparation. Water molecules around the binding pocket (<5 Å) were kept during the protein preparation. The PROPKA (pH of 7.4) and OPLS3 force fields were used for protonation states, structural optimization and minimization, respectively. All the default settings in PrepWizard were used for protein preparation.

### Energetically optimized structure-based pharmacophores

The e-Pharmacophore module in the Schrodinger suite was used for structure-based pharmacophore model generation. Both ligand-based and structure-based methods are an integral part of the drug discovery pipeline. For fast screening of small molecule libraries, 3D pharmacophore modeling can be used. On the contrary, structure-based approaches are helpful to get insights about target protein and active binding sites. The e-Pharmacophore method offers the advantages of both structure and ligand-based techniques by generating energetically optimized pharmacophores (Salam et al., 2009). The receptor-ligand cavity is exploited initially to refine the ligand pose, and the Glide XP scoring function is used to compute the energy terms. Afterward, pharmacophore sites are generated, and Glide XP energies from the atoms are summed up. The energies are ranked, and the best sites are utilized to create the final pharmacophore hypothesis.

## Results

### Construction of the models using ML-based algorithm

GBM inhibitors against U-87 and U-251 glial cell lines were downloaded from the ChEMBL database with IC_50_ values and were used for ML-based QSAR model generation. The IC_50_ activity values were converted into pIC_50_ values because the nature of potency values is logarithmic. Two models were generated using two different cell lines inhibitors. For the first model (U-87), 172 molecules were prepared using the LigPrep module at the neutral pH. In this approach, the internal validation function of the traditional AutoQSAR program was exploited; molecules were divided into two sets, 67% for model generation (116 compounds) and 33% for internal validation set (56 compounds). 116 compounds were subdivided into a 70% (82 compounds) training set and 30% (34 compounds) test set for the model generation. (Tables S1 and S2) To validate the model and verify its prediction power, 169 molecules which are tested experimentally at U-87 cells from ChEMBL were used as an external test set (i.e., true unknown compounds); those molecules did not participate in the model formation and were different from the training and test set. (Table S3) 9 models were generated from this model set, and *kpls_radial_15* was selected as the top model based on the model’s external test set prediction power. Table 1 shows the statistical results of the selected model. Figure 1 shows observed and predicted pIC_50_ values for the training and test set compounds. Figure 2 demonstrates the external test set validation set prediction of *kpls_radial_15* hypothesis.

**Figure 1.**
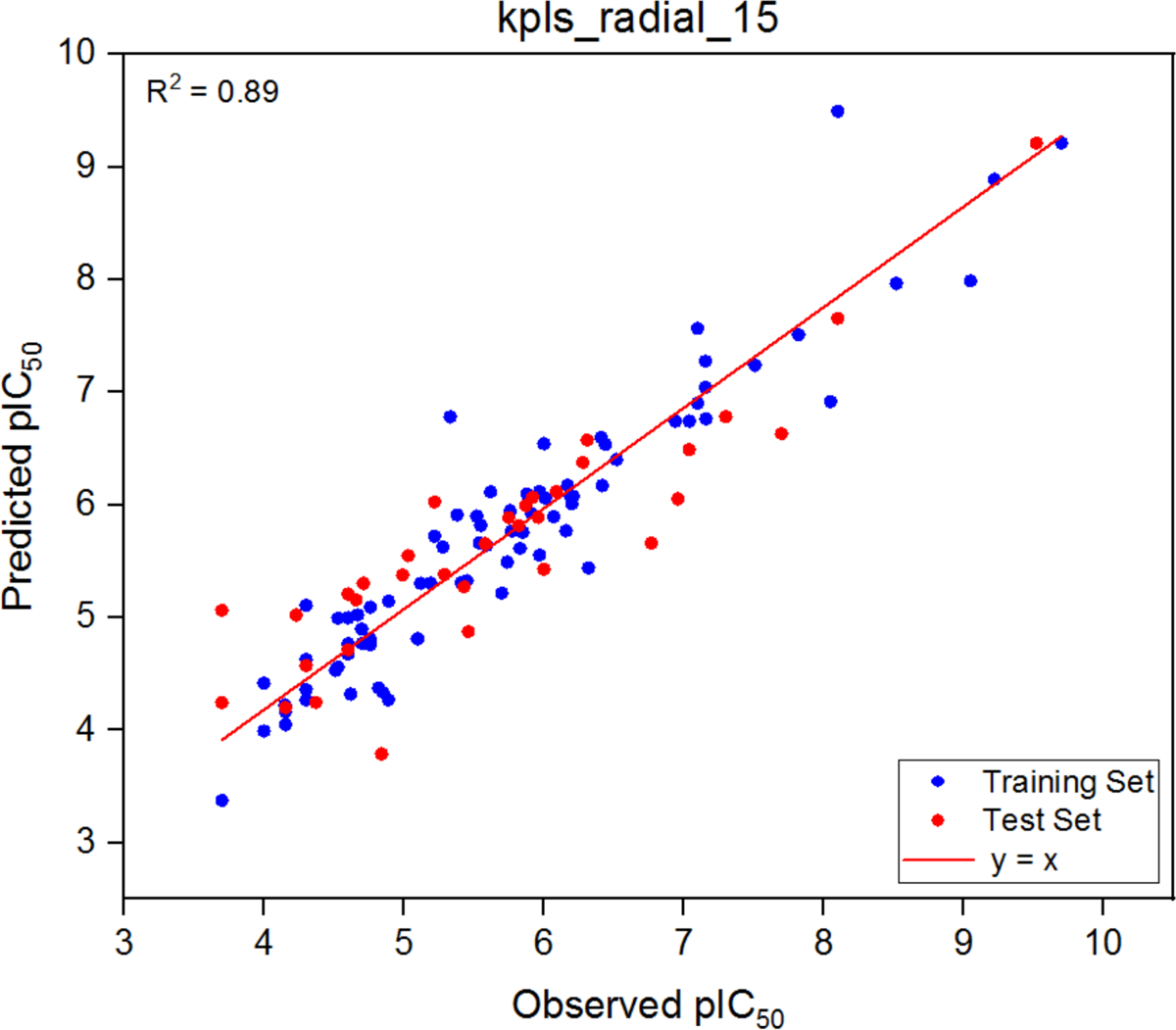
*kpls_radial_15* selected from U-87 cell line inhibitors and their predictive ability for the training and test set. R^2^=0.89; R=0.94.

**Figure 2.**
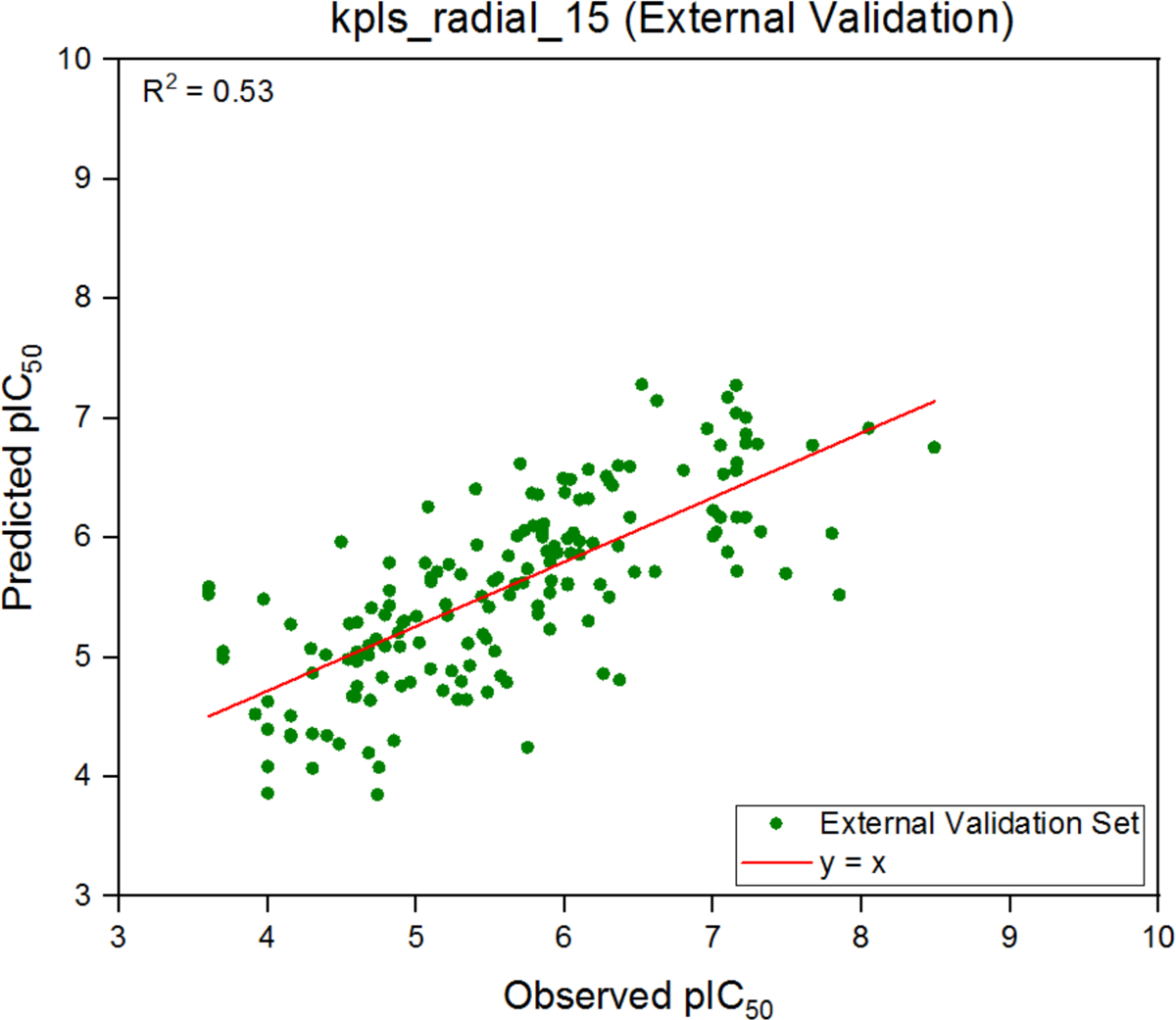
External validation of *kpls_radial_15* model that is constructed by U-87 cell line inhibitors. R^2^ = 0.53; R= 0.73.

**Table 1.** ML-based QSAR models generated from U-87 and U-251 cell line inhibitors.

| Model code | Score | S.D. | R<sup>2</sup> | RMSE | Q<sup>2</sup> |
| --- | --- | --- | --- | --- | --- |
| *kpls_radial_15* (U-87 cell line) | 0.72 | 0.42 | 0.89 | 0.56 | 0.80 |
| *kpls_molprint2D_36* (U-251 cell line) | 0.68 | 0.45 | 0.90 | 0.56 | 0.72 |

Similarly, another model was constructed using inhibitors against the U-251 cell line. 144 inhibitors were used in model generation and validation. Of those 144 molecules, 88 molecules were selected for the model generation, and 56 compounds were used as an external test set to validate the predictive power of the model set. The 88 molecules were then subdivided into 69.3% (61 compounds) as a training set and 30.7% (27 compounds) as a test set. (Tables S4 and S5) 56 molecules were used as external test set compounds which test the predictive power of the constructed model. These molecules at the external test did not participate in the model formation and were different from the training and test set. *kpls_molprint2D_36* model was selected as the best model (Table 1). Observed and predicted pIC_50_ values of training and test set are plotted in Figure 3. Figure 4 demonstrates the external validation set prediction of *kpls_molprint2D_36.* The remaining models constructed using U-87 and U-251 cell line data are detailed in Supplementary Tables S6 and S7, respectively.

**Figure 3.**
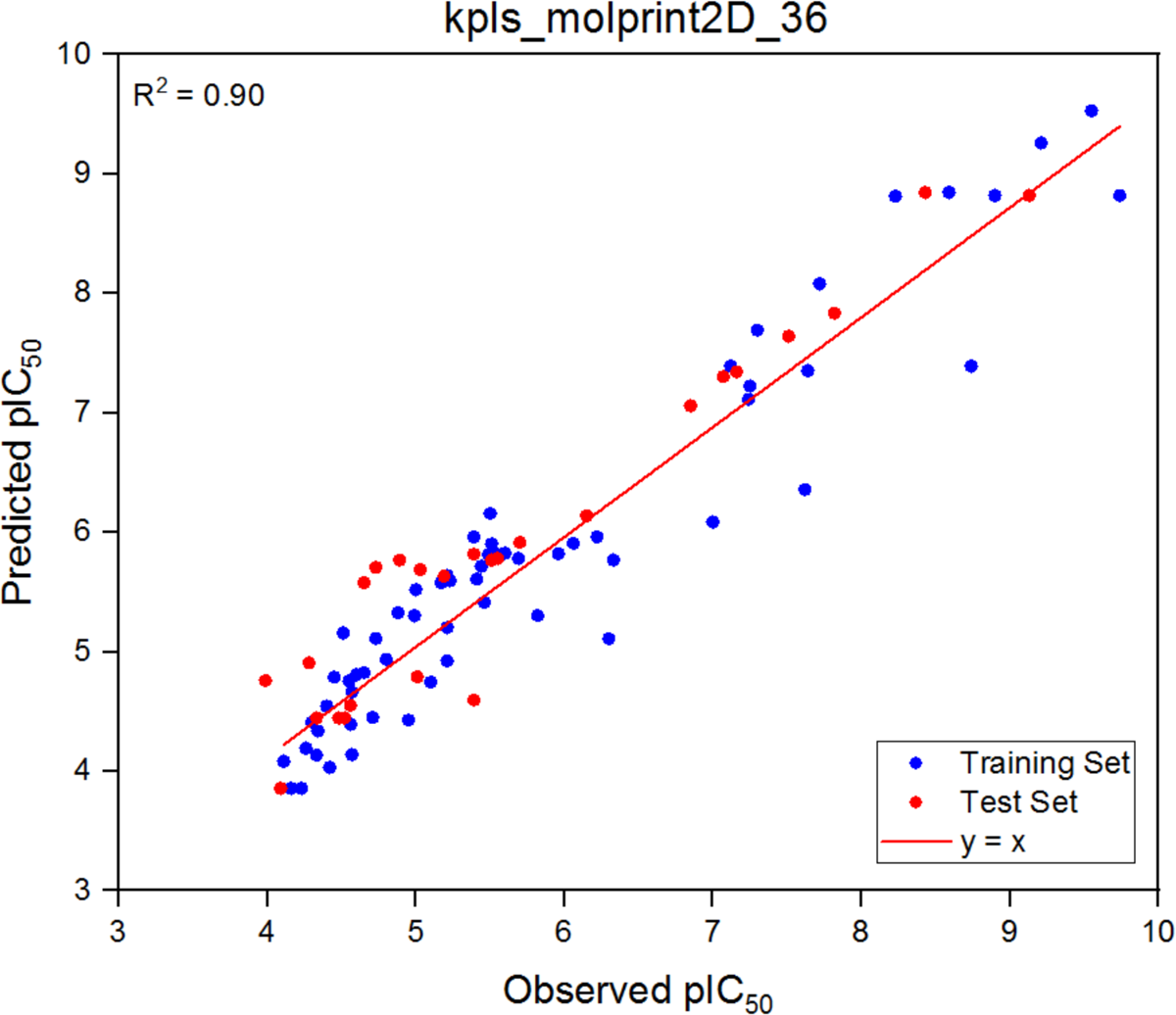
*kpls_molprint2D_36* model and its predictive ability for the training and test set of U- 251 cell line inhibitors. R^2^=0.90; R=0.95.

**Figure 4.**
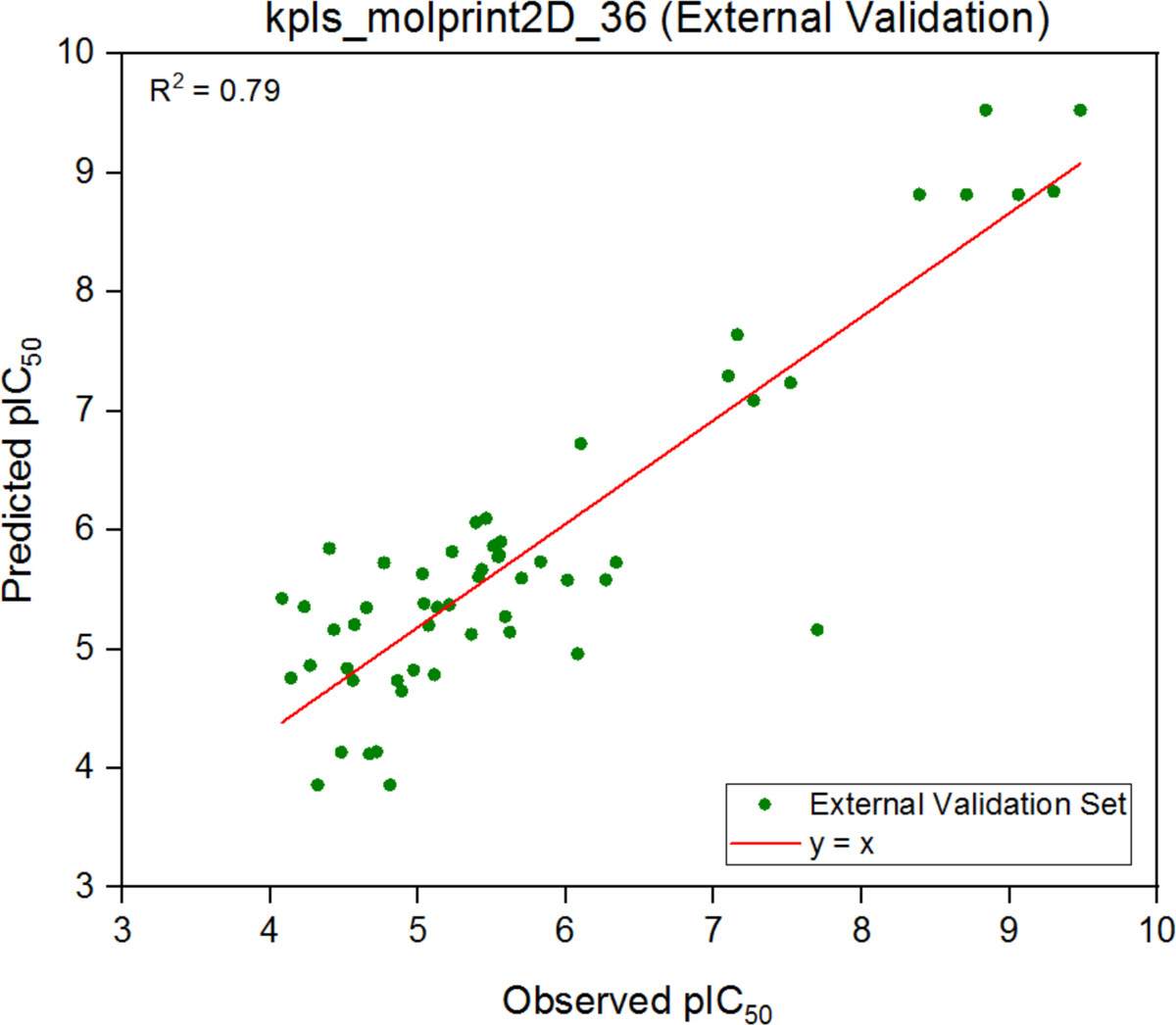
External validation for *kpls_molprint2D_36* model that is constructed by U-251 cell line inhibitors. R^2^=0.79; R=0.89.

### Construction of ML-based models using IDH1 mutant inhibitors

IDH1 mutant (R132H) inhibitors were downloaded from ChEMBL database with IC_50_ values and were used for ML-based model generation. The IC_50_ values were then converted to pIC_50_ values. Initially 197 IDH1 R132H mutant inhibitors were downloaded from ChEMBL and were used for model construction. Model was optimized by removing some outliers and in the end 173 compounds contributed to the learning set. Another set were constructed with 70% training set (122 compounds) and 30% test set (51 compounds) (Table S8). To validate the predictive ability of the models generated, 85 IDH1 mutant inhibitors from ChEMBL were used as external test, those molecules did not participate in the model formation and are different from the training and test set (Table S9). 10 models were generated from this model set and the best model (*kpls_desc_15*) was selected based on the predictive power with the external test set. Figure 5 shows observed and predicted pIC_50_ for the training and test set. Figure 6 demonstrates the external validation set prediction of the model. Table 2 shows the statistical significance of model including R^2^ and Q^2^.

**Figure 5.**
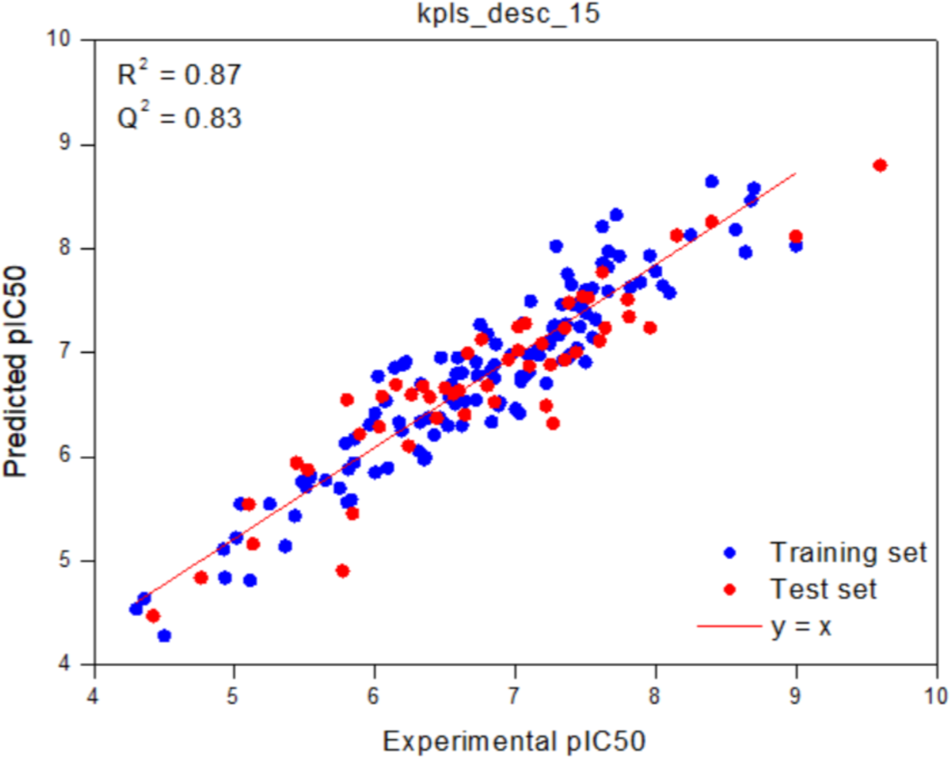
Experimental and predicted pIC_50_ values of learning set (training and test set) using *kpls desc_15* model. R^2^=0.87; R=0.93.

**Figure 6.**
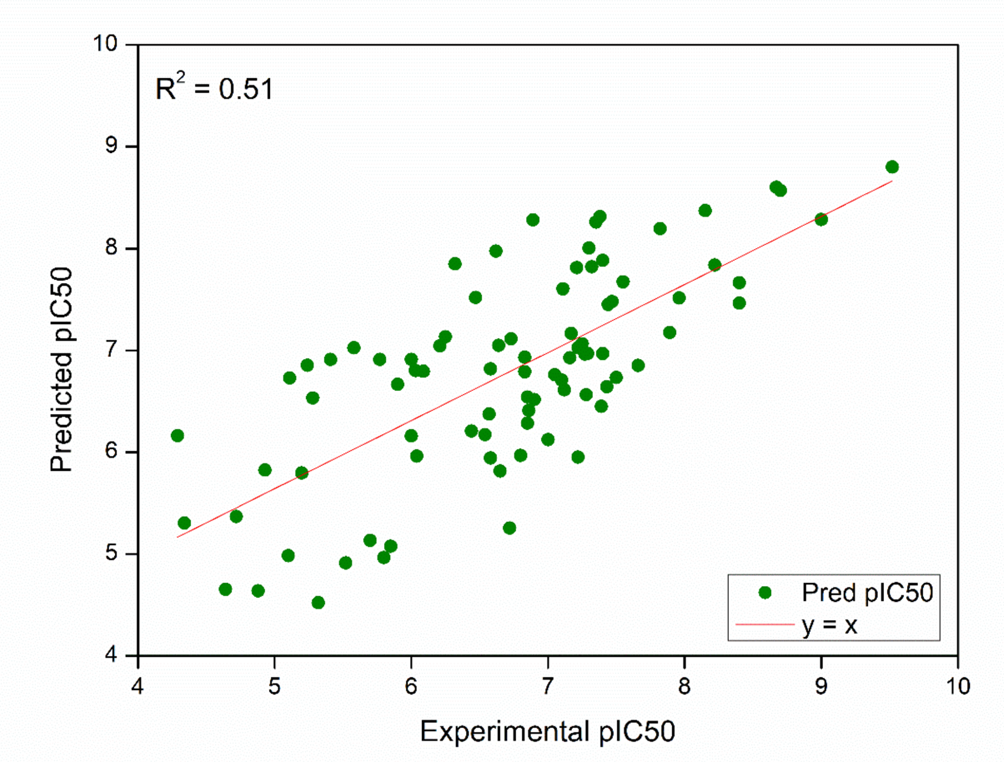
Experimental and predicted pIC_50_ values of external test set using *kpls desc_15* model. R^2^=0.51; R=0.71.

**Table 2.**
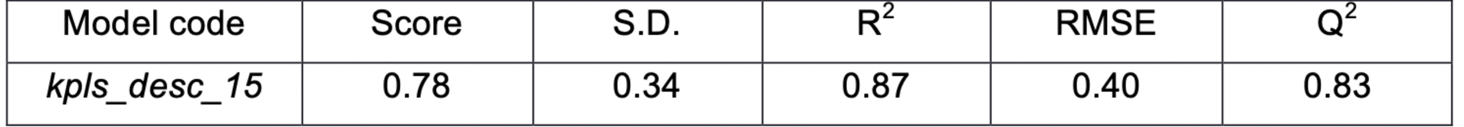
ML-based models generated from IDH1 mutant inhibitors.

| Model code | Score | S.D. | $R^2$ | RMSE | $Q^2$ |
| --- | --- | --- | --- | --- | --- |
| *kpls_desc_15* | 0.78 | 0.34 | 0.87 | 0.40 | 0.83 |

### Ligand-based pharmacophore (3D QSAR) analysis

For 3D QSAR ligand-based pharmacophore generation, U-87 cell line inhibitors were used to develop different 3D QSAR pharmacophore hypotheses, but all the hypotheses were statistically insignificant and were discarded. The dataset of inhibitors against the U-87 cell line used for this process is provided in Supplementary Table S10. Similarly, U-251 glioblastoma cell line inhibitors were downloaded from ChEMBL and were prepared using LigPrep, and conformers were generated using MacroModel. For each ligand, up to 1000 conformers were generated. The dataset was randomly divided into 67% training set and 33% internal test set (Dixon et al., 2006). The dataset for the U-251 cell line used for pharmacophore hypothesis and 3D QSAR model generation are available in Supplementary Table S11. The conformers were used in PHASE to develop common pharmacophore hypotheses module, and 4, 5, 6, 7 sited hypotheses were generated. The activity threshold was set as pIC_50_ ≤ 4.5 for inactive and pIC_50_ ≥ 7 for actives. 3 PLS factors were used, and pharmacophore-based hypotheses were generated. Two hypotheses (6-sited ADHRRR and 5-sited AAAHH) aligned well with the active compounds, so they were chosen as the statistically highest models. To validate generated 3D QSAR models (U-87 cell line data), another dataset of 56 compounds (i.e., external test set) was chosen with known experimental activities.

The inhibitors against the U-251 cell line used as an external test set for the generated PHASE model are detailed in Supplementary Table S12. Table 3 summarizes the obtained statistical results of the 6-sited and 5-sited models. Figures 7 and 8 show the prediction power of the generated models with the training and test set. Figures 9 and 10 demonstrate the external test set prediction ability of ADHRRR and AAAHH models. Figures 11 and 12 illustrate the resultant ADHRRR and AAAHH hypotheses, respectively, along with the best fitting active compounds from the training set. Regarding the construction of models using IDH1 mutant inhibitors, the models generated did not yield satisfactory prediction results for the external test set compounds. Consequently, these models are not included in further analysis.

**Figure 7.**
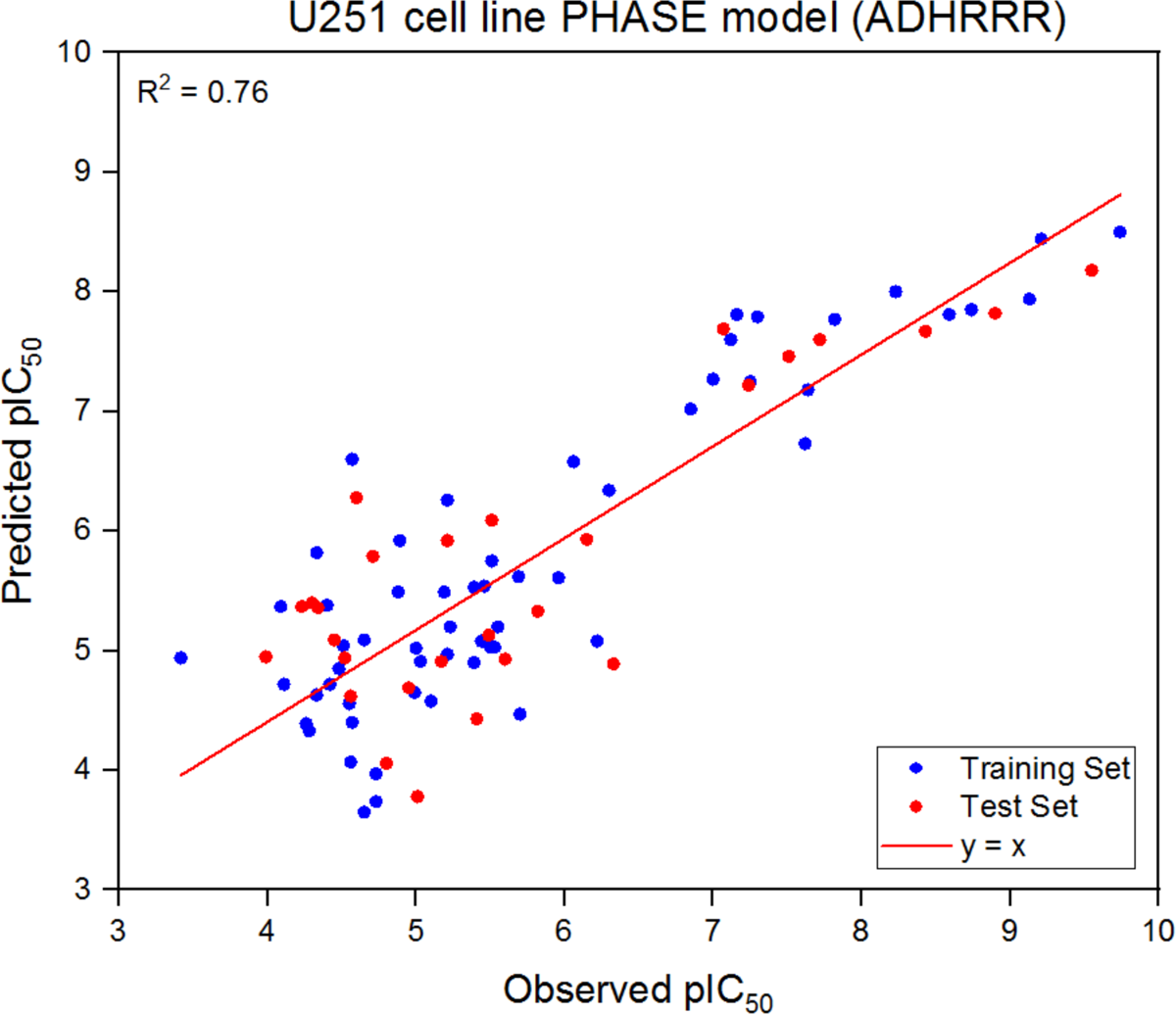
Experimental and predicted pIC_50_ values of 6 sited ADHRRR hypothesis for the training and test set of U-251 cell line inhibitors. R^2^=0.76; R=0.87.

**Figure 8.**
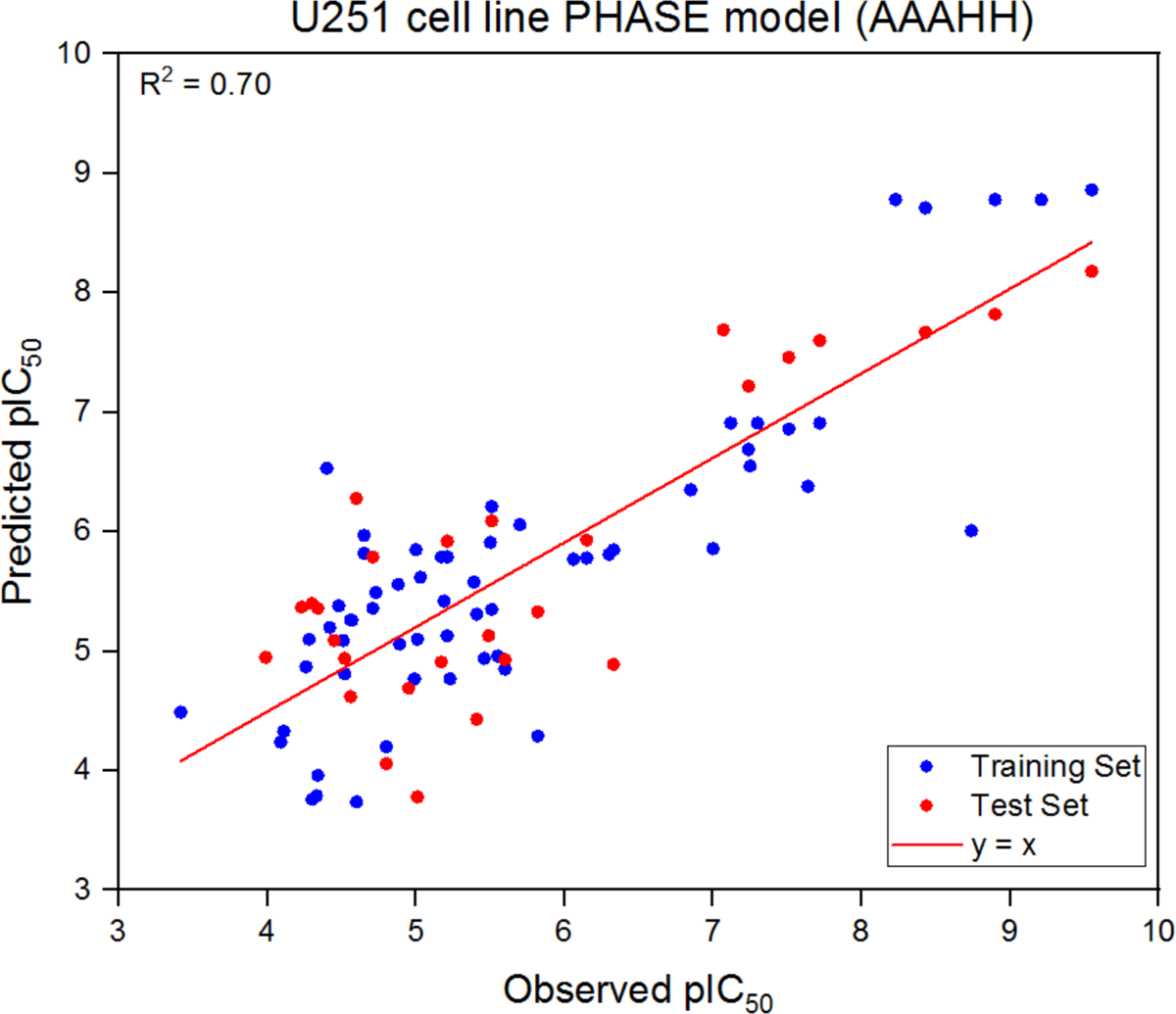
Experimental and predicted pIC_50_ values of 5 sited AAAHH hypothesis for the training and test set of U-251 cell line inhibitors. R^2^=0.70; R=0.84.

**Figure 9.**
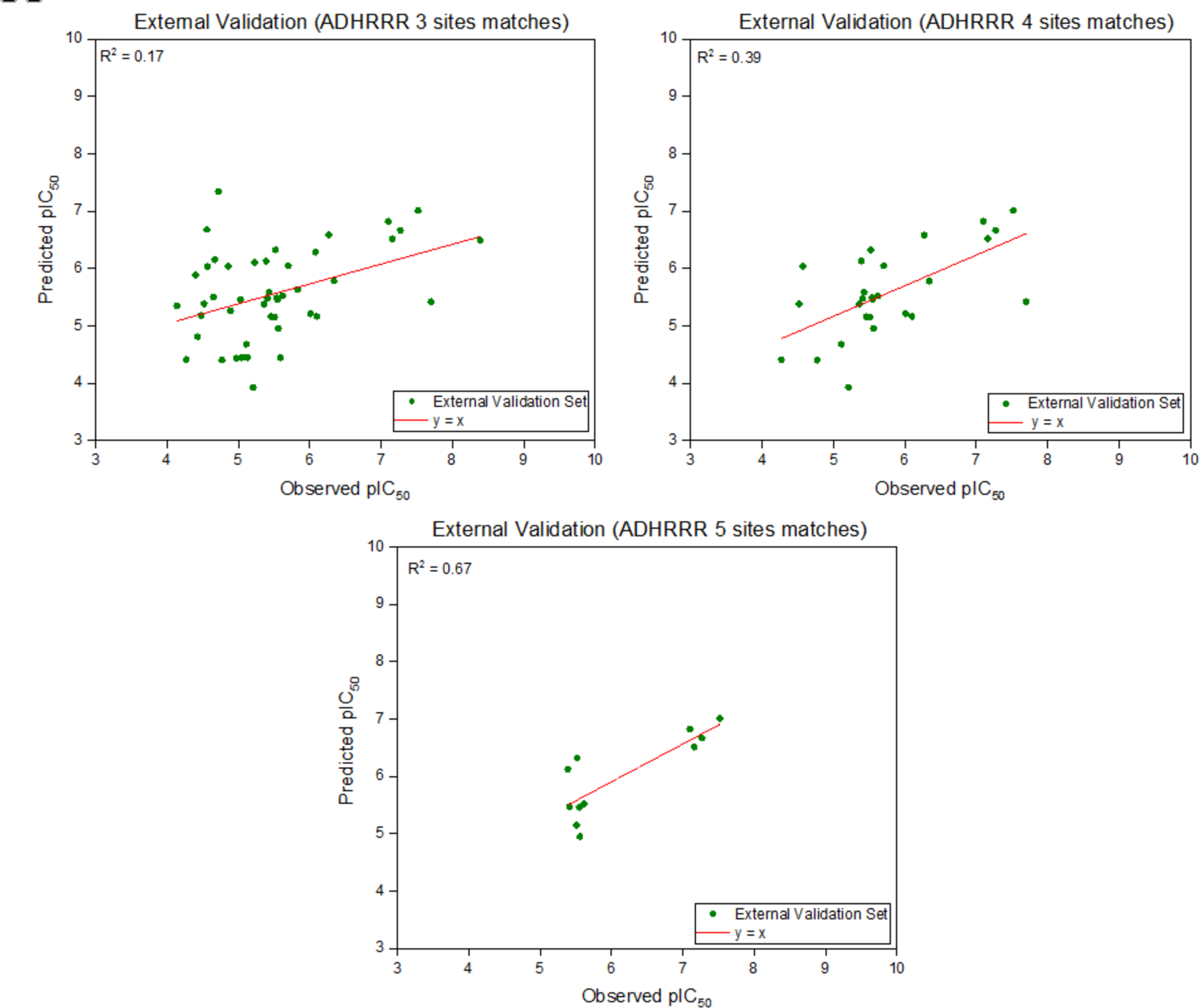
Validation of ADHRRR model using external test set (experimental and predicted pIC_50_ values).

**Figure 10.**
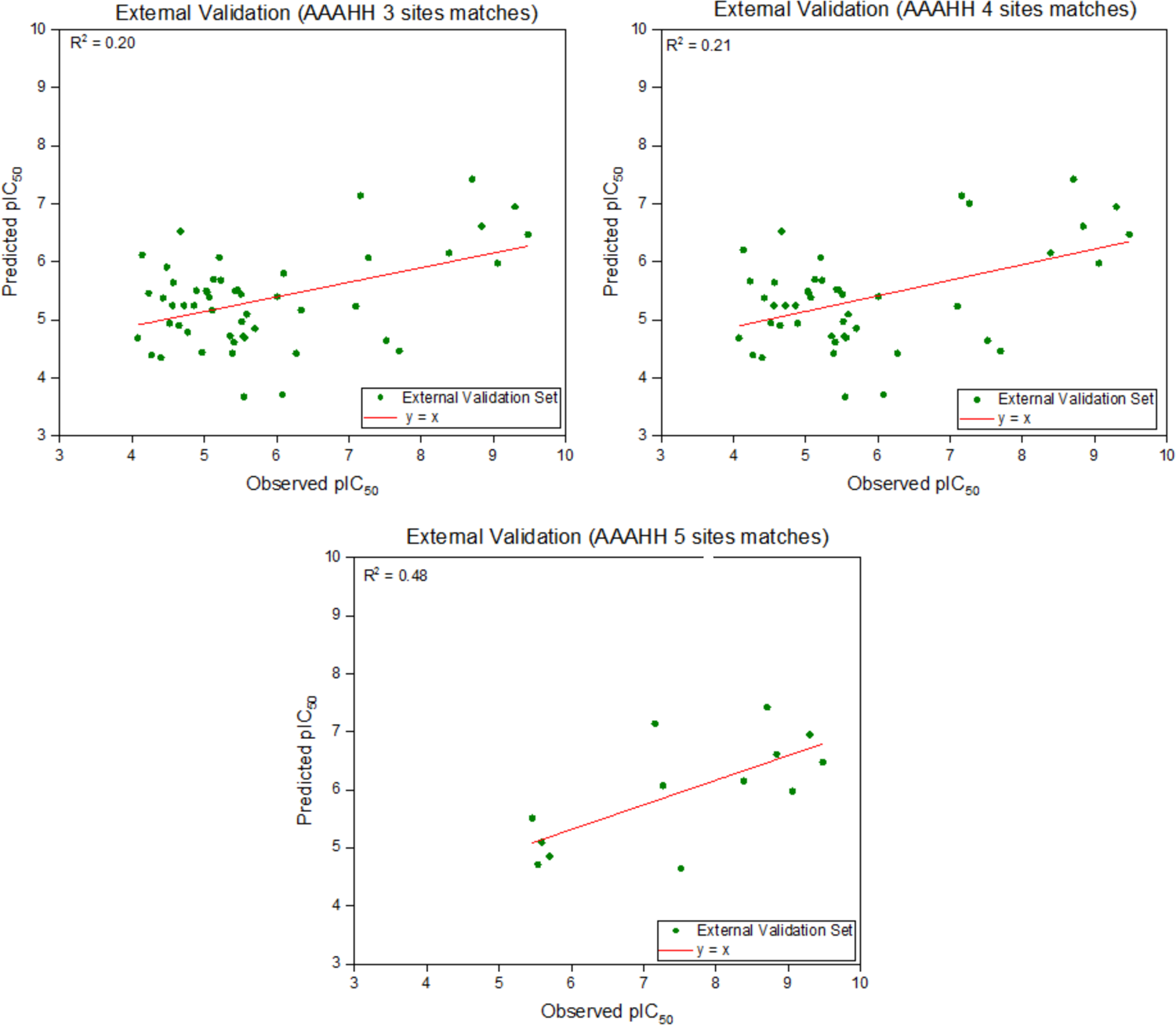
Validation of AAAHH model using external test set (experimental and predicted pIC_50_ values).

**Figure 11.**
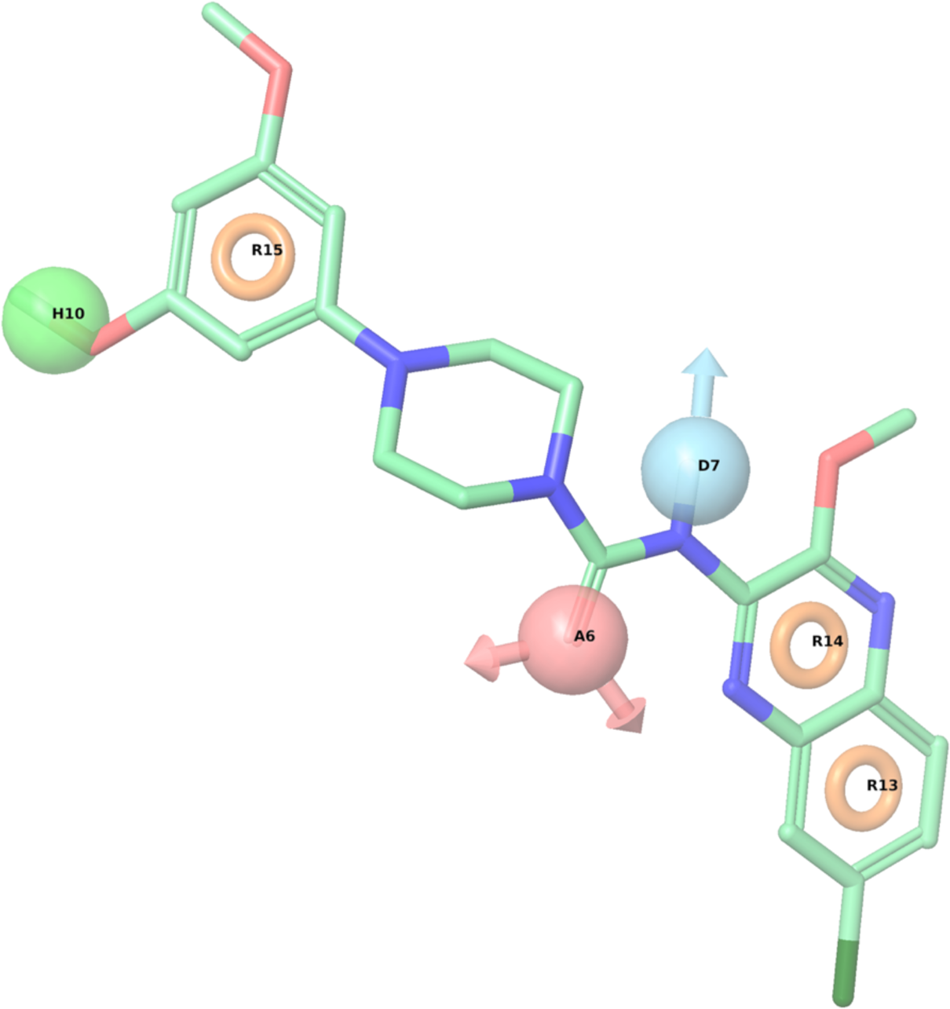
6 sited ADHRRR pharmacophore model (U-251 cell line inhibitors) aligned with a best active compound from the training set.

**Figure 12.**
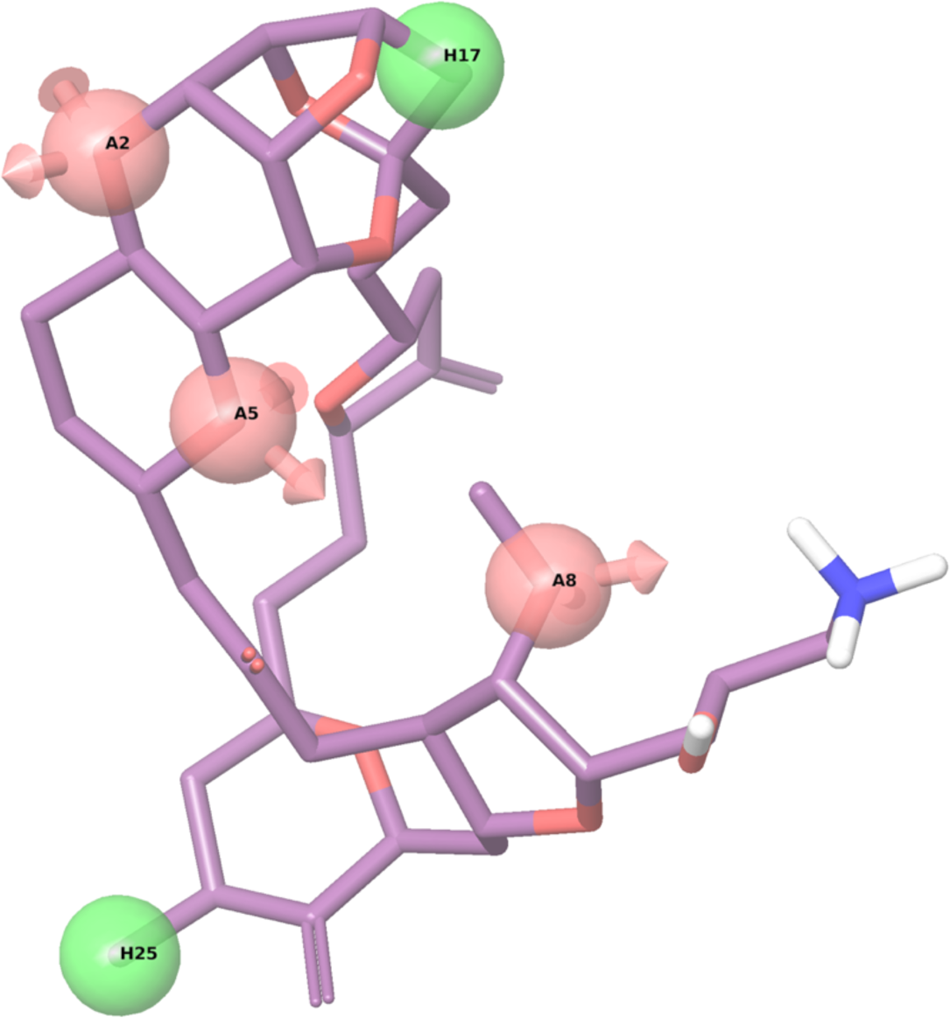
5 sited AAAHH pharmacophore model (U-251 cell line inhibitors) aligned with a best active compound from the training set.

**Table 3.** Ligand-based pharmacophore models generated from glioblastoma U-251 cell line inhibitors.

| Sites | Hypothesis | R<sup>2</sup> | Q<sup>2</sup> | Stability |
| --- | --- | --- | --- | --- |
| 6 | ADHRRR.612 | 0.76 | 0.73 | 0.72 |
| 5 | AAAHH.2487825 | 0.70 | 0.72 | 0.94 |

### Guner Henry Scoring Method

QSAR models were also checked based on the Guner Henry scoring method. The ability of the models to distinguish between active and inactives were validated using an external test set with observed pIC_50_ values. Guner Henry score was calculated for validation, and according to the reported criteria, scores > 0.7 indicates a statistically significant model. Some significant parameters such as total number of compounds in database (D), % of known actives in database (A), active hits (*Ha*), total hits (*Ht*), and goodness-of-hit score (*GH*) were calculated (Table 4). It was observed to be 0.97 for the *kpls_molprint2D_36*, an AutoQSAR based model. The GH score shows the efficiency of the QSAR models. (Vyas et al., 2012) For the *kpls_radial_15* model, corresponding result was 0.57. The GH score for the PHASE model (ADHRRR) was calculated as 0.75, and (AAAHH) was calculated as 0.8, indicating these models are reliable for virtual screening. However, in contrast to the ML-based models, external test set statistical results show that the ADHRRR and AAAHH hypotheses did not predict a high correlation between observed and predicted values.

**Table 4.** GH score of the selected models.

| Approach Name | Model Name | Total No in Database (D) | % of Known Actives (A) | Active Hits (Ha) | Total Hits (Ht) | GH score |
| --- | --- | --- | --- | --- | --- | --- |
| ML-based models | *kpls Molprint2D 36* | 56 | 11 | 10 | 10 | 0.97 |
|  | *kpls radial 15* | 170 | 28 | 5 | 7 | 0.57 |
| PHASE (3D-QSAR) | ADHRRR | 11 | 4 | 1 | 1 | 0.75 |
|  | AAAHH | 47 | 11 | 3 | 3 | 0.80 |

### Structure-based pharmacophore (e-Pharmacophore) model generation

e-Pharmacophore hypotheses were generated using 9 IDH1 mutant complexes from protein databank (6B0Z, 6ADG, 5SUN, 5TQH, 5LGE, 5SVF, 5L57, 5L58, 4UMX). (Wang et al., 2020) Generated hypotheses aligned with their reference ligands are reported in Table 5.

**Table 5.**
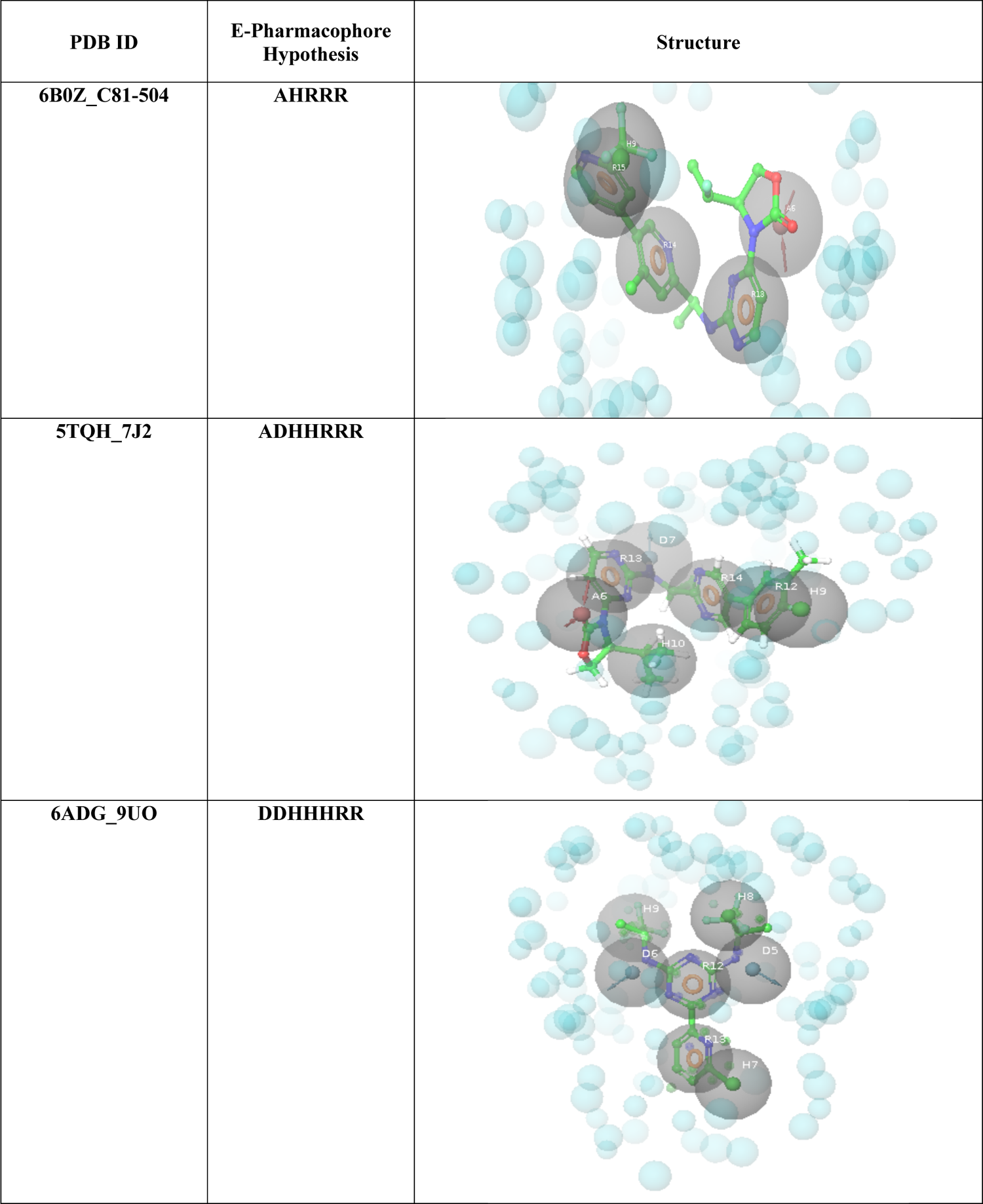
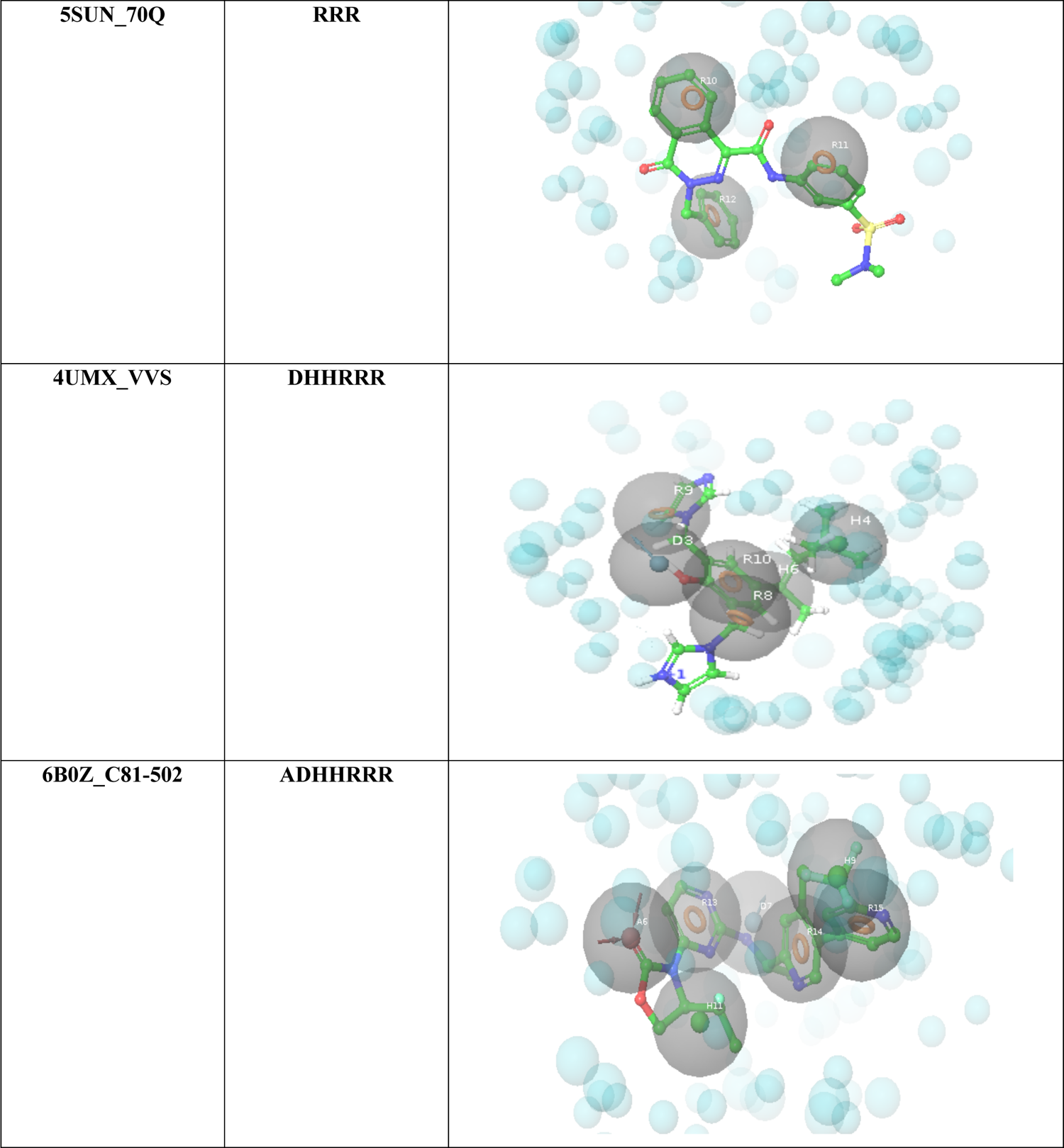
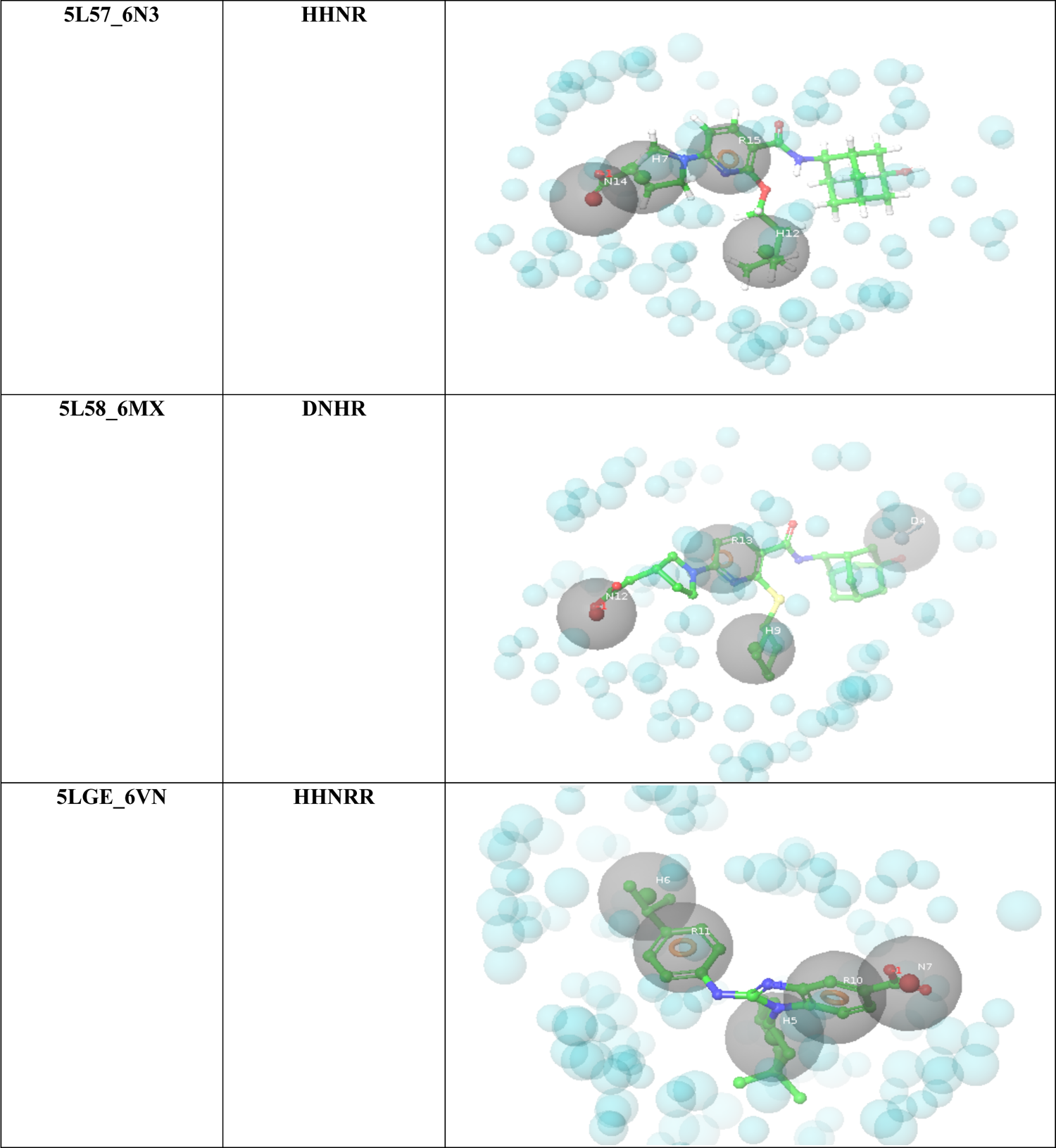
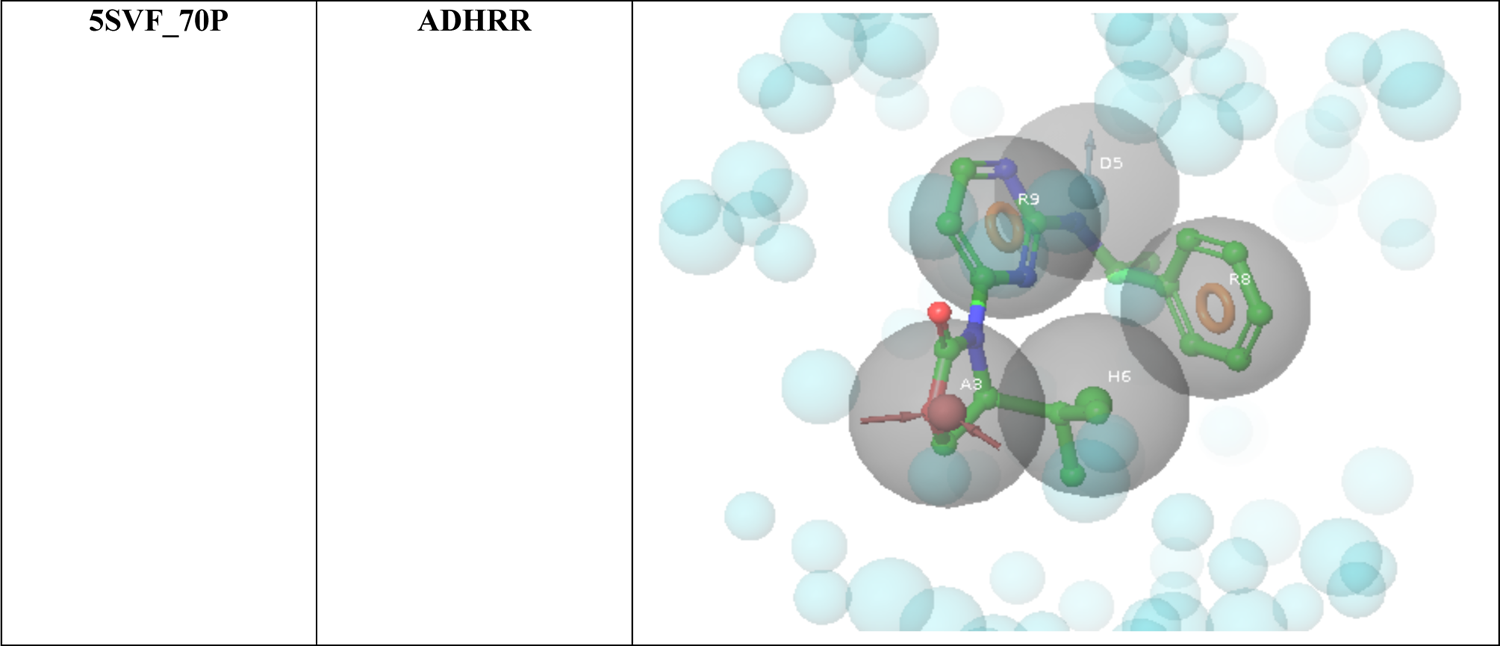
e-Pharmacophore hypotheses from 9 PDB targets and their reference ligands fitted on hypotheses.

## Conclusions

Glioblastoma is a very complex brain tumor, and the efficiency of the current therapeutic approaches are still low. This invokes the necessity to search for alternative approaches in finding and identifying novel drugs that have good efficacy and can slow down the progression and invasiveness of this disease. In this study, diverse approaches were used to develop models that have good predictive ability and selectivity for possible hits against GBM. The ligand-based models generated using both cell-line inhibitors and IDH1 mutant inhibitors were tested internally (during model generation) and validated externally, then the GH score for each ligand-based model was calculated. The ML-based constructed models showed good predictive ability, statistical significance, and accuracy, that makes them a promising tool for the virtual screening of diverse small molecule libraries. These ML-based QSAR models are ready to screen extensive small molecule libraries against GBM.

