## supporting information for "Development of Machine Learning-based QSAR Models for the Designing of Novel Anti-cancer Therapeutics Against Malignant Glioma"

[1]Equal contribution.

**Table S1:** Construction of AutoQSAR U87 using machine learning-based algorithm (116 compounds training and test set from U87 cell line) along with $pIC_{50}$.

| Sr. No | Compound ID | 2D structure | $pIC_{50}$ | Dataset Division (Train & Test) |
|---|---|---|---|---|
| 1. | CHEMBL589346 | 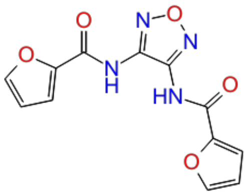 | 3.70 | test |
| 2. | CHEMBL591046 | 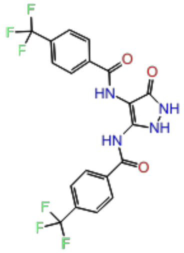 | 3.70 | train |
| 3. | CHEMBL592492 | 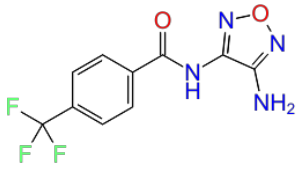 | 3.70 | test |
| 4. | CHEMBL331760 | 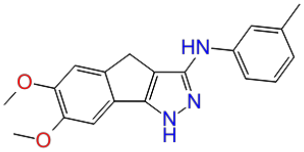 | 4.00 | train |
| 5. | CHEMBL592219 | 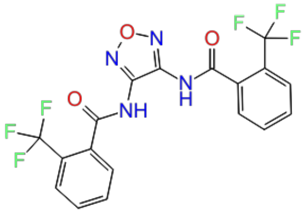 | 4.00 | train |

| | | | | |
|---|---|---|---|---|
| 6. | CHEMBL602439 | 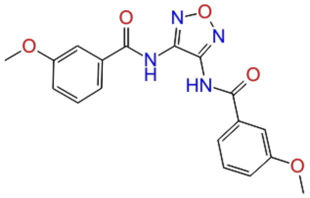 | 4.15 | train |
| 7. | CHEMBL213022 | 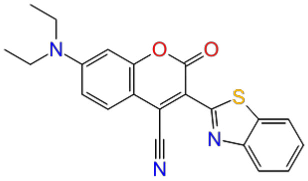 | 4.15 | test |
| 8. | CHEMBL213488 | 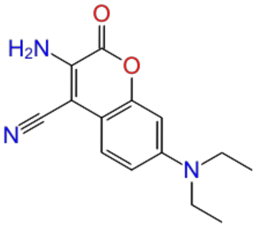 | 4.15 | train |
| 9. | CHEMBL214947 | 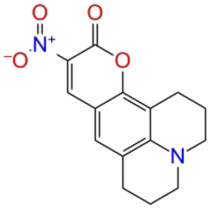 | 4.15 | train |
| 10. | CHEMBL378647 | 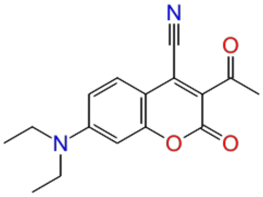 | 4.15 | train |
| 11. | CHEMBL4291595 | 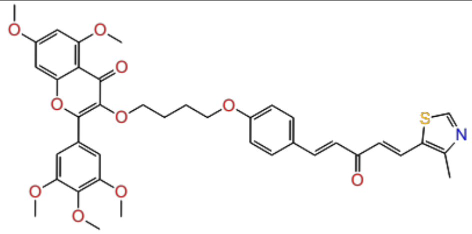 | 4.23 | test |

| | | | | |
|---|---|---|---|---|
| 12. | CHEMBL121161 | 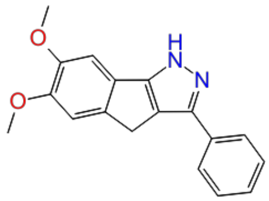 | 4.30 | train |
| 13. | CHEMBL4226663 | 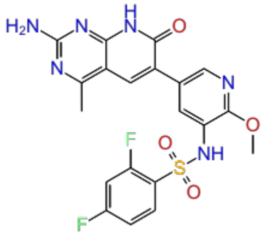 | 4.30 | train |
| 14. | CHEMBL4291032 | 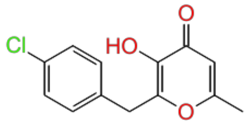 | 4.30 | train |
| 15. | CHEMBL4293250 | 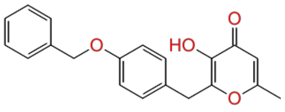 | 4.30 | train |
| 16. | CHEMBL4294325 | 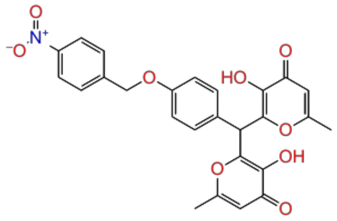 | 4.30 | test |
| 17. | CHEMBL213746 | 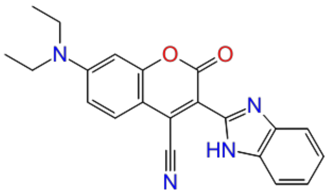 | 4.37 | test |
| 18. | CHEMBL4289867 | 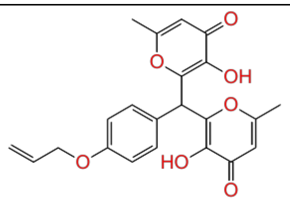 | 4.51 | train |

| | | | | |
|---|---|---|---|---|
| 19. | CHEMBL3588944 | 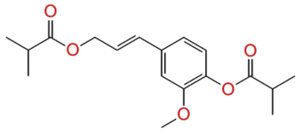 | 4.53 | train |
| 20. | CHEMBL4279183 | 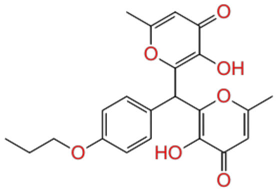 | 4.53 | train |
| 21. | CHEMBL4077217 | 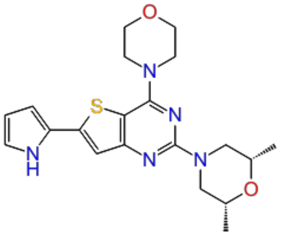 | 4.60 | train |
| 22. | CHEMBL4080009 | 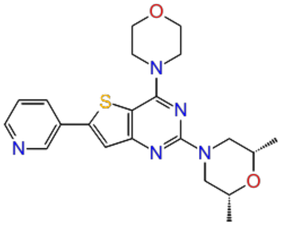 | 4.60 | test |
| 23. | CHEMBL4082222 | 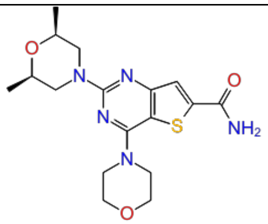 | 4.60 | train |
| 24. | CHEMBL4084746 | 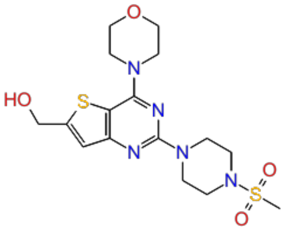 | 4.60 | test |
| 25. | CHEMBL4101802 | 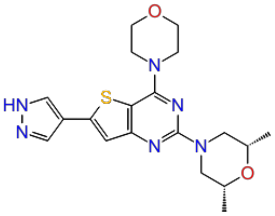 | 4.60 | train |

| | | | | |
|---|---|---|---|---|
| 26. | CHEMBL589334 | 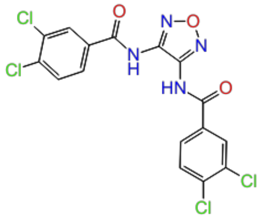 | 4.62 | train |
| 27. | CHEMBL4291163 | 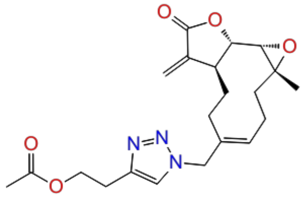 | 4.66 | test |
| 28. | CHEMBL4063923 | 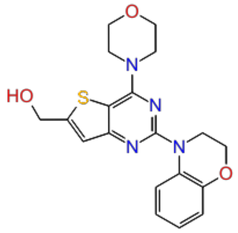 | 4.67 | train |
| 29. | CHEMBL4283154 | 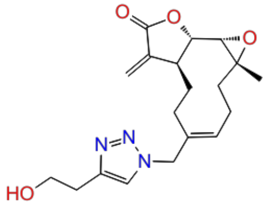 | 4.70 | train |
| 30. | CHEMBL4205181 | 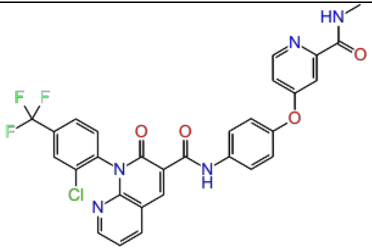 | 4.70 | train |
| 31. | CHEMBL4288767 | 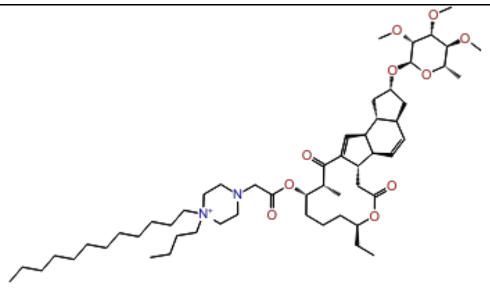 | 4.71 | test |

| | | | | |
|---|---|---|---|---|
| 32. | CHEMBL465843 | 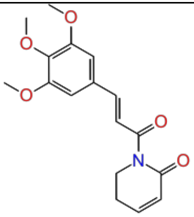 | 4.76 | train |
| 33. | CHEMBL3588945 | 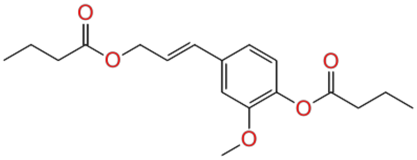 | 4.76 | train |
| 34. | CHEMBL4088600 | 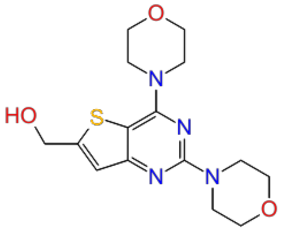 | 4.76 | train |
| 35. | CHEMBL374632 | 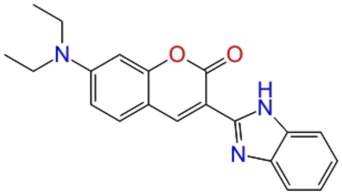 | 4.82 | train |
| 36. | CHEMBL600476 | 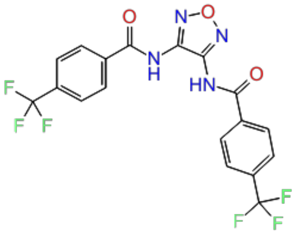 | 4.84 | test |
| 37. | CHEMBL589654 | 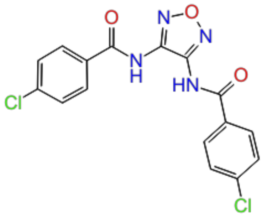 | 4.85 | train |

| | | | | |
|---|---|---|---|---|
| 38. | CHEMBL120077 | 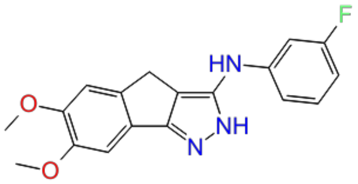 | 4.89 | train |
| 39. | CHEMBL4287584 | 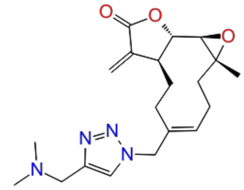 | 4.89 | train |
| 40. | CHEMBL4218367 | 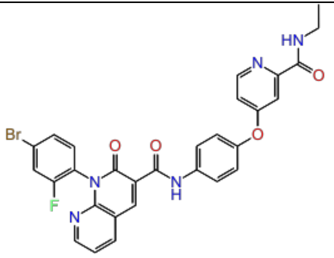 | 4.99 | test |
| 41. | CHEMBL4293071 | 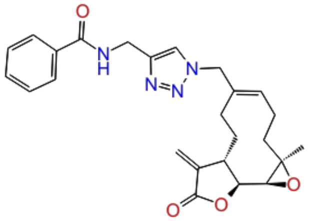 | 5.03 | test |
| 42. | CHEMBL591753 | 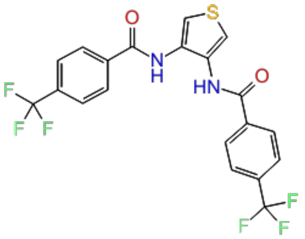 | 5.10 | train |
| 43. | CHEMBL4291994 | 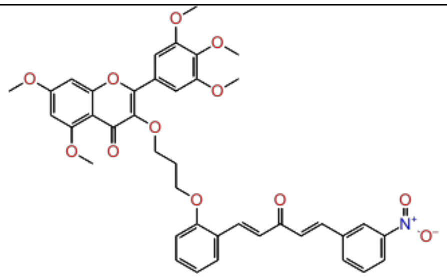 | 5.10 | train |

| | | | | |
|---|---|---|---|---|
| 44. | CHEMBL4285411 | 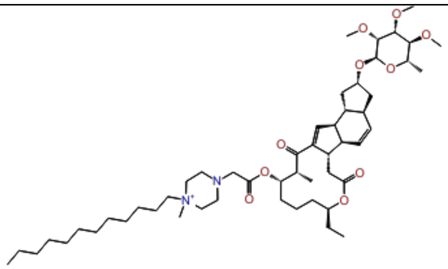 | 5.12 | train |
| 45. | CHEMBL4293675 | 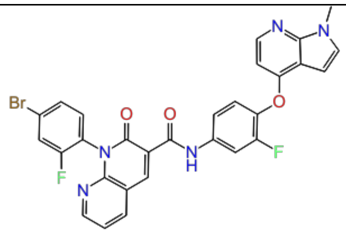 | 5.19 | train |
| 46. | CHEMBL119037 | 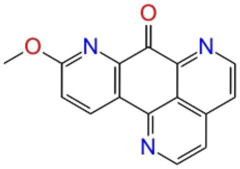 | 5.22 | train |
| 47. | CHEMBL119894 | 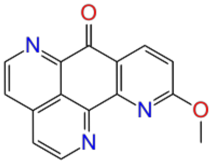 | 5.22 | test |
| 48. | CHEMBL4286245 | 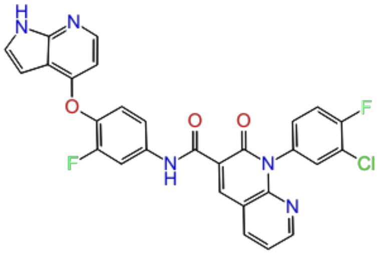 | 5.28 | train |
| 49. | CHEMBL4209066 | 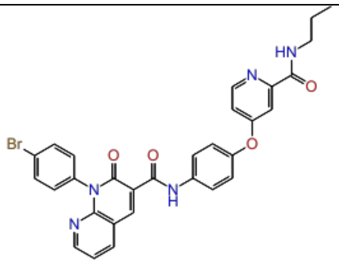 | 5.29 | test |

| | | | | |
|---|---|---|---|---|
| 50. | CHEMBL111845 | 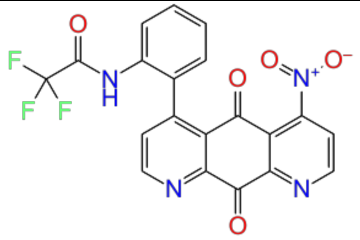 | 5.33 | train |
| 51. | CHEMBL4089188 | 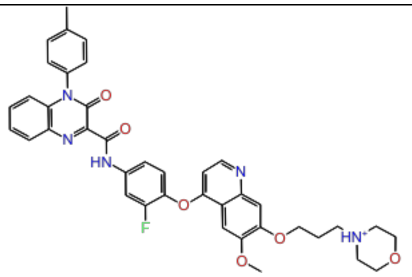 | 5.38 | train |
| 52. | CHEMBL4281382 | 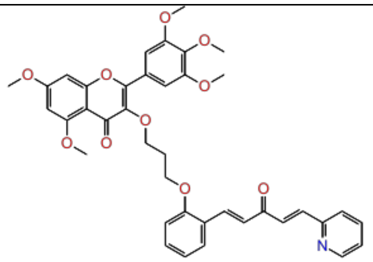 | 5.41 | train |
| 53. | CHEMBL4288699 | 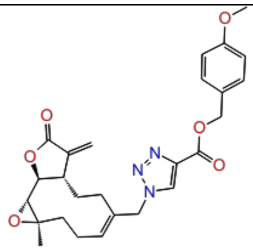 | 5.43 | test |
| 54. | CHEMBL4207763 | 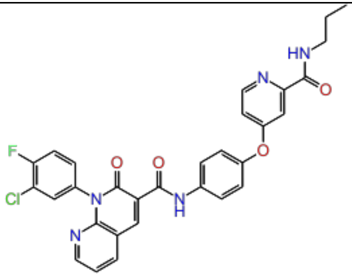 | 5.45 | train |

| | | | | |
|---|---|---|---|---|
| 55. | CHEMBL4062698 | 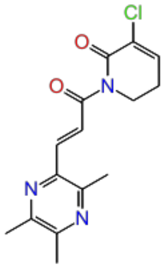 | 5.46 | test |
| 56. | CHEMBL119393 | 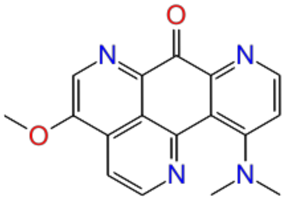 | 5.52 | train |
| 57. | CHEMBL4207018 | 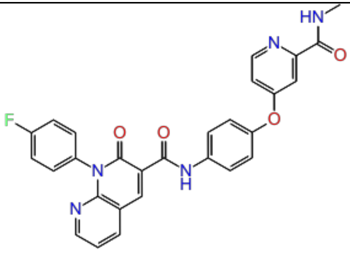 | 5.54 | train |
| 58. | CHEMBL4284206 | 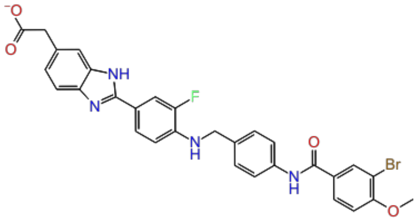 | 5.55 | train |
| 59. | CHEMBL4213745 | 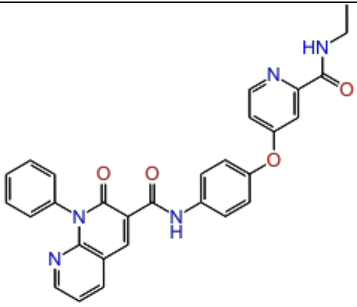 | 5.58 | test |

| | | | | |
|---|---|---|---|---|
| 60. | CHEMBL4286454 | 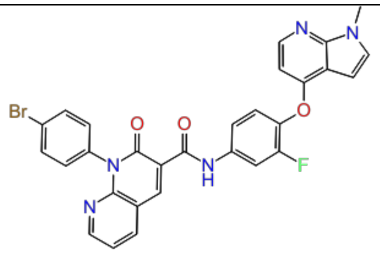 | 5.59 | train |
| 61. | CHEMBL3612198 | 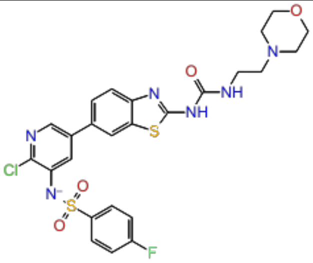 | 5.62 | train |
| 62. | CHEMBL4289953 | 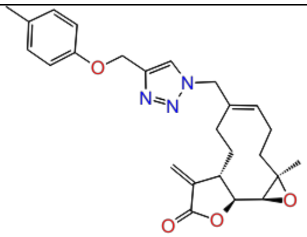 | 5.70 | train |
| 63. | CHEMBL352708 | 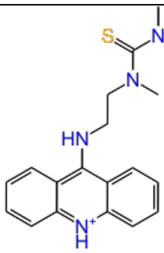 | 5.74 | train |
| 64. | CHEMBL4291079 | 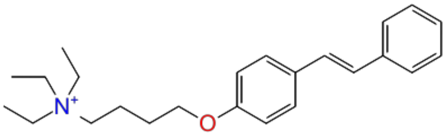 | 5.75 | test |
| 65. | CHEMBL3612196 | 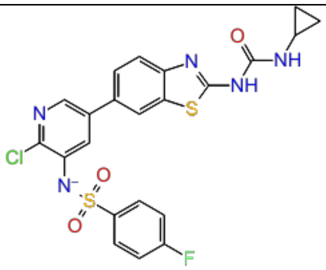 | 5.76 | train |

| | | | | |
|---|---|---|---|---|
| 66. | CHEMBL4080590 | 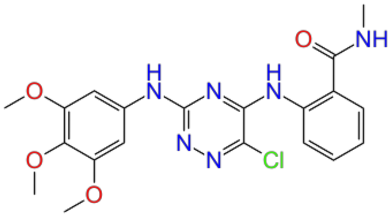 | 5.77 | train |
| 67. | CHEMBL4288088 | 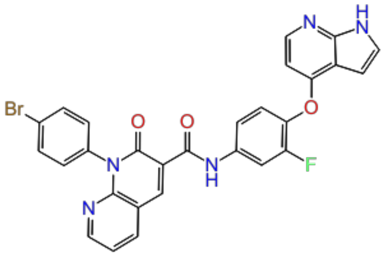 | 5.82 | test |
| 68. | CHEMBL4292276 | 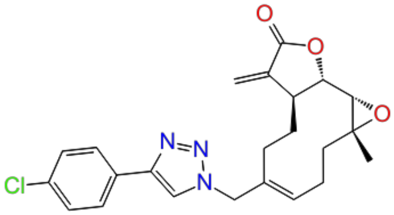 | 5.83 | train |
| 69. | CHEMBL513 | 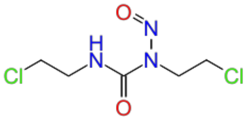 | 5.85 | train |
| 70. | CHEMBL1230609 | 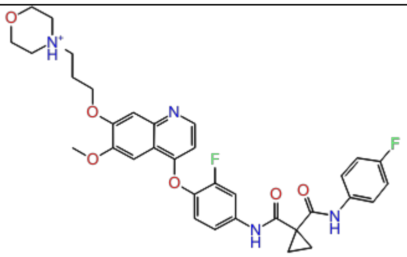 | 5.87 | test |
| 71. | CHEMBL1230609 | 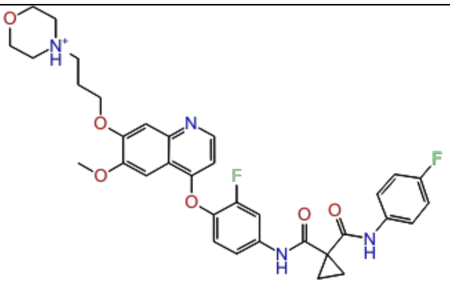 | 5.87 | train |

| | | | | |
|---|---|---|---|---|
| 72. | CHEMBL4081754 | 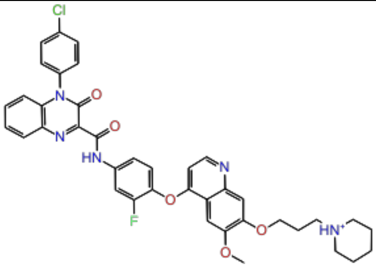 | 5.88 | train |
| 73. | CHEMBL4071208 | 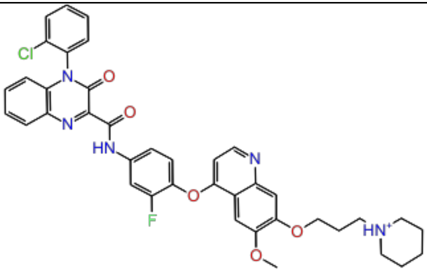 | 5.91 | train |
| 74. | CHEMBL4092999 | 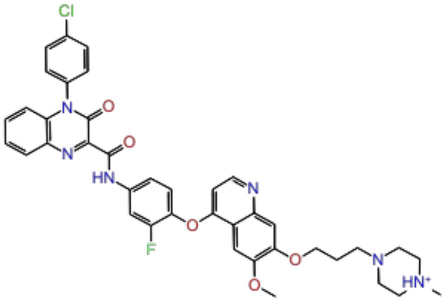 | 5.92 | test |
| 75. | CHEMBL4072236 | 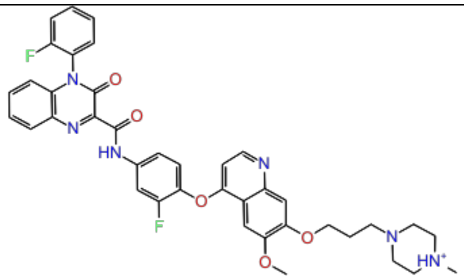 | 5.96 | test |
| 76. | CHEMBL3588939 | 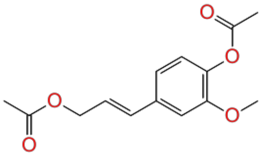 | 5.97 | train |

| | | | | |
|---|---|---|---|---|
| 77. | CHEMBL4085110 | 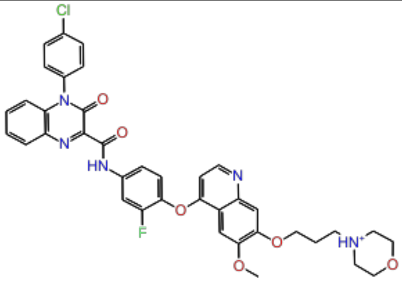 | 5.97 | train |
| 78. | CHEMBL4246923 | 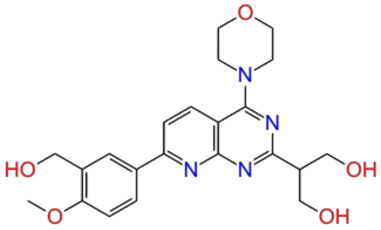 | 6.00 | train |
| 79. | CHEMBL4290751 | 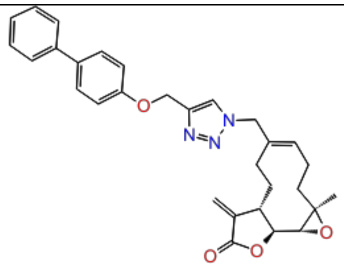 | 6.00 | test |
| 80. | CHEMBL4073794 | 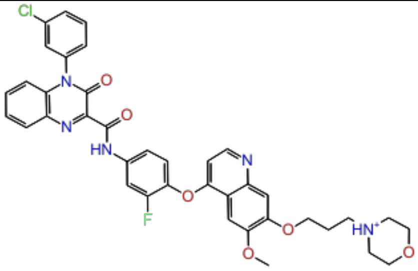 | 6.01 | train |
| 81. | CHEMBL4276776 | 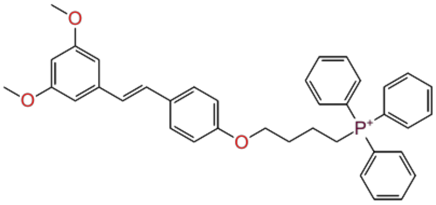 | 6.07 | train |

| | | | | |
|---|---|---|---|---|
| 82. | CHEMBL4095304 | 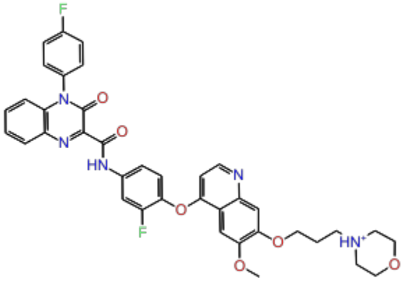 | 6.09 | test |
| 83. | CHEMBL334217 | 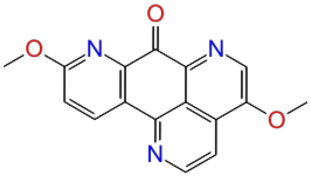 | 6.16 | train |
| 84. | CHEMBL4288506 | 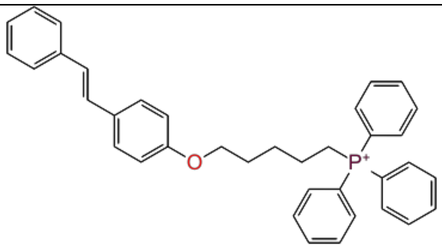 | 6.17 | train |
| 85. | CHEMBL4284717 | 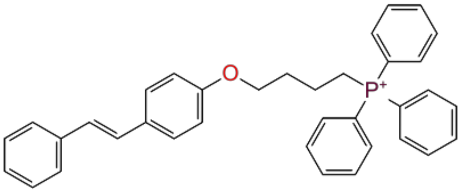 | 6.19 | train |
| 86. | CHEMBL3112718 | 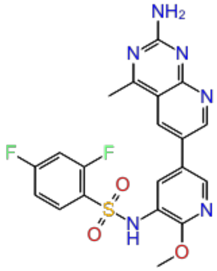 | 6.20 | train |
| 87. | CHEMBL4289577 | 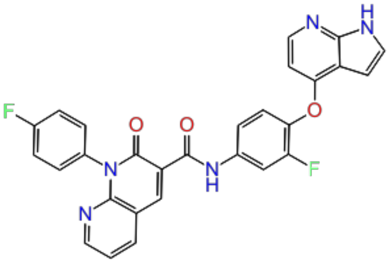 | 6.21 | train |

| | | | | |
|---|---|---|---|---|
| 88. | CHEMBL3612309 | 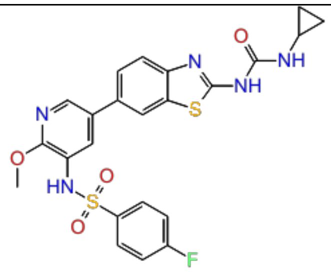 | 6.28 | test |
| 89. | CHEMBL3612311 | 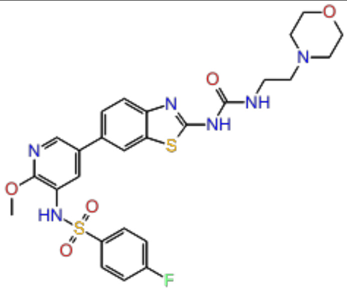 | 6.31 | test |
| 90. | CHEMBL3588937 | 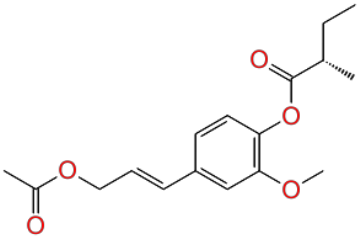 | 6.32 | train |
| 91. | CHEMBL3612310 | 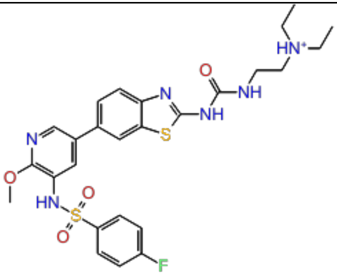 | 6.41 | train |
| 92. | CHEMBL4277852 | 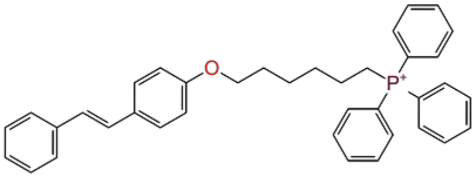 | 6.42 | train |

| | | | | |
|---|---|---|---|---|
| 93. | CHEMBL3612316 | 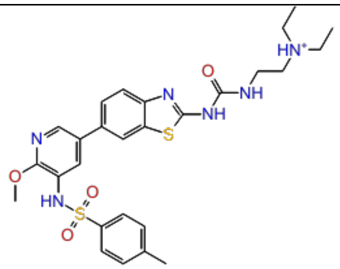 | 6.44 | train |
| 94. | CHEMBL4228946 | 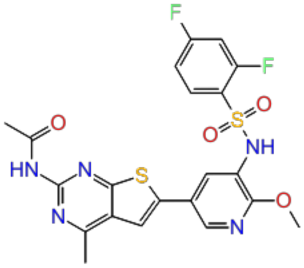 | 6.52 | train |
| 95. | CHEMBL458997 | 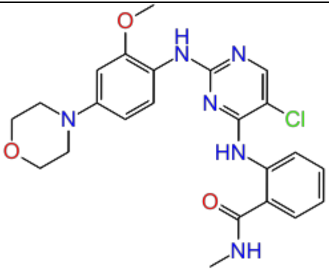 | 6.77 | test |
| 96. | CHEMBL4169540 | 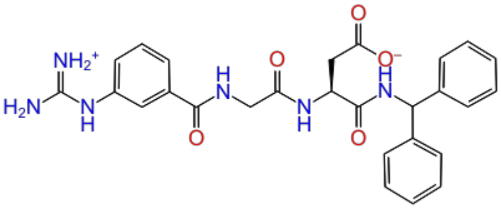 | 6.94 | train |
| 97. | CHEMBL4285772 | 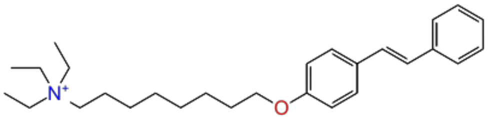 | 6.96 | test |

| | | | | |
|---|---|---|---|---|
| 98. | CHEMBL4240170 | 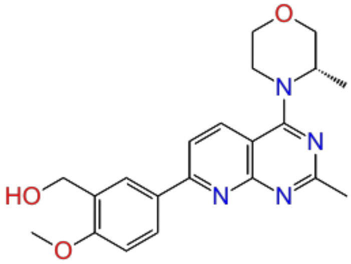 | 7.04 | train |
| 99. | CHEMBL4248529 | 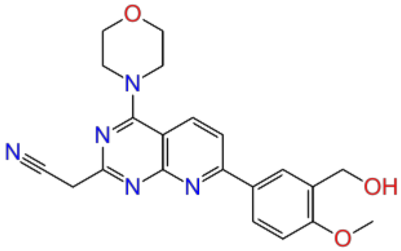 | 7.04 | test |
| 100. | CHEMBL424062 | 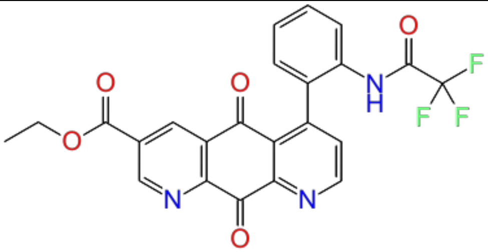 | 7.10 | train |
| 101. | CHEMBL428647 | 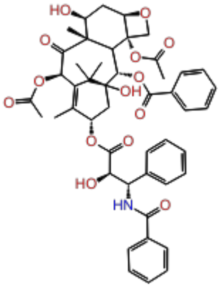 | 7.10 | train |
| 102. | CHEMBL125395 | 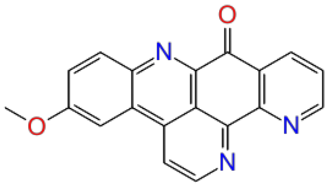 | 7.16 | train |

| | | | | |
|---|---|---|---|---|
| 103. | CHEMBL340160 | 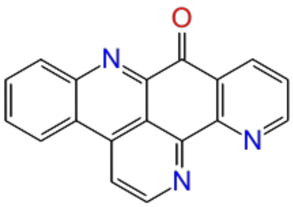 | 7.16 | train |
| 104. | CHEMBL322816 | 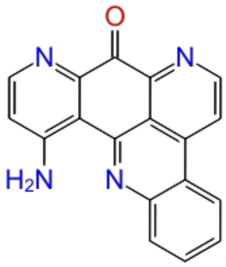 | 7.16 | train |
| 105. | CHEMBL109479 | 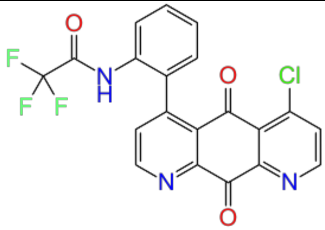 | 7.30 | test |
| 106. | CHEMBL252164 | 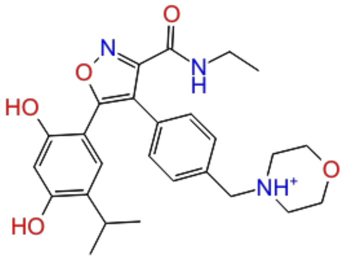 | 7.51 | train |
| 107. | CHEMBL113387 | 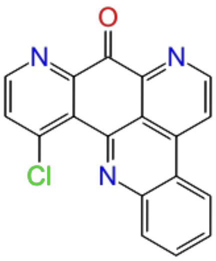 | 7.70 | test |

| | | | | |
|---|---|---|---|---|
| 108. | CHEMBL4277900 | 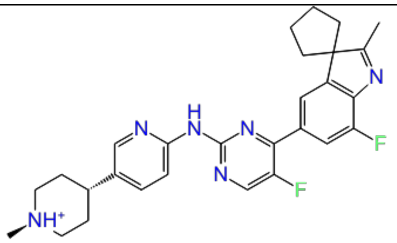 | 7.82 | train |
| 109. | CHEMBL111490 | 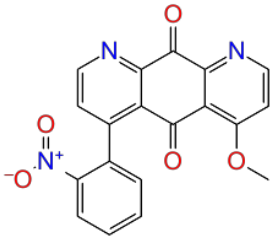 | 8.05 | train |
| 110. | CHEMBL4162270 | 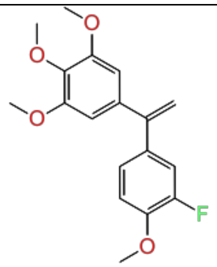 | 8.10 | test |
| 111. | CHEMBL498271 | 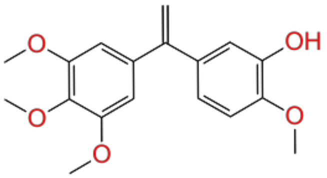 | 8.10 | train |
| 112. | CHEMBL67 | 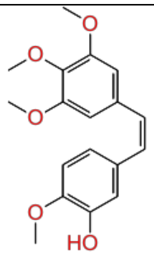 | 8.52 | train |
| 113. | CHEMBL110742 | 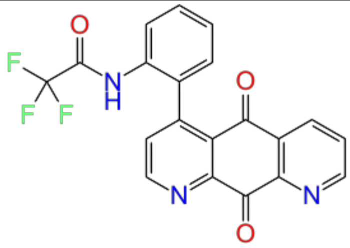 | 9.05 | train |

| | | | | |
|---|---|---|---|---|
| 114. | CHEMBL4172942 | 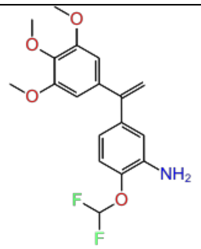 | 9.22 | train |
| 115. | CHEMBL508617 | 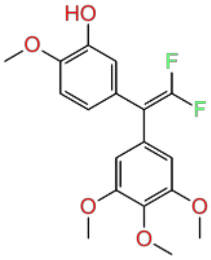 | 9.52 | test |
| 116. | CHEMBL4159373 | 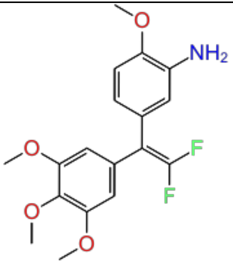 | 9.70 | train |

**Table S2:** AutoQSAR U87 internal validation set (56 compounds U87 cell line) along with $pIC_{50}$.

| Sr. No | Compound ID | 2D structure | $pIC_{50}$ |
|---|---|---|---|
| 1. | CHEMBL590092 | 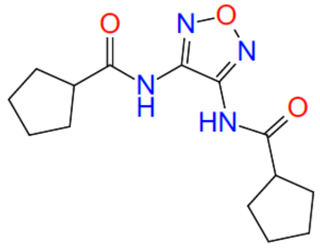 | 3.70 |
| 2. | CHEMBL120634 | 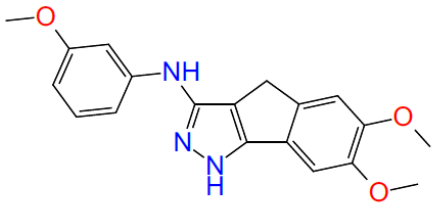 | 4.00 |
| 3. | CHEMBL4280797 | 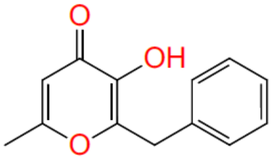 | 4.30 |
| 4. | CHEMBL602040 | 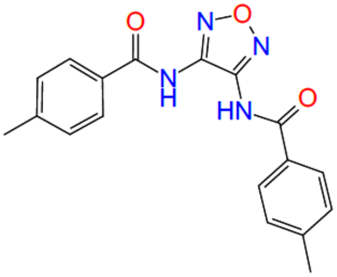 | 4.39 |
| 5. | CHEMBL4295010 | 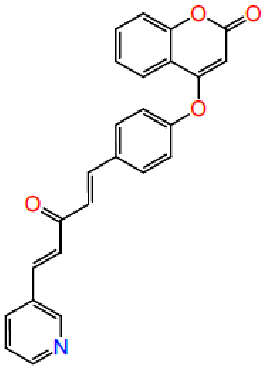 | 4.43 |

| | | | |
|---|---|---|---|
| 6. | CHEMBL4063088 | 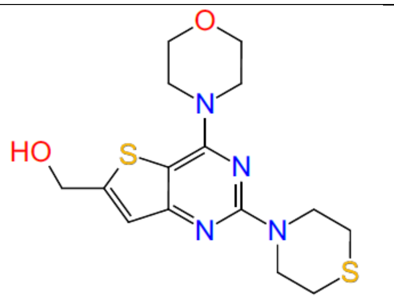 | 4.60 |
| 7. | CHEMBL200735 | 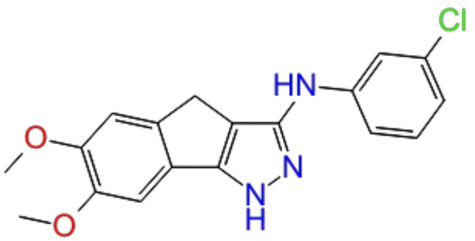 | 4.62 |
| 8. | CHEMBL589123 | 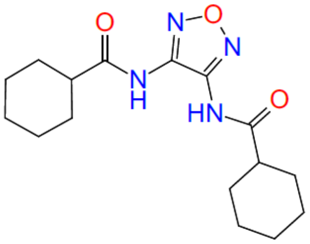 | 4.70 |
| 9. | CHEMBL2180727 | 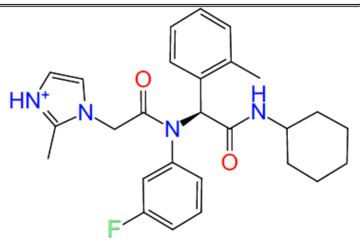 | 4.70 |
| 10. | CHEMBL4088848 | 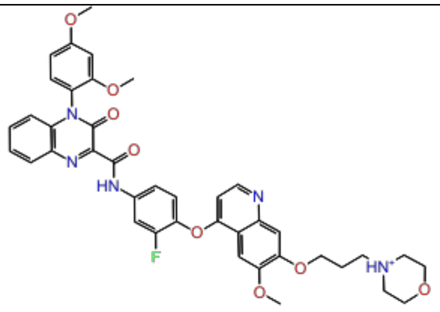 | 4.79 |

| | | | |
|---|---|---|---|
| 11. | CHEMBL4286248 | 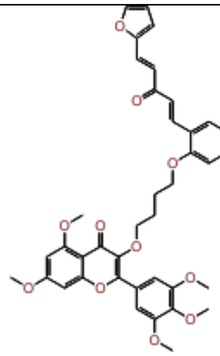 | 4.97 |
| 12. | CHEMBL3588942 | 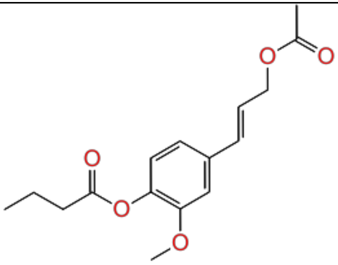 | 4.98 |
| 13. | CHEMBL4279607 | 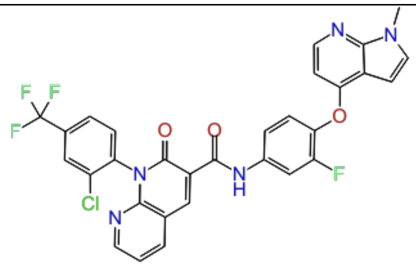 | 4.99 |
| 14. | CHEMBL4205819 | 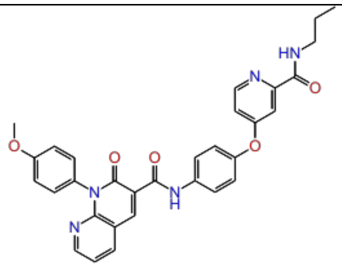 | 5.09 |
| 15. | CHEMBL4100831 | 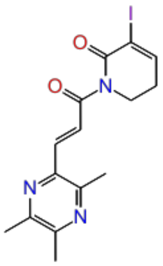 | 5.14 |

| | | | |
|---|---|---|---|
| 16. | CHEMBL4226543 | 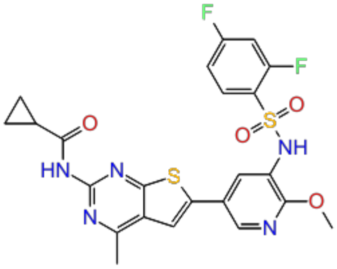 | 5.19 |
| 17. | CHEMBL4286146 | 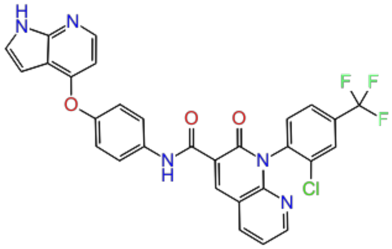 | 5.22 |
| 18. | CHEMBL4278984 | 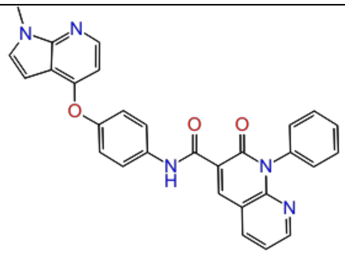 | 5.30 |
| 19. | CHEMBL1257065 | 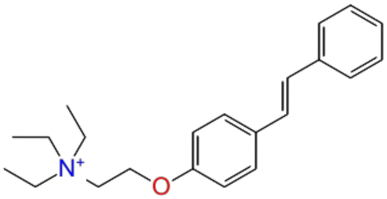 | 5.36 |
| 20. | CHEMBL4090663 | 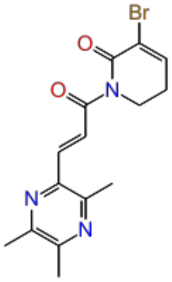 | 5.37 |

| | | | |
|---|---|---|---|
| 21. | CHEMBL4287907 | 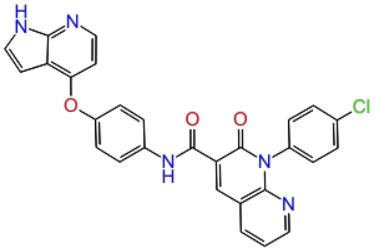 | 5.47 |
| 22. | CHEMBL233930 | 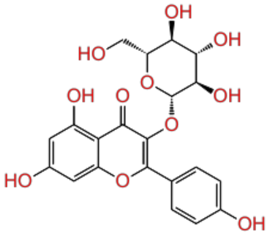 | 5.51 |
| 23. | CHEMBL4281371 | 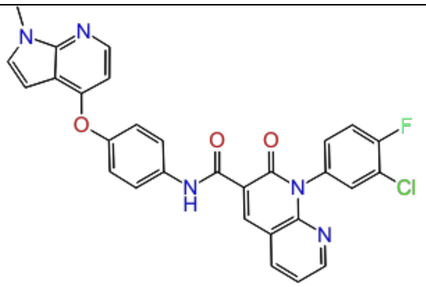 | 5.63 |
| 24. | CHEMBL4225608 | 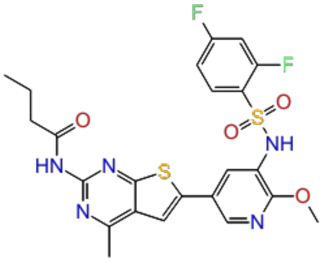 | 5.64 |
| 25. | CHEMBL4100714 | 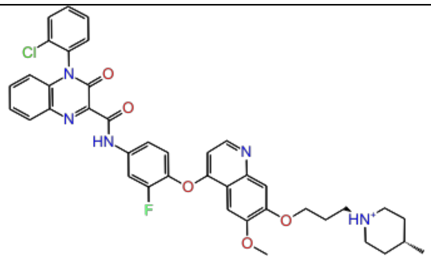 | 5.66 |
| 26. | CHEMBL331163 | 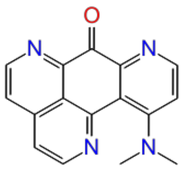 | 5.70 |

| | | | |
|---|---|---|---|
| 27. | CHEMBL4282597 | 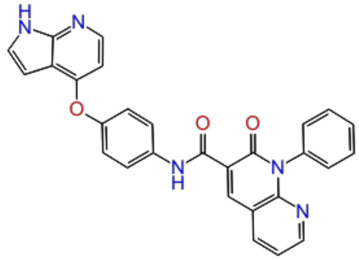 | 5.80 |
| 28. | CHEMBL4099282 | 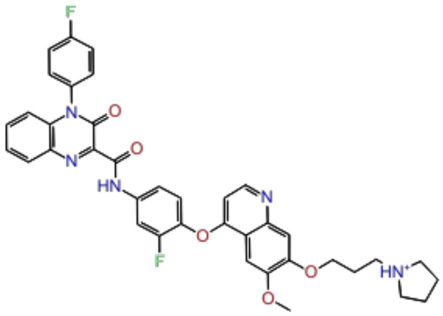 | 5.82 |
| 29. | CHEMBL4280528 | 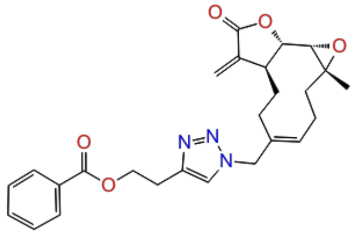 | 5.82 |
| 30. | CHEMBL4278178 | 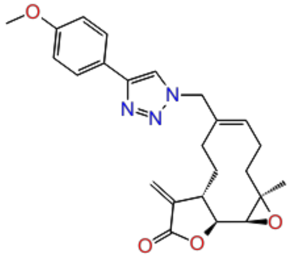 | 5.84 |
| 31. | CHEMBL3612199 | 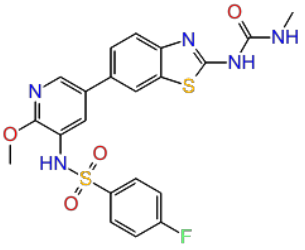 | 5.91 |

| | | | |
|---|---|---|---|
| 32. | CHEMBL3612320 | 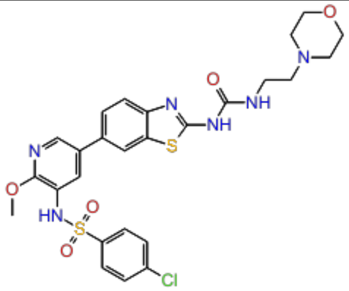 | 5.98 |
| 33. | CHEMBL4063742 | 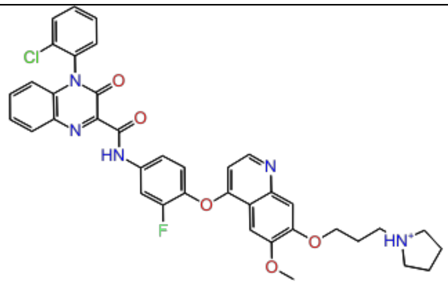 | 5.99 |
| 34. | CHEMBL601898 | 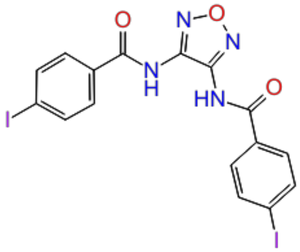 | 6.00 |
| 35. | CHEMBL53463 | 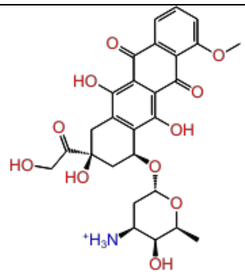 | 6.03 |
| 36. | CHEMBL4072335 | 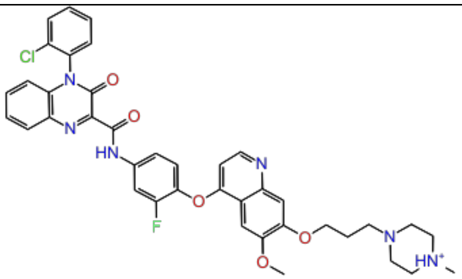 | 6.03 |

| | | | |
|---|---|---|---|
| 37. | CHEMBL4239931 | 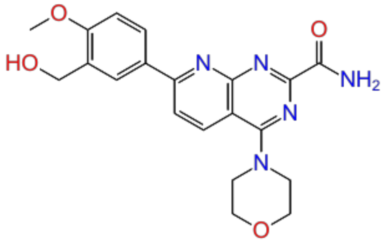 | 6.09 |
| 38. | CHEMBL3612313 | 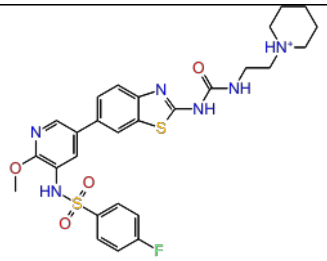 | 6.12 |
| 39. | CHEMBL4226926 | 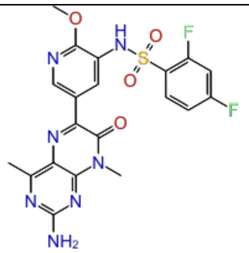 | 6.20 |
| 40. | CHEMBL53463 | 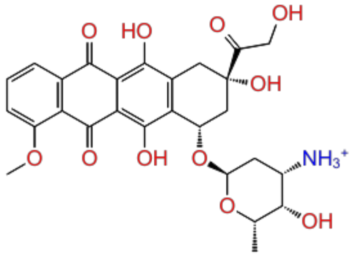 | 6.32 |
| 41. | CHEMBL1615189 | 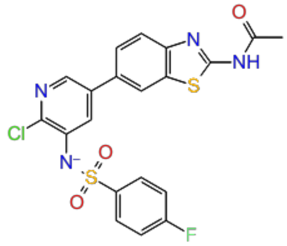 | 6.36 |
| 42. | CHEMBL118655 | 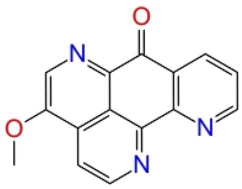 | 6.52 |

| | | | |
|---|---|---|---|
| 43. | CHEMBL210065 | 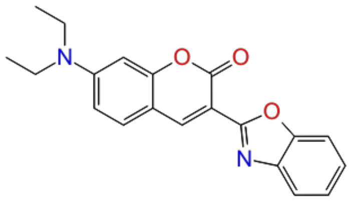 | 6.68 |
| 44. | CHEMBL4169444 | 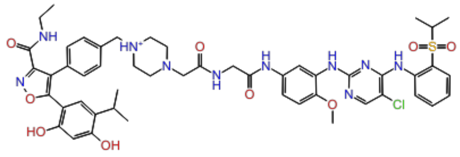 | 6.69 |
| 45. | CHEMBL4251185 | 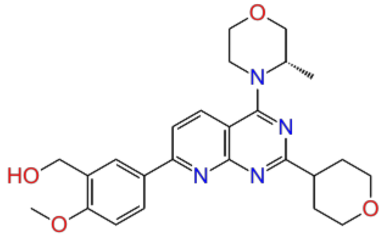 | 7.03 |
| 46. | CHEMBL121911 | 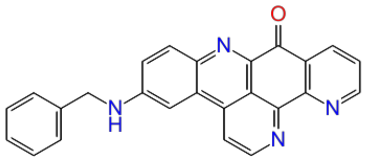 | 7.15 |
| 47. | CHEMBL110525 | 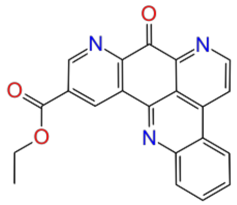 | 7.16 |
| 48. | CHEMBL111477 | 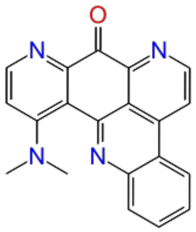 | 7.30 |

| | | | |
|---|---|---|---|
| 49. | CHEMBL117427 | 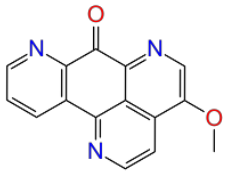 | 7.30 |
| 50. | CHEMBL109640 | 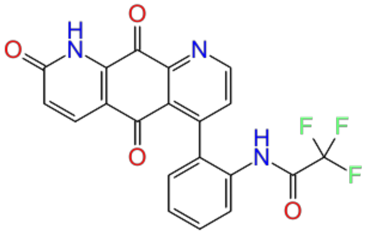 | 7.40 |
| 51. | CHEMBL4250280 | 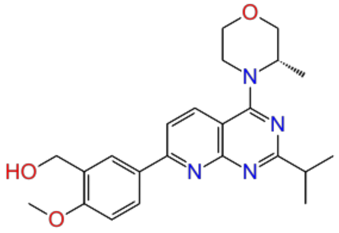 | 7.47 |
| 52. | CHEMBL4166186 | 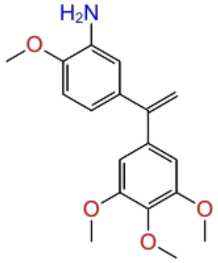 | 8.05 |
| 53. | CHEMBL119700 | 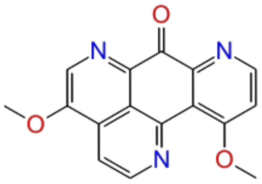 | 8.10 |
| 54. | CHEMBL4238656 | 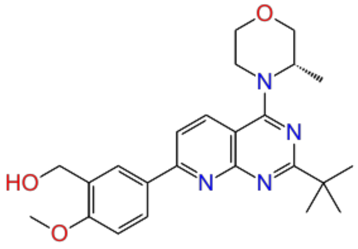 | 8.49 |

| | | | |
|---|---|---|---|
| 55. | CHEMBL1096754 | 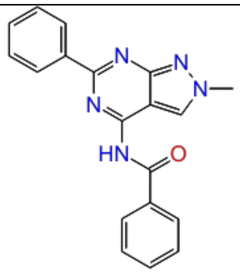 | 9.39 |
| 56. | CHEMBL1096754 | 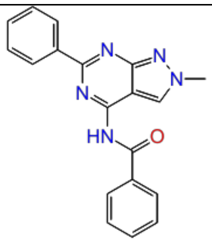 | 9.77 |

**Table S3:** AutoQSAR U87 external test set (169 compounds U87 cell line) along with $pIC_{50}$.

| Sr. No | Compound ID | 2D structure | pIC50 | Dataset Division (Train & Test) |
|---|---|---|---|---|
| 1. | CHEMBL266090 | 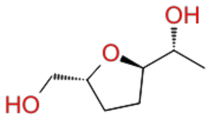 | 3.60 | test |
| 2. | CHEMBL7701 | 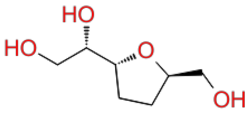 | 3.60 | train |
| 3. | CHEMBL589122 | 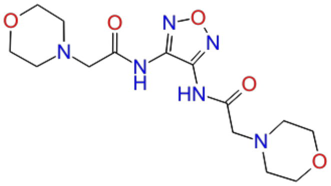 | 3.70 | test |

| | | | | |
|---|---|---|---|---|
| 4. | CHEMBL590093 | 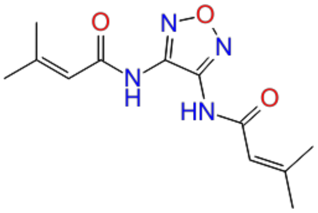 | 3.70 | train |
| 5. | CHEMBL600046 | 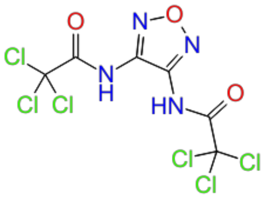 | 3.70 | train |
| 6. | CHEMBL4293736 | 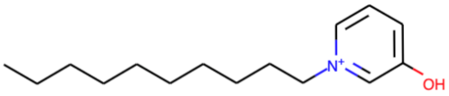 | 3.80 | train |
| 7. | CHEMBL590720 | 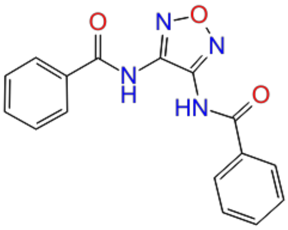 | 3.92 | Test |
| 8. | CHEMBL7476 | 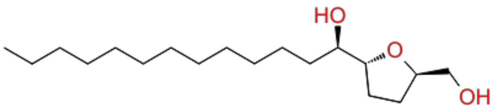 | 3.97 | Train |
| 9. | CHEMBL120862 | 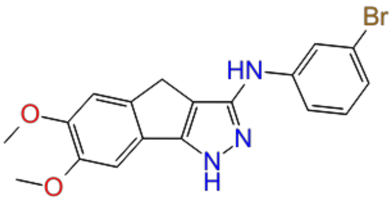 | 4.00 | train |

| | | | | |
|---|---|---|---|---|
| 10. | CHEMBL184948 | 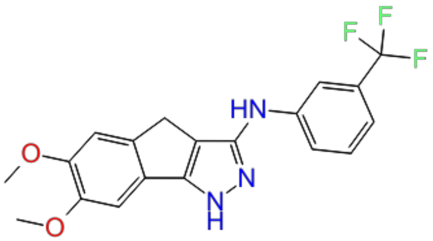 | 4.00 | test |
| 11. | CHEMBL371910 | 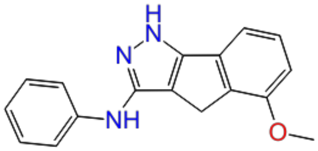 | 4.00 | train |
| 12. | CHEMBL371949 | 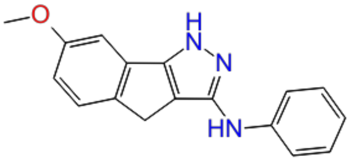 | 4.00 | train |
| 13. | CHEMBL212037 | 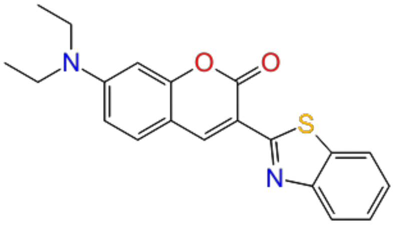 | 4.16 | test |
| 14. | CHEMBL212070 | 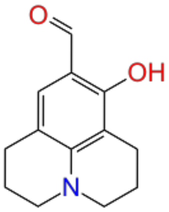 | 4.16 | train |
| 15. | CHEMBL213438 | 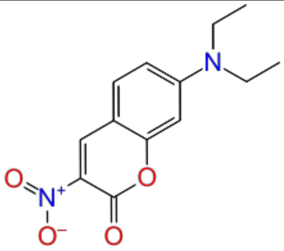 | 4.16 | train |

| | | | | |
|---|---|---|---|---|
| 16. | CHEMBL379350 | 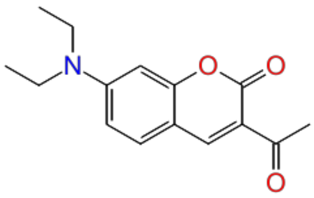 | 4.16 | test |
| 17. | CHEMBL4287207 | 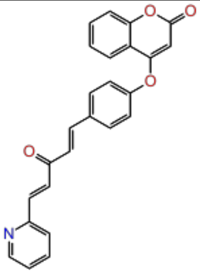 | 4.29 | train |
| 18. | CHEMBL4100030 | 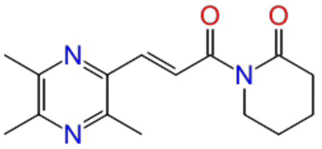 | 4.30 | train |
| 19. | CHEMBL4280398 | 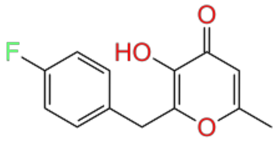 | 4.30 | train |
| 20. | CHEMBL4282653 | 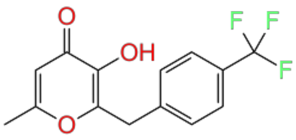 | 4.30 | train |
| 21. | CHEMBL4283735 | 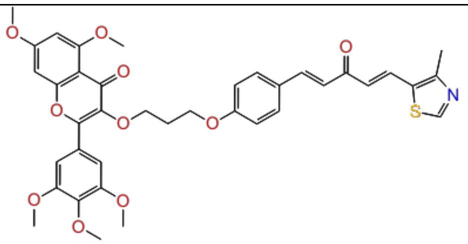 | 4.39 | test |

| | | | | |
|---|---|---|---|---|
| 22. | CHEMBL589107 | 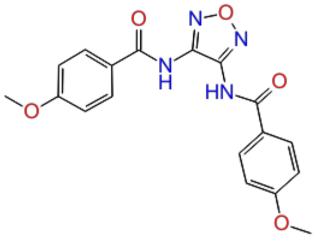 | 4.40 | train |
| 23. | CHEMBL212242 | 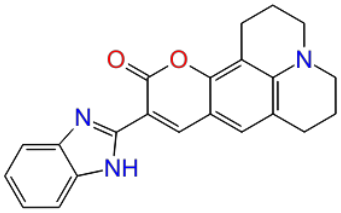 | 4.48 | train |
| 24. | CHEMBL4080967 | 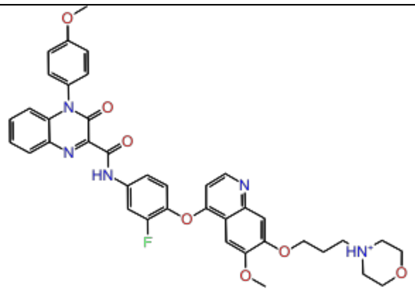 | 4.50 | train |
| 25. | CHEMBL3588940 | 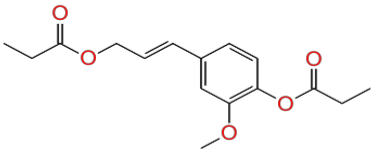 | 4.54 | train |
| 26. | CHEMBL4287242 | 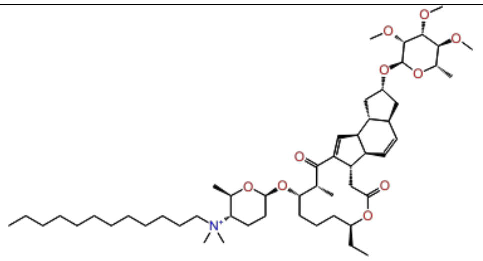 | 4.55 | test |
| 27. | CHEMBL4286444 | 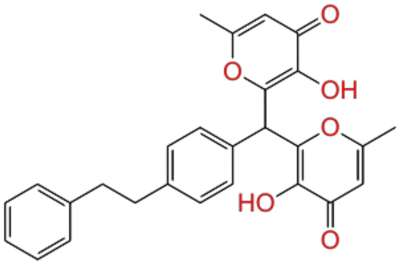 | 4.57 | test |

| | | | | |
|---|---|---|---|---|
| 28. | CHEMBL591985 | 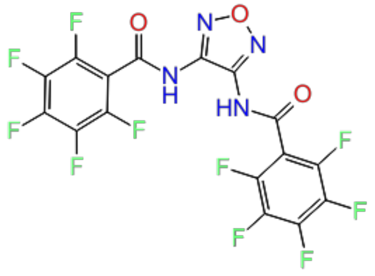 | 4.59 | train |
| 29. | CHEMBL4060108 | 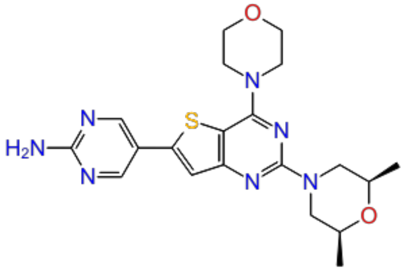 | 4.60 | train |
| 30. | CHEMBL4065513 | 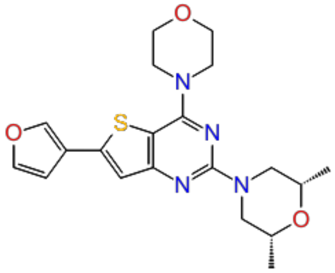 | 4.60 | train |
| 31. | CHEMBL4068220 | 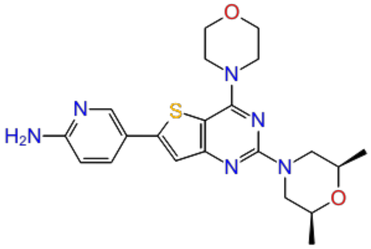 | 4.60 | test |
| 32. | CHEMBL4092492 | 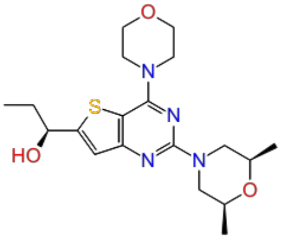 | 4.60 | train |

| | | | | |
|---|---|---|---|---|
| 33. | CHEMBL4100237 | 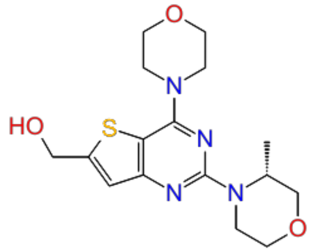 | 4.60 | train |
| 34. | CHEMBL432219 | 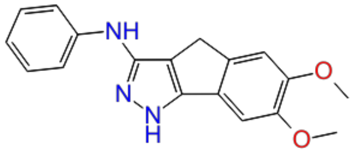 | 4.68 | train |
| 35. | CHEMBL4284792 | 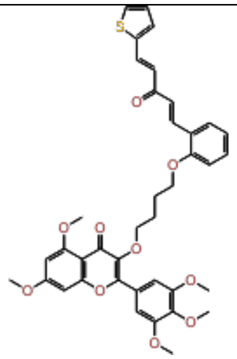 | 4.68 | train |
| 36. | CHEMBL4276849 | 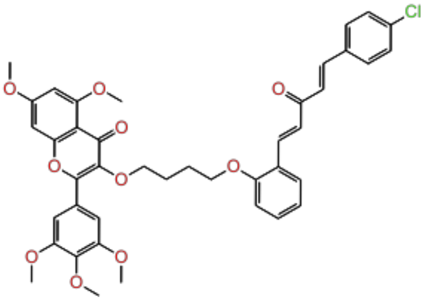 | 4.68 | test |
| 37. | CHEMBL590061 | 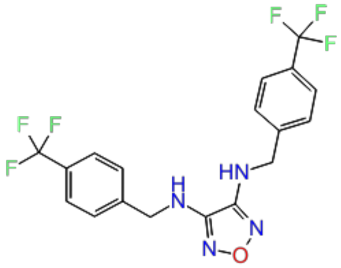 | 4.69 | train |

| | | | | |
|---|---|---|---|---|
| 38. | CHEMBL4283971 | 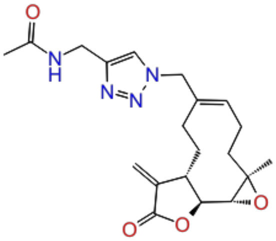 | 4.70 | test |
| 39. | CHEMBL3588941 | 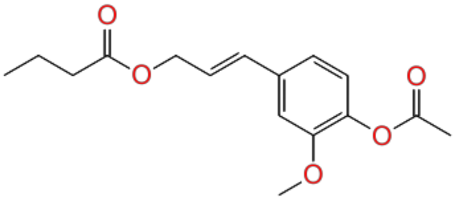 | 4.73 | train |
| 40. | CHEMBL590060 | 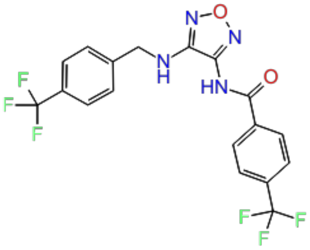 | 4.74 | train |
| 41. | CHEMBL591752 | 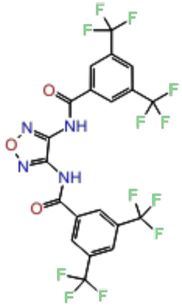 | 4.75 | test |
| 42. | CHEMBL600041 | 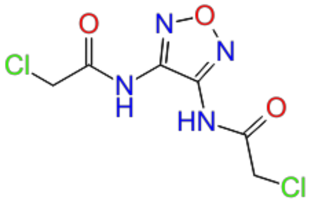 | 4.77 | train |

| | | | | |
|---|---|---|---|---|
| 43. | CHEMBL465843 | 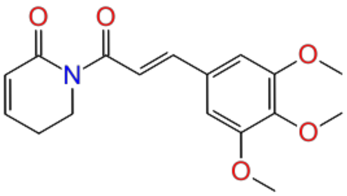 | 4.79 | train |
| 44. | CHEMBL303697 | 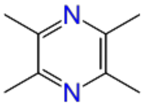 | 4.79 | train |
| 45. | CHEMBL4227076 | 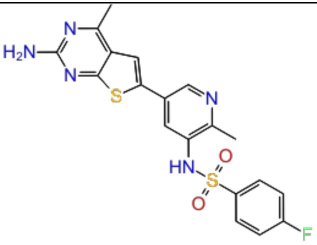 | 4.82 | test |
| 46. | CHEMBL3588943 | 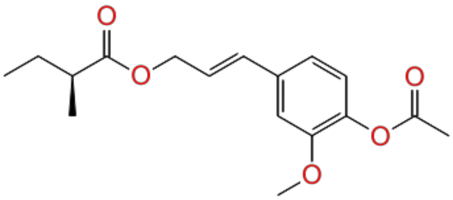 | 4.82 | train |
| 47. | CHEMBL591033 | 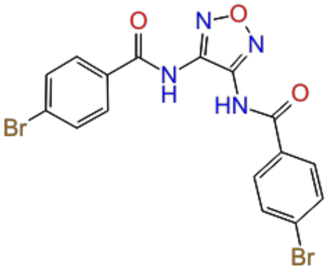 | 4.85 | test |

| | | | | |
|---|---|---|---|---|
| 48. | CHEMBL4294821 | 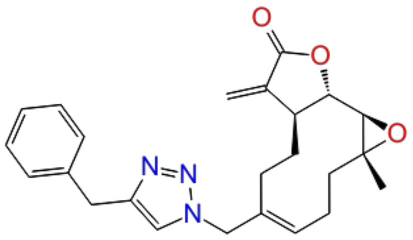 | 4.88 | train |
| 49. | CHEMBL4086949 | 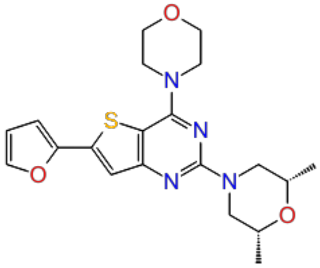 | 4.89 | train |
| 50. | CHEMBL4293259 | 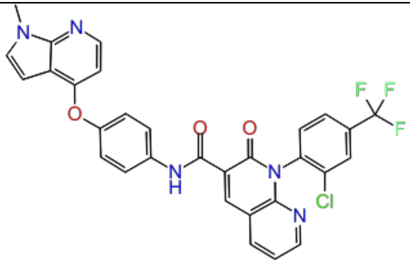 | 4.90 | train |
| 51. | CHEMBL4280764 | 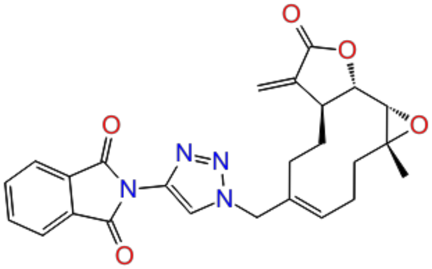 | 4.91 | train |
| 52. | CHEMBL4293238 | 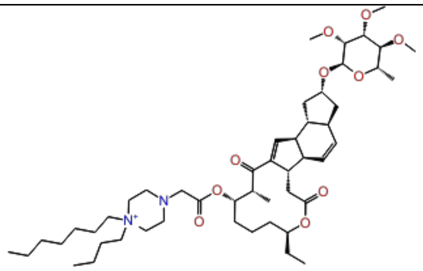 | 4.92 | test |

| | | | | |
|---|---|---|---|---|
| 53. | CHEMBL4214300 | 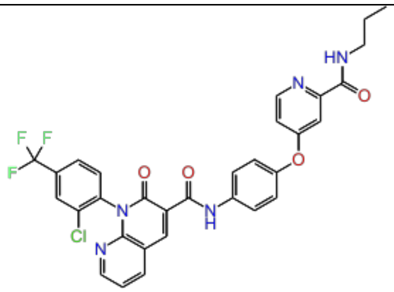 | 4.96 | train |
| 54. | CHEMBL4100335 | 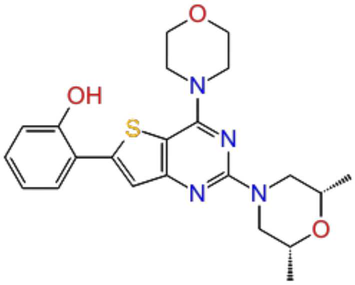 | 5.00 | train |
| 55. | CHEMBL4279750 | 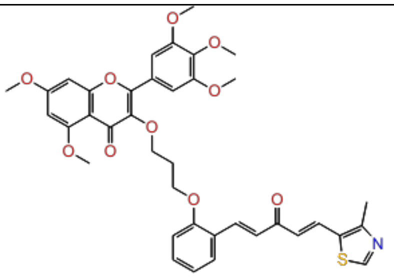 | 5.02 | train |
| 56. | CHEMBL4228112 | 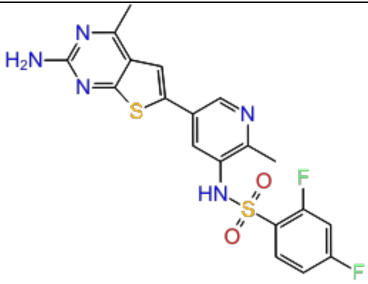 | 5.06 | test |
| 57. | CHEMBL3612318 | 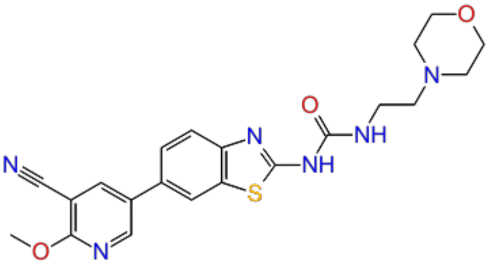 | 5.08 | train |

| | | | | |
|---|---|---|---|---|
| 58. | CHEMBL590806 | 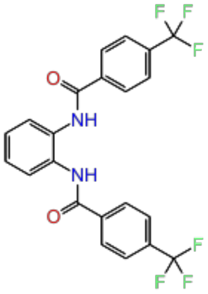 | 5.10 | test |
| 59. | CHEMBL118201 | 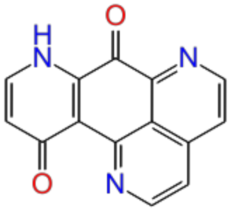 | 5.10 | train |
| 60. | CHEMBL1644111 | 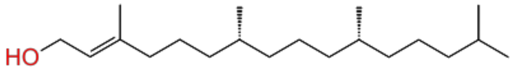 | 5.10 | train |
| 61. | CHEMBL4215210 | 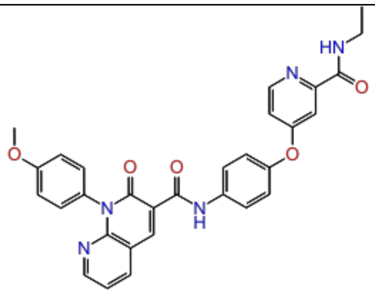 | 5.14 | train |
| 62. | CHEMBL4078390 | 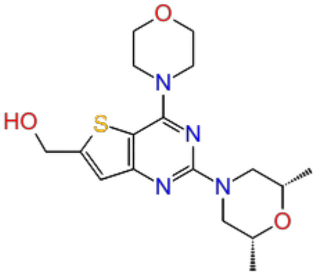 | 5.18 | test |

| | | | | |
|---|---|---|---|---|
| 63. | CHEMBL563939 | 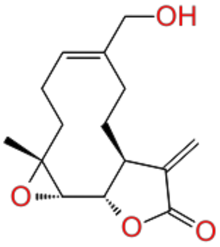 | 5.20 | train |
| 64. | CHEMBL4279674 | 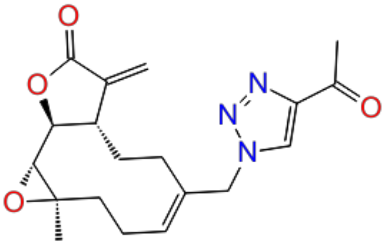 | 5.21 | train |
| 65. | CHEMBL118845 | 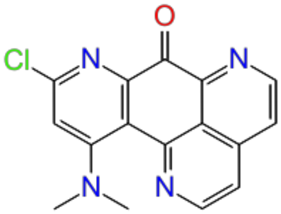 | 5.22 | train |
| 66. | CHEMBL4291410 | 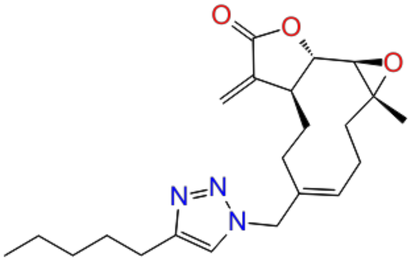 | 5.24 | test |
| 67. | CHEMBL4278116 | 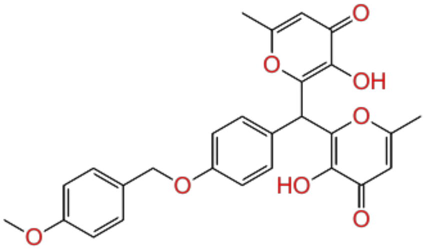 | 5.28 | test |

| | | | | |
|---|---|---|---|---|
| 68. | CHEMBL331562 | 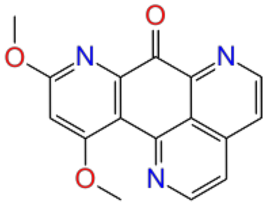 | 5.30 | train |
| 69. | CHEMBL4280439 | 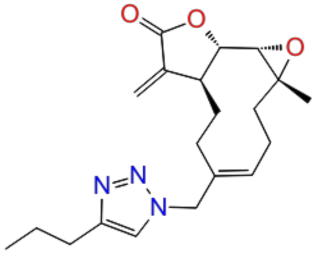 | 5.30 | train |
| 70. | CHEMBL4286032 | 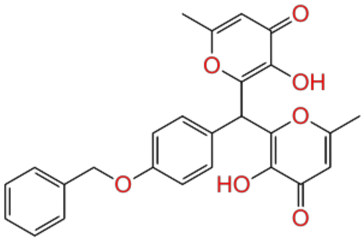 | 5.34 | train |
| 71. | CHEMBL4098035 | 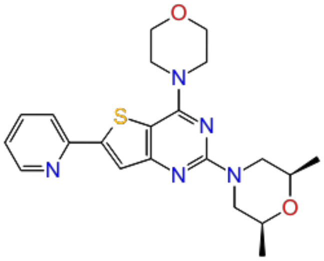 | 5.35 | test |
| 72. | CHEMBL4294566 | 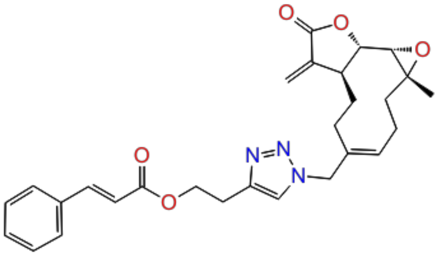 | 5.36 | train |

| | | | | |
|---|---|---|---|---|
| 73. | CHEMBL122557 | 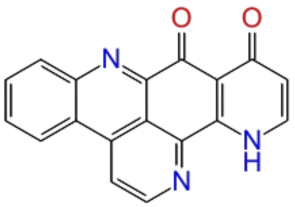 | 5.40 | train |
| 74. | CHEMBL2152488 | 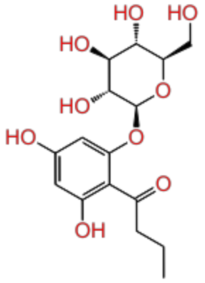 | 5.41 | train |
| 75. | CHEMBL385786 | 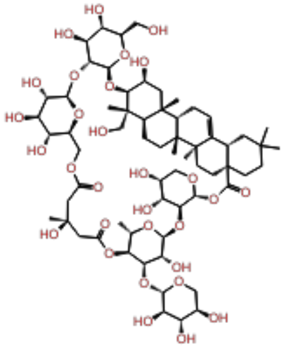 | 5.44 | train |
| 76. | CHEMBL4281496 | 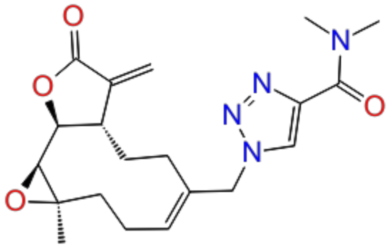 | 5.45 | test |
| 77. | CHEMBL4204238 | 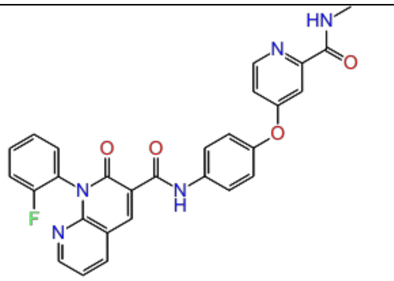 | 5.47 | train |

| | | | | |
|---|---|---|---|---|
| 78. | CHEMBL4064087 | 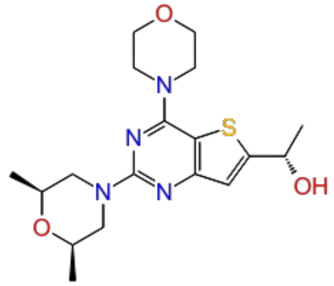 | 5.48 | train |
| 79. | CHEMBL4209984 | 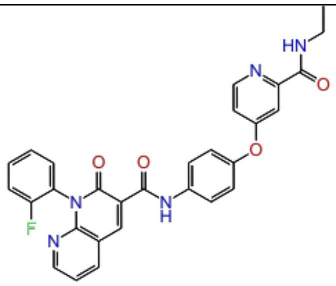 | 5.49 | train |
| 80. | CHEMBL4225944 | 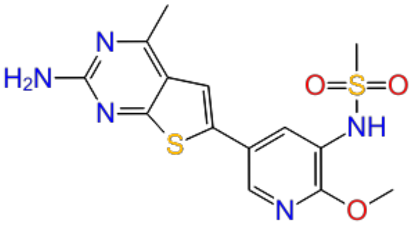 | 5.52 | test |
| 81. | CHEMBL4282461 | 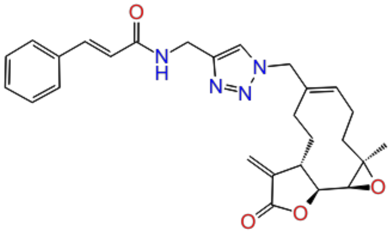 | 5.53 | train |
| 82. | CHEMBL4280635 | 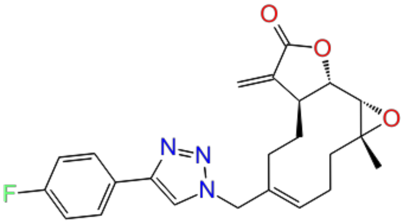 | 5.55 | train |

| | | | | |
|---|---|---|---|---|
| 83. | CHEMBL3659991 | 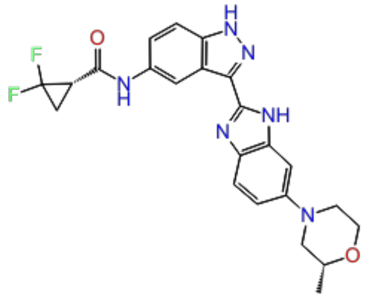 | 5.57 | test |
| 84. | CHEMBL4066144 | 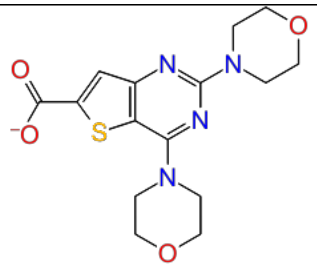 | 5.61 | train |
| 85. | CHEMBL4226082 | 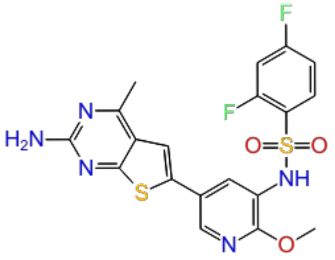 | 5.62 | train |
| 86. | CHEMBL4285130 | 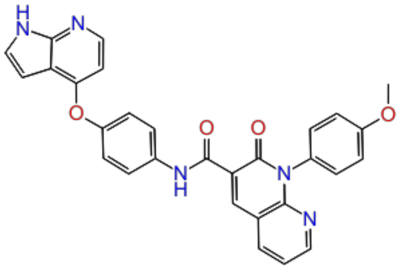 | 5.63 | test |
| 87. | CHEMBL4293054 | 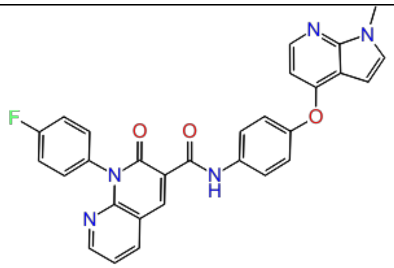 | 5.67 | test |

| | | | | |
|---|---|---|---|---|
| 88. | CHEMBL4225722 | 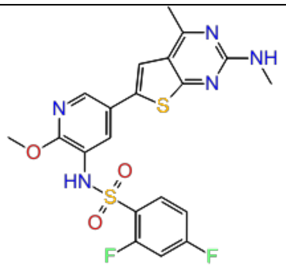 | 5.68 | train |
| 89. | CHEMBL111249 | 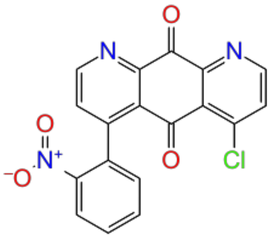 | 5.70 | train |
| 90. | CHEMBL4288866 | 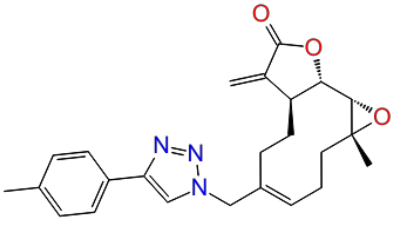 | 5.72 | train |
| 91. | CHEMBL4072336 | 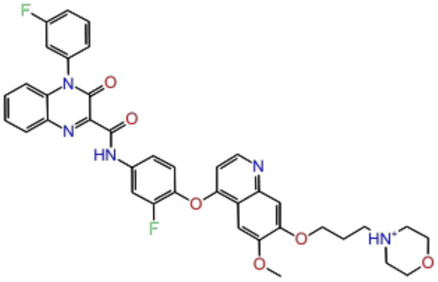 | 5.73 | test |
| 92. | CHEMBL373587 | 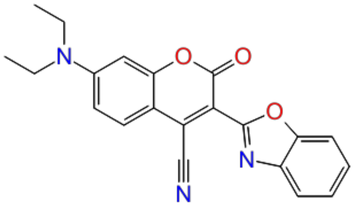 | 5.75 | train |

| | | | | |
|---|---|---|---|---|
| 93. | CHEMBL4074866 | 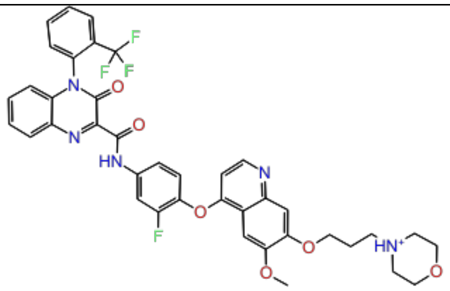 | 5.75 | train |
| 94. | CHEMBL3612319 | 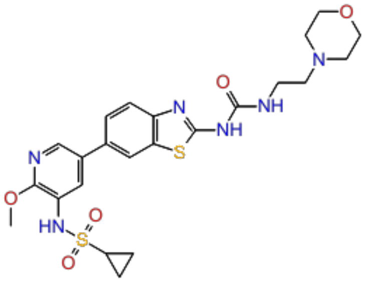 | 5.78 | train |
| 95. | CHEMBL4084146 | 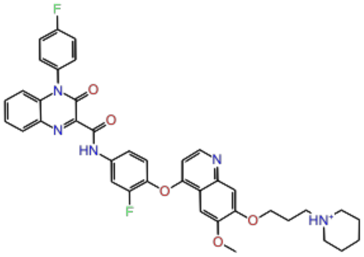 | 5.79 | train |
| 96. | CHEMBL4097578 | 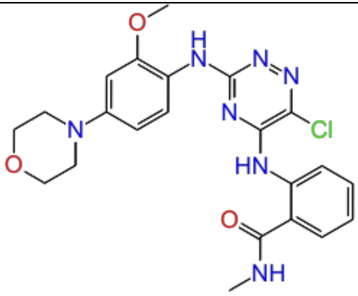 | 5.82 | test |
| 97. | CHEMBL4207773 | 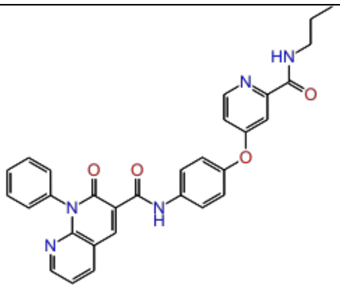 | 5.82 | train |

| | | | | |
|---|---|---|---|---|
| 98. | CHEMBL3612194 | 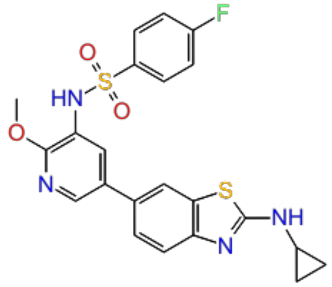 | 5.82 | train |
| 99. | CHEMBL4288118 | 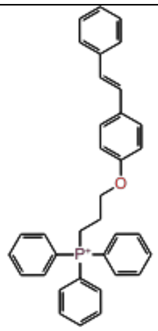 | 5.85 | train |
| 100. | CHEMBL4075562 | 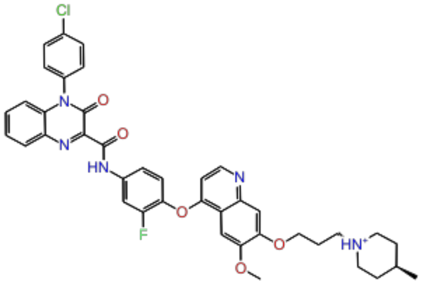 | 5.85 | test |
| 101. | CHEMBL3612197 | 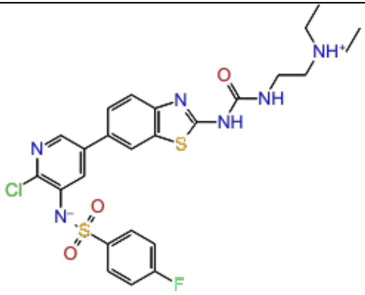 | 5.86 | test |
| 102. | CHEMBL4289873 | 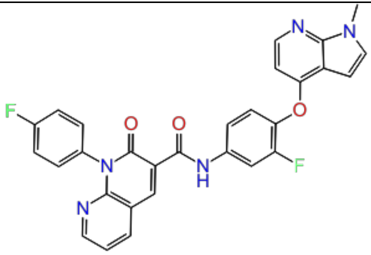 | 5.88 | train |

| | | | | |
|---|---|---|---|---|
| 103. | CHEMBL4277119 | 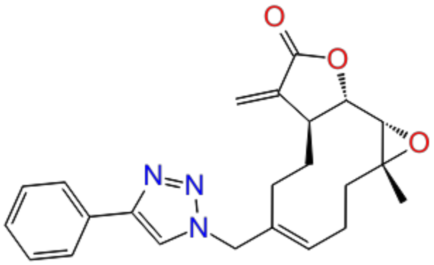 | 5.90 | train |
| 104. | CHEMBL4278233 | 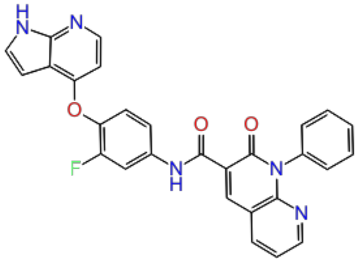 | 5.90 | test |
| 105. | CHEMBL4285131 | 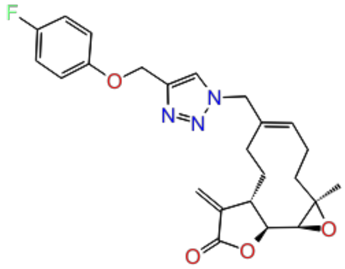 | 5.90 | train |
| 106. | CHEMBL4294165 | 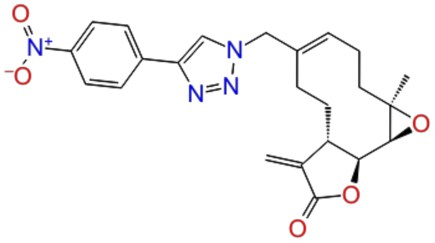 | 5.91 | train |
| 107. | CHEMBL4059832 | 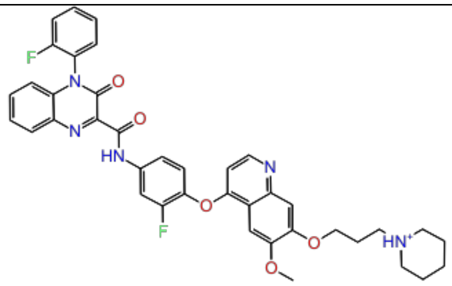 | 5.93 | train |

| | | | | |
|---|---|---|---|---|
| 108. | CHEMBL4214671 | 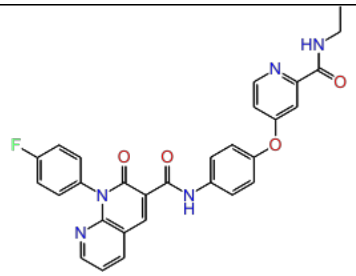 | 5.95 | train |
| 109. | CHEMBL3612315 | 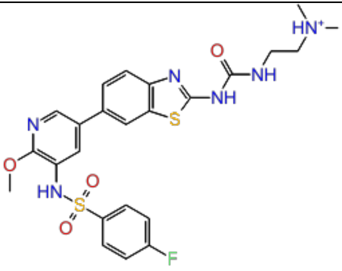 | 5.99 | test |
| 110. | CHEMBL4244147 | 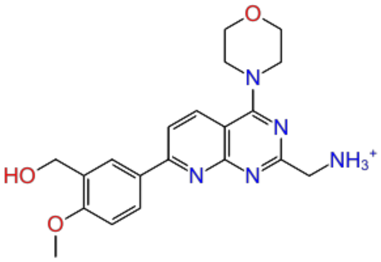 | 6.00 | train |
| 111. | CHEMBL4242668 | 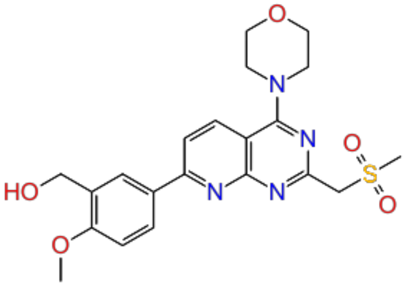 | 6.00 | train |
| 112. | CHEMBL4225819 | 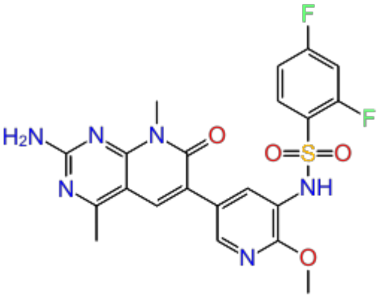 | 6.02 | test |

| | | | | |
|---|---|---|---|---|
| 113. | CHEMBL4204920 | 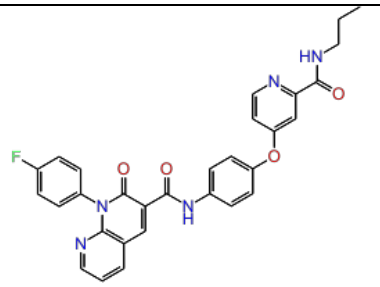 | 6.02 | train |
| 114. | CHEMBL1230609 | 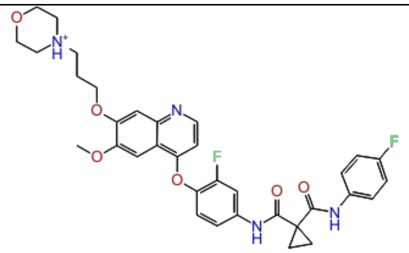 | 6.02 | train |
| 115. | CHEMBL4286698 | 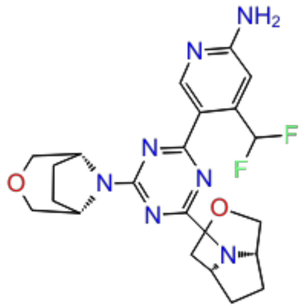 | 6.04 | train |
| 116. | CHEMBL4239569 | 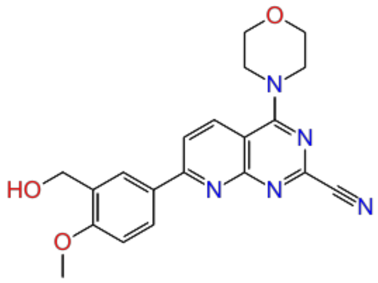 | 6.04 | test |
| 117. | CHEMBL4098250 | 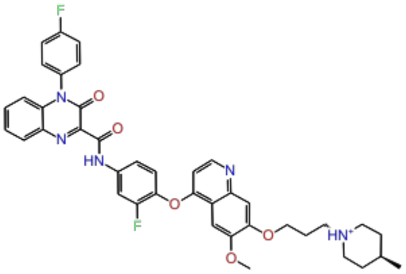 | 6.06 | train |

| | | | | |
|---|---|---|---|---|
| 118. | CHEMBL117616 | 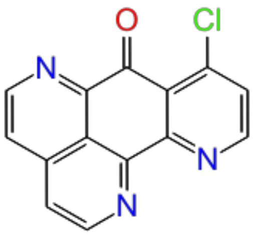 | 6.10 | train |
| 119. | CHEMBL118709 | 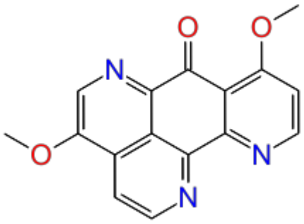 | 6.10 | train |
| 120. | CHEMBL111300 | 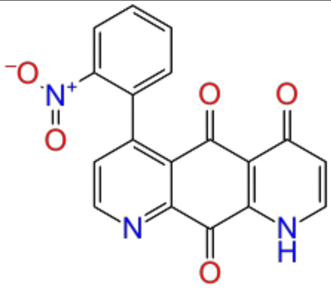 | 6.10 | test |
| 121. | CHEMBL111383 | 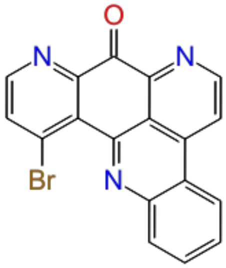 | 6.16 | train |
| 122. | CHEMBL4065141 | 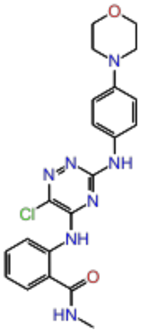 | 6.16 | train |

| | | | | |
|---|---|---|---|---|
| 123. | CHEMBL4225483 | 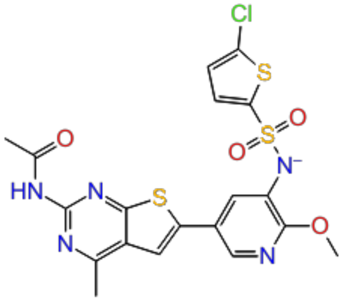 | 6.16 | test |
| 124. | CHEMBL4092858 | 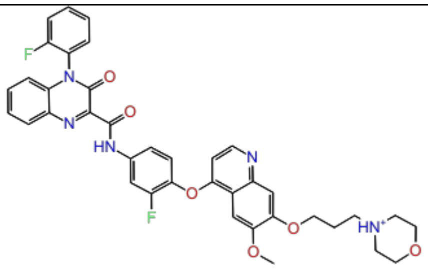 | 6.19 | train |
| 125. | CHEMBL4280452 | 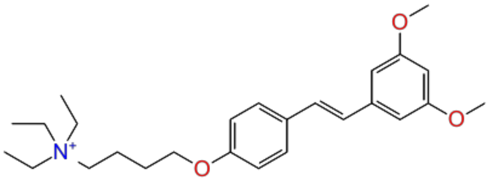 | 6.24 | test |
| 126. | CHEMBL521851 | 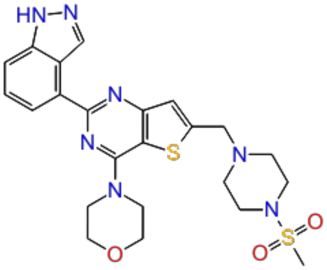 | 6.26 | train |
| 127. | CHEMBL3612321 | 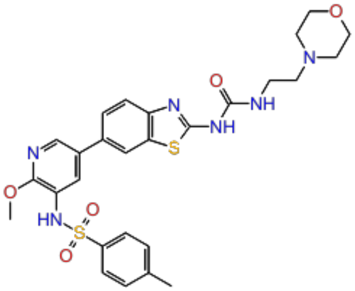 | 6.28 | train |

| | | | | |
|---|---|---|---|---|
| 128. | CHEMBL325202 | 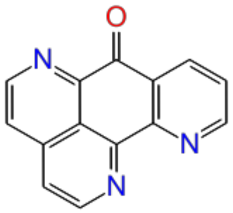 | 6.30 | test |
| 129. | CHEMBL4092812 | 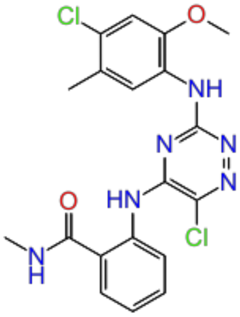 | 6.30 | train |
| 130. | CHEMBL3612322 | 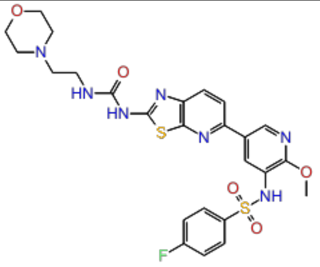 | 6.32 | train |
| 131. | CHEMBL4284998 | 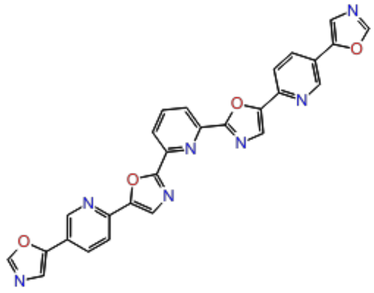 | 6.36 | test |
| 132. | CHEMBL3612314 | 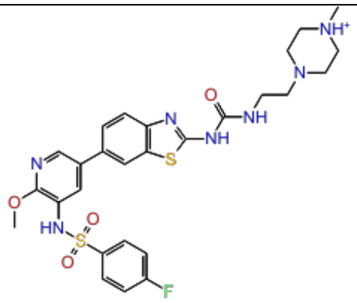 | 6.36 | train |

| | | | | |
|---|---|---|---|---|
| 133. | CHEMBL4071083 | 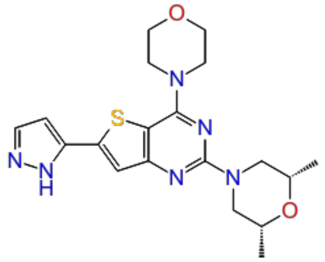 | 6.37 | train |
| 134. | CHEMBL4291927 | 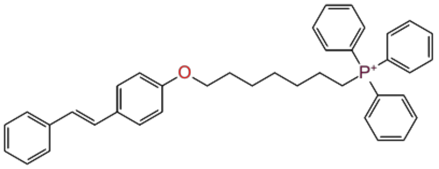 | 6.44 | train |
| 135. | CHEMBL3612312 | 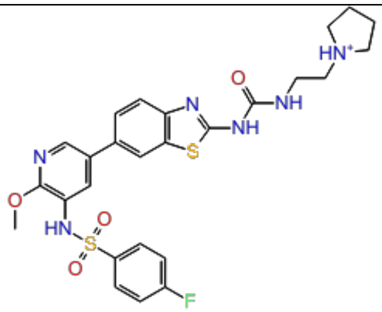 | 6.44 | train |
| 136. | CHEMBL1234354 | 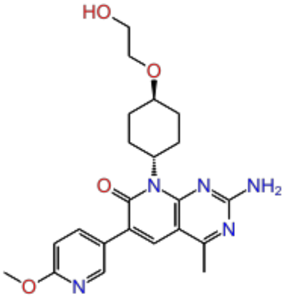 | 6.47 | test |
| 137. | CHEMBL112938 | 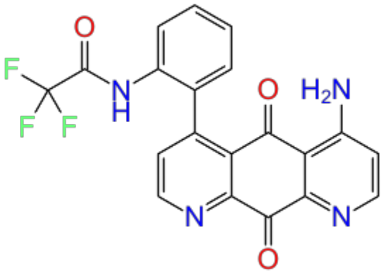 | 6.52 | train |

| | | | | |
|---|---|---|---|---|
| 138. | CHEMBL4284890 | 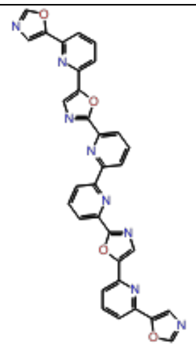 | 6.61 | train |
| 139. | CHEMBL4159104 | 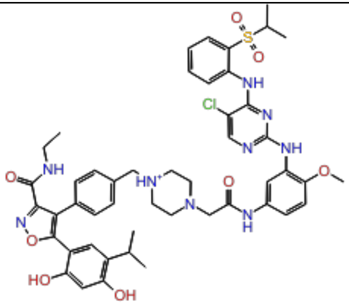 | 6.62 | train |
| 140. | CHEMBL359744 | 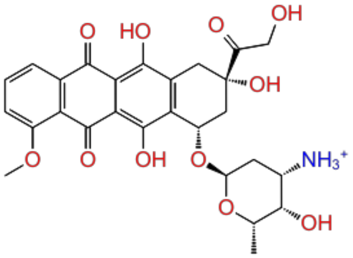 | 6.80 | test |
| 141. | CHEMBL4072042 | 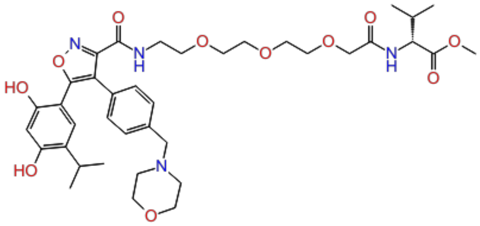 | 6.96 | train |
| 142. | CHEMBL327101 | 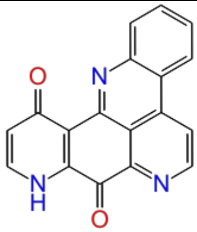 | 7.00 | train |

| | | | | |
|---|---|---|---|---|
| 143. | CHEMBL117784 | 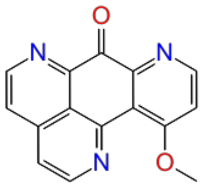 | 7.00 | train |
| 144. | CHEMBL4280053 | 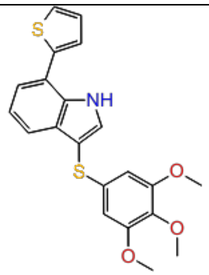 | 7.02 | test |
| 145. | CHEMBL4290427 | 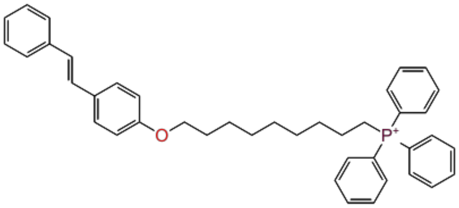 | 7.05 | test |
| 146. | CHEMBL4246481 | 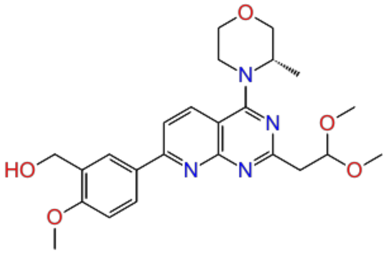 | 7.05 | train |
| 147. | CHEMBL4246121 | 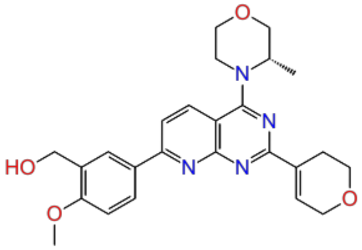 | 7.07 | train |

| | | | | |
|---|---|---|---|---|
| 148. | CHEMBL122076 | 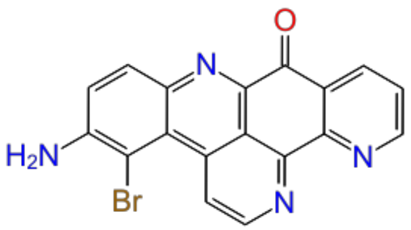 | 7.10 | train |
| 149. | CHEMBL125395 | 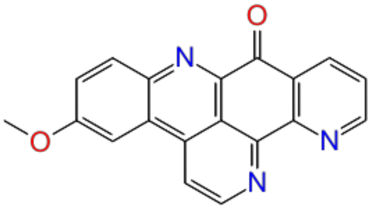 | 7.16 | train |
| 150. | CHEMBL121911 | 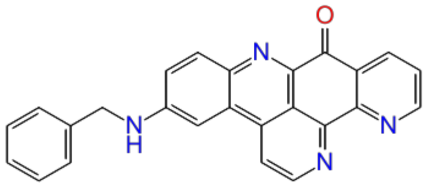 | 7.16 | train |
| 151. | CHEMBL340160 | 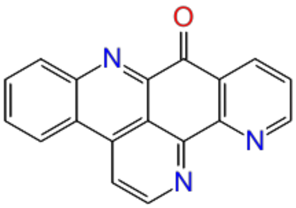 | 7.16 | test |
| 152. | CHEMBL4060227 | 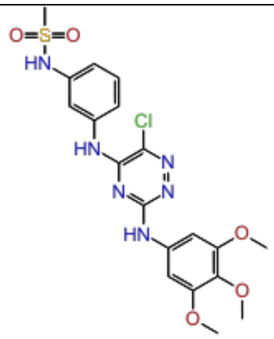 | 7.16 | train |

| | | | | |
|---|---|---|---|---|
| 153. | CHEMBL4281312 | 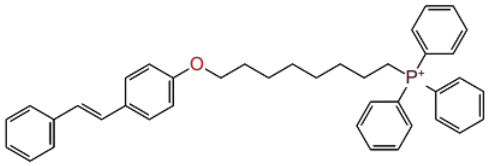 | 7.16 | test |
| 154. | CHEMBL323280 | 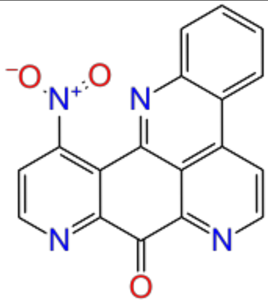 | 7.16 | train |
| 155. | CHEMBL324780 | 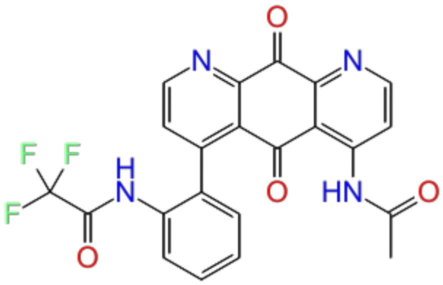 | 7.22 | test |
| 156. | CHEMBL324884 | 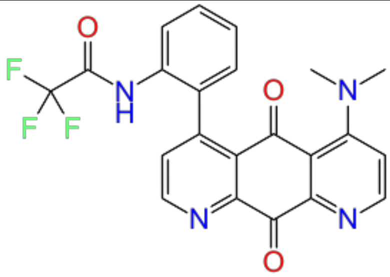 | 7.22 | train |
| 157. | CHEMBL4280206 | 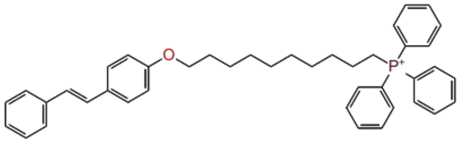 | 7.22 | train |

| | | | | |
|---|---|---|---|---|
| 158. | CHEMBL333669 | 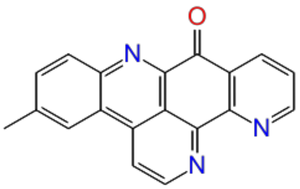 | 7.22 | test |
| 159. | CHEMBL109479 | 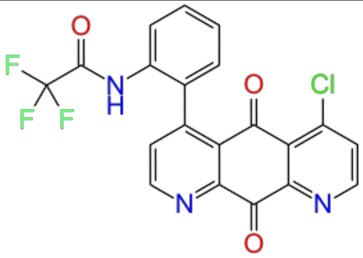 | 7.30 | train |
| 160. | CHEMBL3301610 | 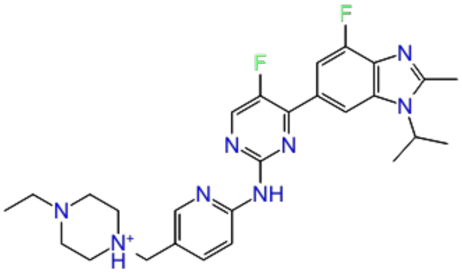 | 7.32 | train |
| 161. | CHEMBL4239712 | 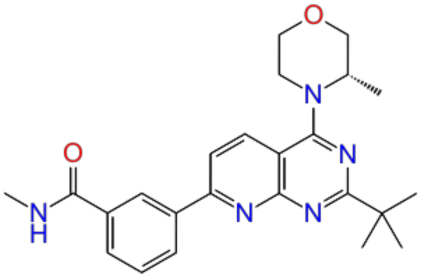 | 7.49 | train |
| 162. | CHEMBL1801204 | 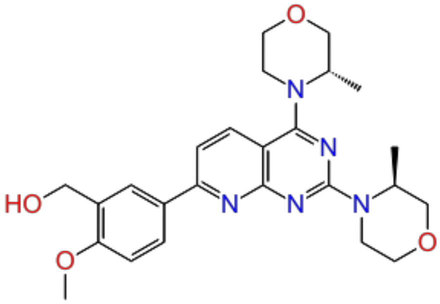 | 7.67 | train |

| | | | | |
|---|---|---|---|---|
| 163. | CHEMBL4278554 | 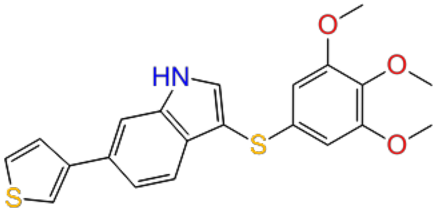 | 7.80 | train |
| 164. | CHEMBL4098626 | 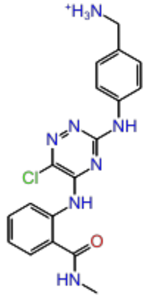 | 7.85 | test |
| 165. | CHEMBL498271 | 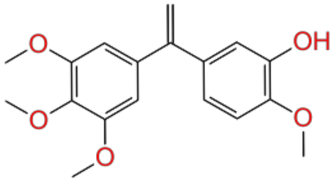 | 8.10 | test |
| 166. | CHEMBL4162270 | 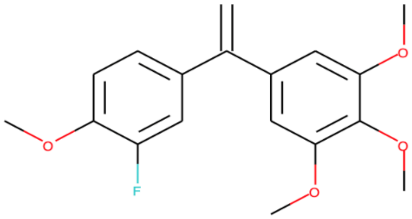 | 8.10 | train |
| 167. | CHEMBL4238656 | 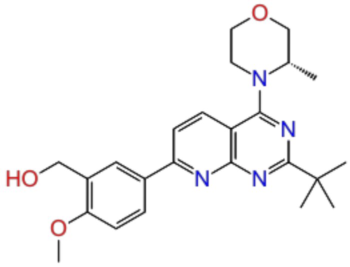 | 8.49 | train |
| 168. | CHEMBL4172942 | 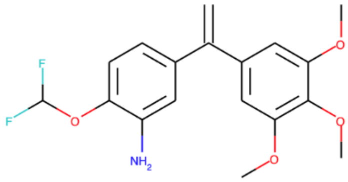 | 9.22 | train |

| | | | | |
|---|---|---|---|---|
| 169. | CHEMBL508617 | 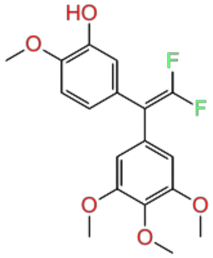 | 9.52 | train |

**Table S4:** Construction of AutoQSAR-U251 using machine learning based algorithm (kpls_molprint2D_36) U251 cell line.

| Sr. No | Compound ID | 2D structure | pIC50 | Dataset Division (Train & Test) |
|---|---|---|---|---|
| 1. | CHEMBL20883 | 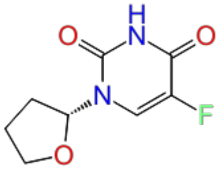 | 3.42 | train |
| 2. | CHEMBL253353 | 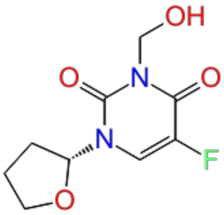 | 3.99 | test |
| 3. | CHEMBL500473 | 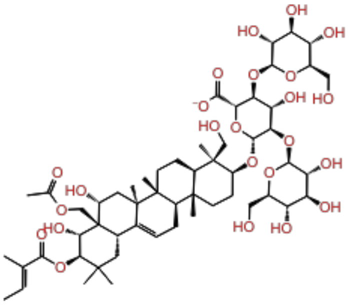 | 4.09 | test |

| | | | | |
|---|---|---|---|---|
| 4. | CHEMBL50 | 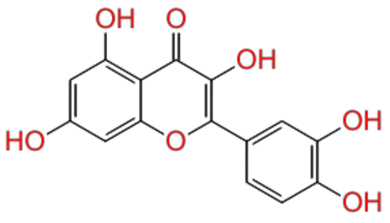 | 4.11 | train |
| 5. | CHEMBL509354 | 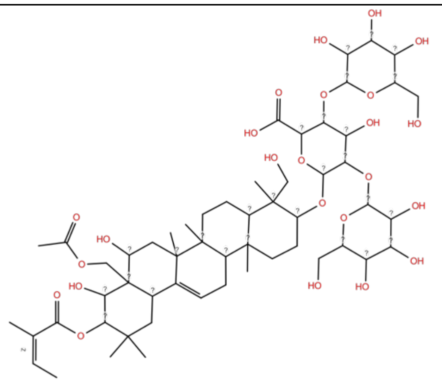 | 4.16 | train |
| 6. | CHEMBL180828 | 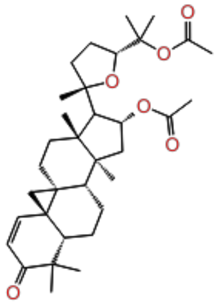 | 4.23 | train |
| 7. | CHEMBL252368 | 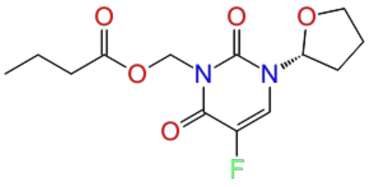 | 4.26 | train |
| 8. | CHEMBL3588940 | 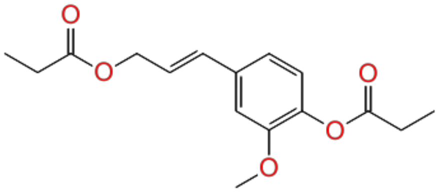 | 4.28 | test |

| | | | | |
|---|---|---|---|---|
| 9. | CHEMBL75267 | 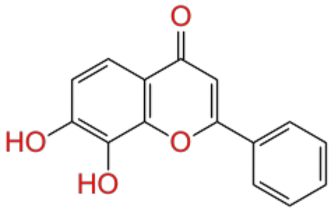 | 4.30 | train |
| 10. | CHEMBL432108 | 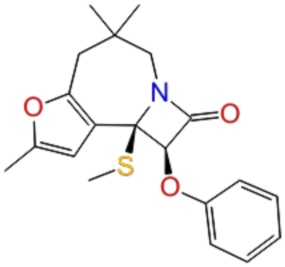 | 4.33 | train |
| 11. | CHEMBL1958318 | 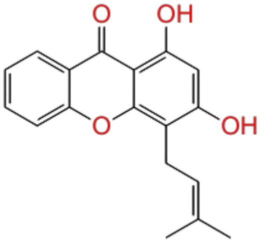 | 4.33 | test |
| 12. | CHEMBL3588942 | 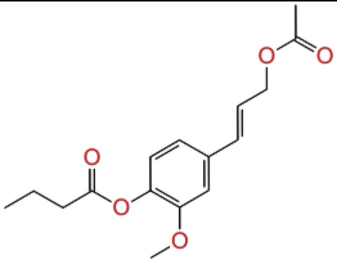 | 4.34 | train |
| 13. | CHEMBL412883 | 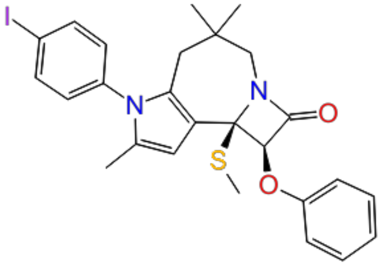 | 4.40 | train |

| | | | | |
|---|---|---|---|---|
| 14. | CHEMBL233930 | 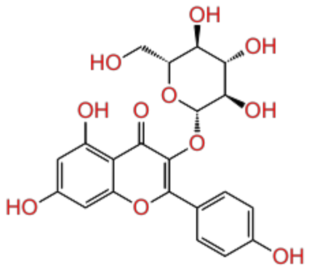 | 4.42 | train |
| 15. | CHEMBL4093795 | 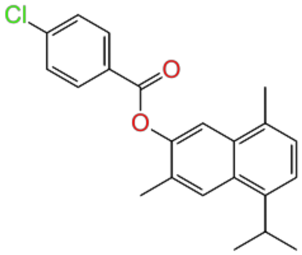 | 4.45 | train |
| 16. | CHEMBL3588941 | 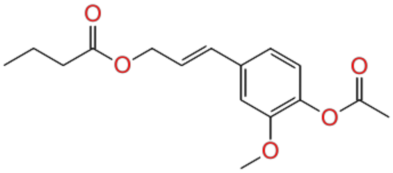 | 4.48 | test |
| 17. | CHEMBL1784141 | 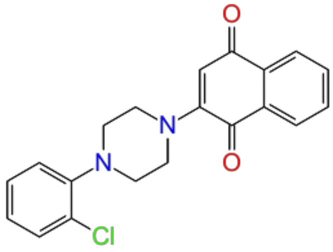 | 4.51 | train |
| 18. | CHEMBL359600 | 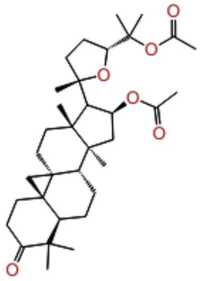 | 4.52 | test |

| | | | | |
|---|---|---|---|---|
| 19. | CHEMBL3588943 | 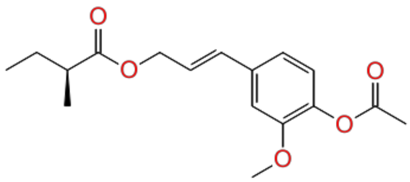 | 4.55 | train |
| 20. | CHEMBL187044 | 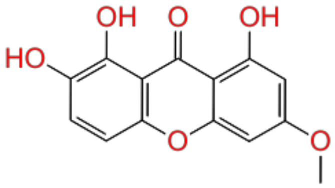 | 4.56 | test |
| 21. | CHEMBL1644111 | 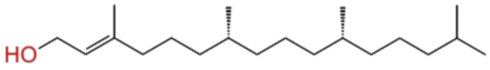 | 4.56 | train |
| 22. | CHEMBL182093 | 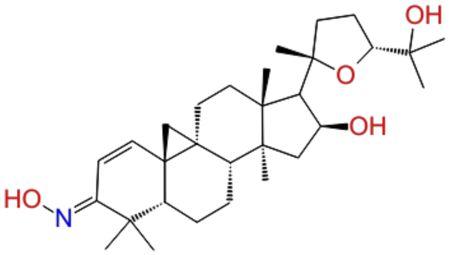 | 4.57 | train |
| 23. | CHEMBL3588938 | 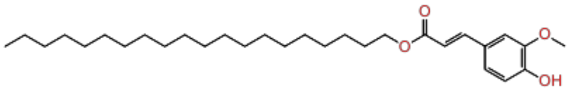 | 4.57 | train |

| | | | | |
|---|---|---|---|---|
| 24. | CHEMBL298047 | 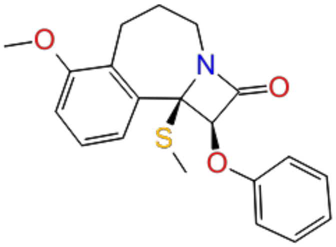 | 4.60 | train |
| 25. | CHEMBL1958315 | 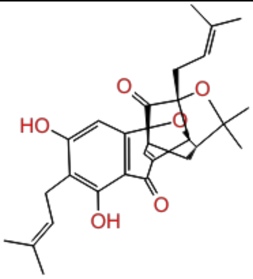 | 4.65 | test |
| 26. | CHEMBL4103953 | 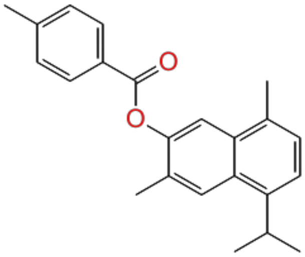 | 4.65 | train |
| 27. | CHEMBL2152488 | 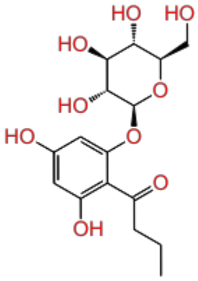 | 4.71 | train |
| 28. | CHEMBL1958317 | 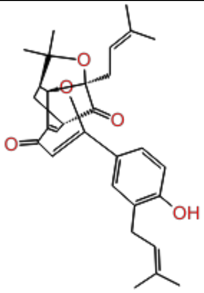 | 4.73 | train |

| | | | | |
|---|---|---|---|---|
| 29. | CHEMBL1958319 | 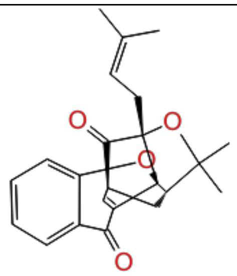 | 4.73 | test |
| 30. | CHEMBL4087813 | 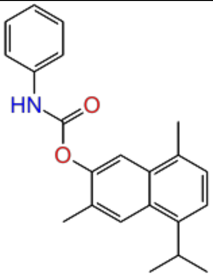 | 4.80 | train |
| 31. | CHEMBL4172865 | 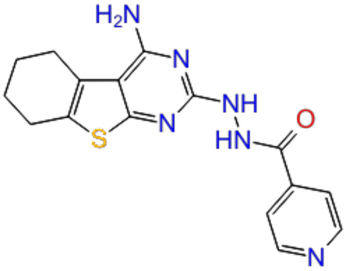 | 4.88 | train |
| 32. | CHEMBL3331344 | 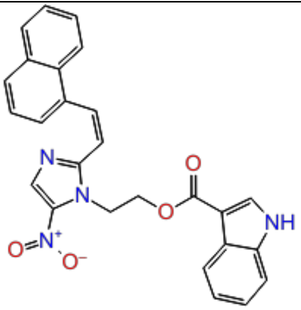 | 4.89 | test |
| 33. | CHEMBL178232 | 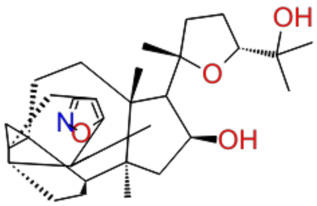 | 4.95 | train |

| | | | | |
|---|---|---|---|---|
| 34. | CHEMBL1784156 | 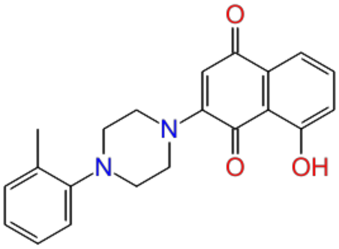 | 4.99 | train |
| 35. | CHEMBL3311152 | 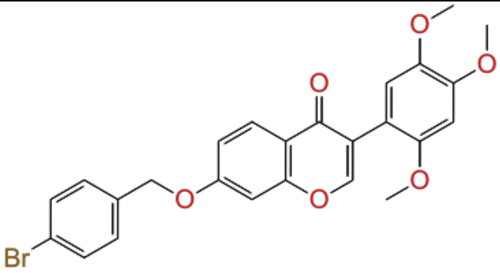 | 5.00 | train |
| 36. | CHEMBL3588939 | 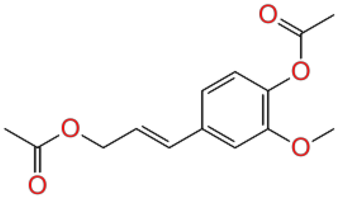 | 5.01 | test |
| 37. | CHEMBL3311154 | 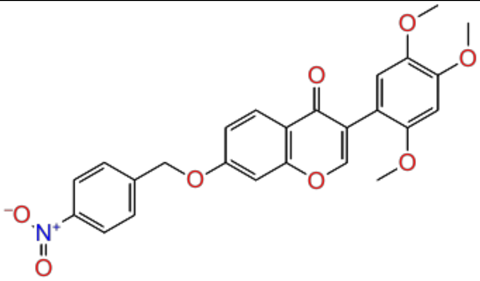 | 5.03 | test |
| 38. | CHEMBL1784154 | 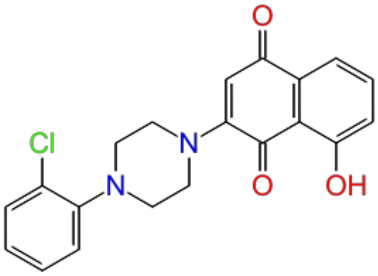 | 5.10 | train |

| | | | | |
|---|---|---|---|---|
| 39. | CHEMBL3311147 | 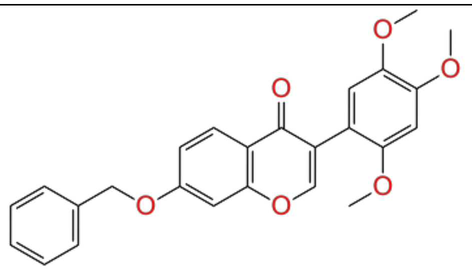 | 5.17 | train |
| 40. | CHEMBL3311146 | 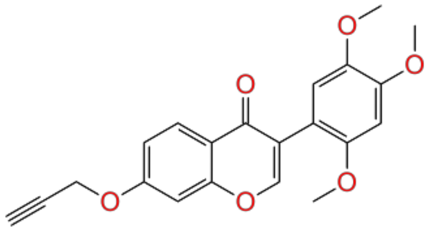 | 5.19 | test |
| 41. | CHEMBL1784150 | 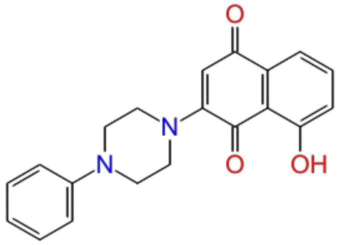 | 5.21 | train |
| 42. | CHEMBL1784161 | 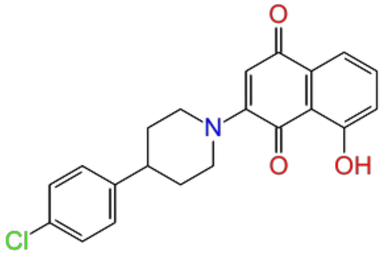 | 5.21 | train |
| 43. | CHEMBL3311157 | 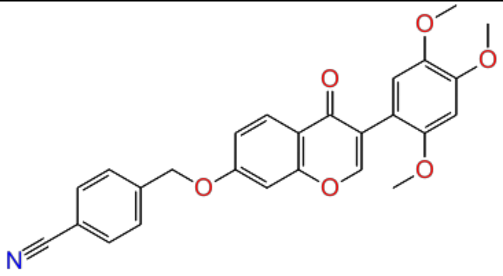 | 5.21 | train |

| | | | | |
|---|---|---|---|---|
| 44. | CHEMBL3311145 | 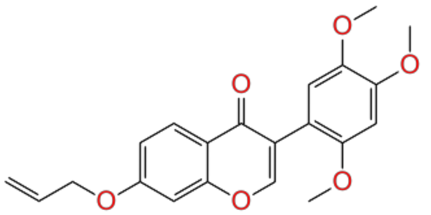 | 5.23 | train |
| 45. | CHEMBL446125 | 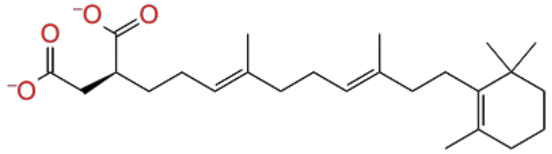 | 5.39 | test |
| 46. | CHEMBL3311038 | 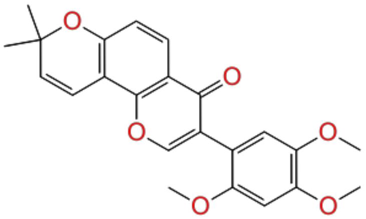 | 5.39 | train |
| 47. | CHEMBL3330998 | 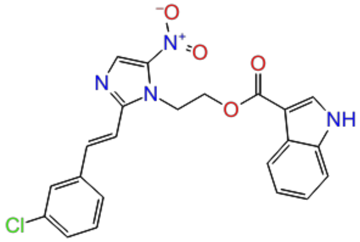 | 5.39 | test |
| 48. | CHEMBL3331001 | 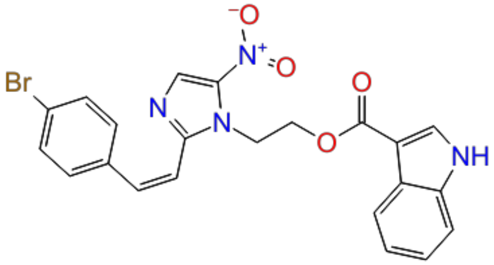 | 5.41 | train |

| | | | | |
|---|---|---|---|---|
| 49. | CHEMBL258765 | 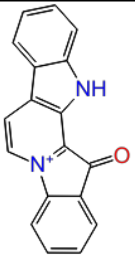 | 5.44 | train |
| 50. | CHEMBL3311061 | 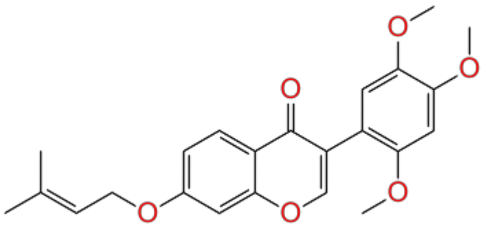 | 5.46 | train |
| 51. | CHEMBL3329220 | 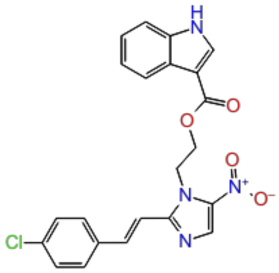 | 5.49 | train |
| 52. | CHEMBL3311151 | 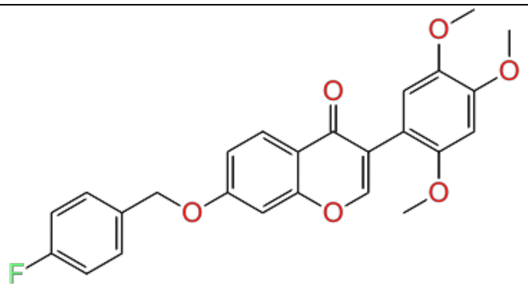 | 5.50 | train |
| 53. | CHEMBL3330999 | 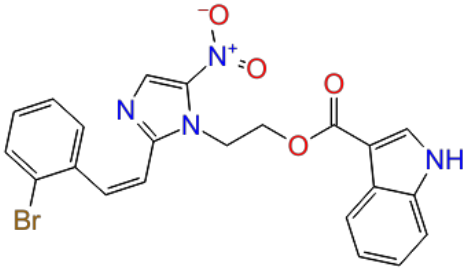 | 5.51 | test |

| | | | | |
|---|---|---|---|---|
| 54. | CHEMBL3331010 | 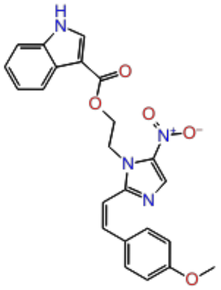 | 5.51 | train |
| 55. | CHEMBL3330997 | 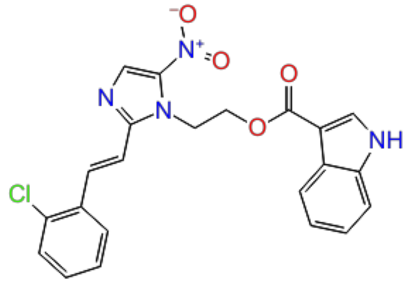 | 5.53 | train |
| 56. | CHEMBL3331003 | 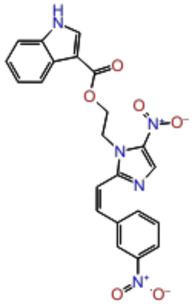 | 5.55 | test |
| 57. | CHEMBL3311052 | 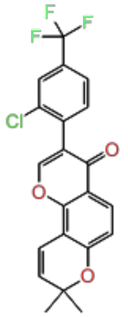 | 5.60 | train |
| 58. | CHEMBL3331004 | 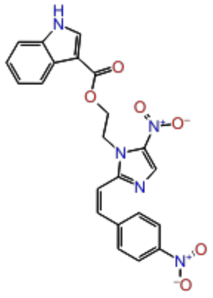 | 5.69 | train |

| | | | | |
|---|---|---|---|---|
| 59. | CHEMBL1164847 | 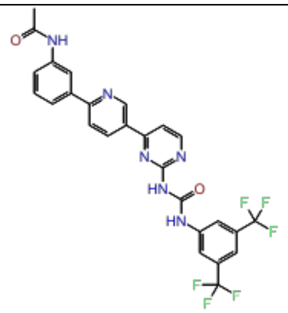 | 5.70 | test |
| 60. | CHEMBL1271866 | 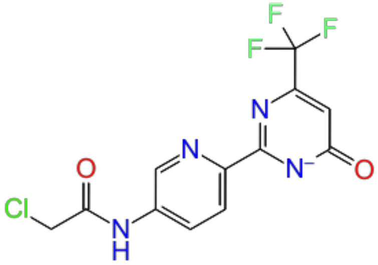 | 5.82 | train |
| 61. | CHEMBL3331002 | 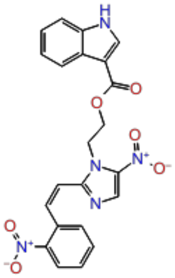 | 5.96 | train |
| 62. | CHEMBL3331011 | 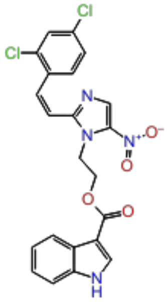 | 6.06 | train |
| 63. | CHEMBL1956190 | 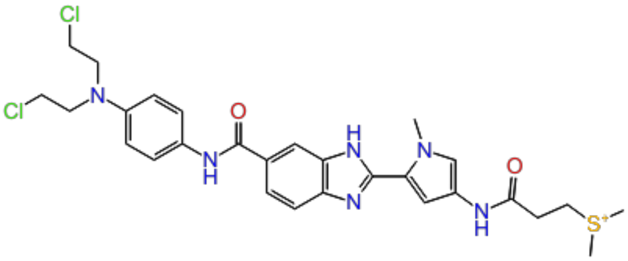 | 6.15 | test |

| | | | | |
|---|---|---|---|---|
| 64. | CHEMBL53463 | 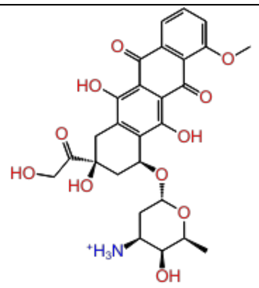 | 6.22 | train |
| 65. | CHEMBL84 | 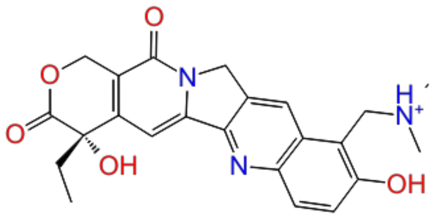 | 6.30 | train |
| 66. | CHEMBL3311160 | 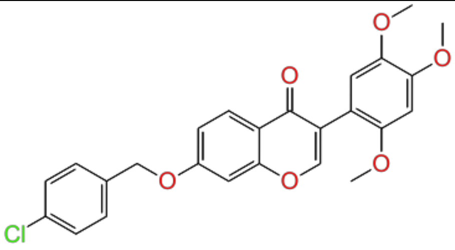 | 6.33 | train |
| 67. | CHEMBL1278024 | 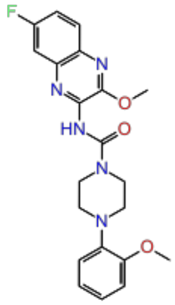 | 6.85 | test |
| 68. | CHEMBL3758224 | 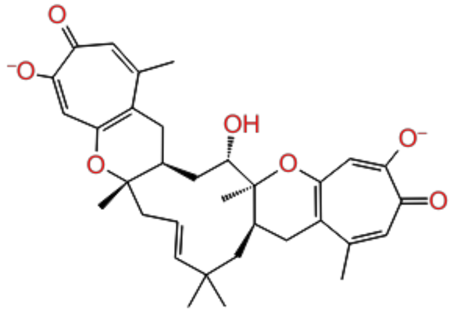 | 7.00 | train |

| | | | | |
|---|---|---|---|---|
| 69. | CHEMBL1276954 | 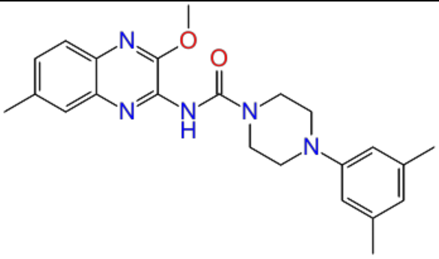 | 7.07 | test |
| 70. | CHEMBL1277041 | 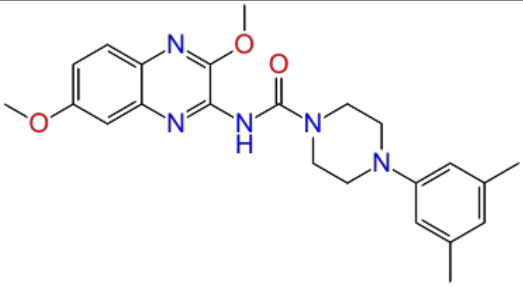 | 7.12 | train |
| 71. | CHEMBL1276866 | 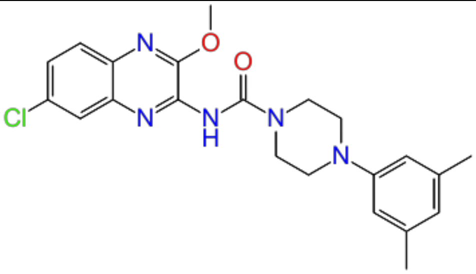 | 7.16 | test |
| 72. | CHEMBL1277939 | 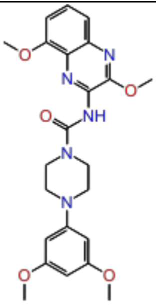 | 7.24 | train |
| 73. | CHEMBL1276865 | 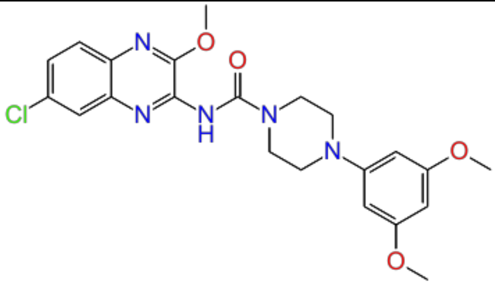 | 7.25 | train |

| | | | | |
|---|---|---|---|---|
| 74. | CHEMBL1278119 | 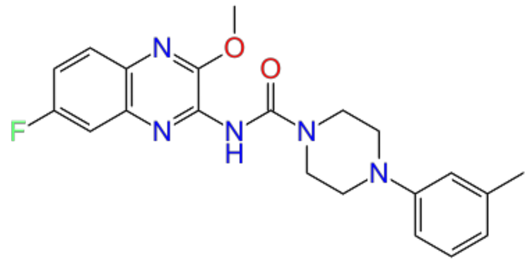 | 7.30 | train |
| 75. | CHEMBL1276953 | 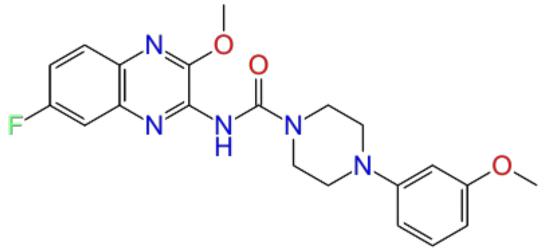 | 7.51 | test |
| 76. | CHEMBL3338195 | 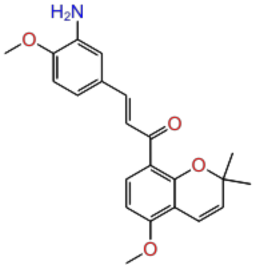 | 7.62 | train |
| 77. | CHEMBL428647 | 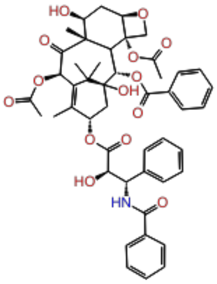 | 7.64 | train |
| 78. | CHEMBL1278205 | 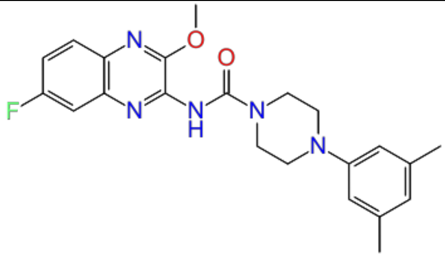 | 7.72 | train |

| | | | | |
|---|---|---|---|---|
| 79. | CHEMBL1278118 | 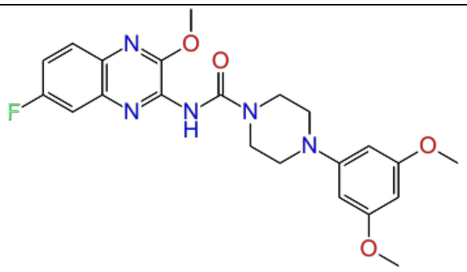 | 7.82 | test |
| 80. | CHEMBL1683544 | 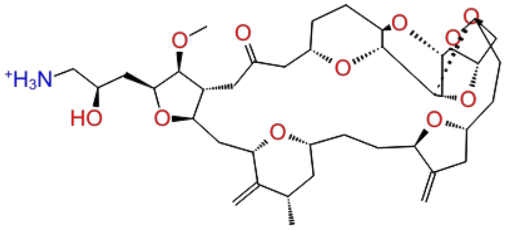 | 8.23 | train |
| 81. | CHEMBL1683553 | 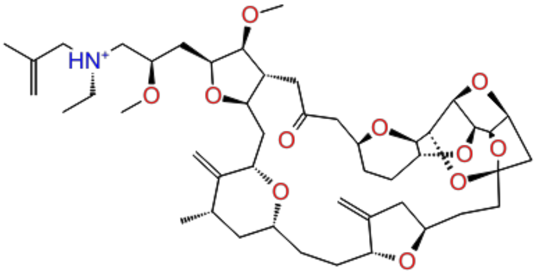 | 8.43 | test |
| 82. | CHEMBL1683552 | 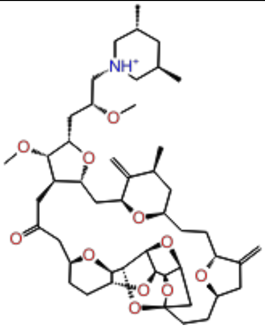 | 8.59 | train |
| 83. | CHEMBL2420629 | 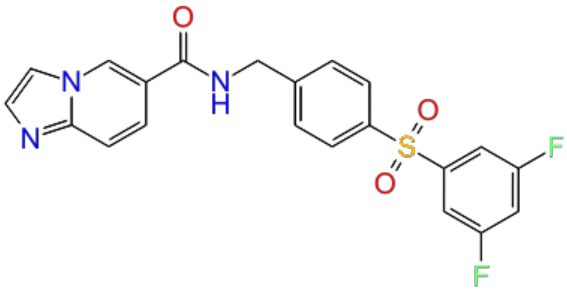 | 8.74 | train |

| | | | | |
|---|---|---|---|---|
| 84. | CHEMBL1683546 | 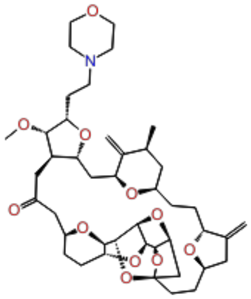 | 8.90 | train |
| 85. | CHEMBL1683523 | 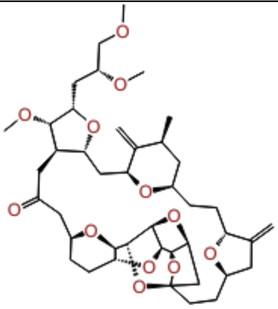 | 9.13 | test |
| 86. | CHEMBL1683549 | 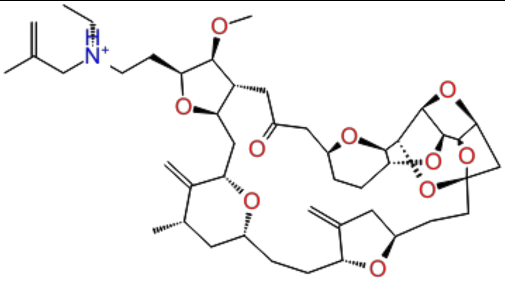 | 9.21 | train |
| 87. | CHEMBL1683550 | 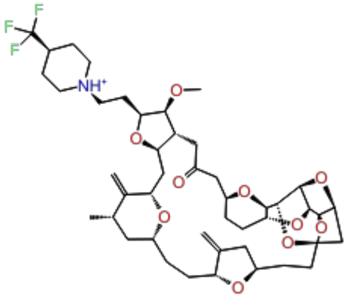 | 9.55 | train |
| 88. | CHEMBL1683556 | 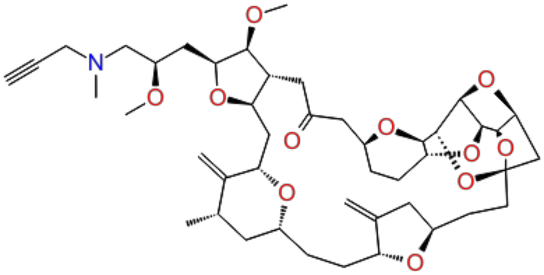 | 9.74 | train |

**Table S5:** AutoQSAR-U251 external test set from U251 cell line (56 compounds) along with $pIC_{50}$.

| Sr. No | Compound ID | 2D structure | $pIC_{50}$ |
|---|---|---|---|
| 1. | CHEMBL1958321 | 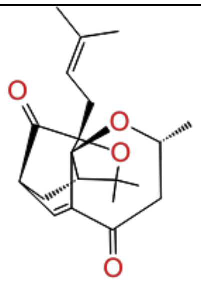 | 4.08 |
| 2. | CHEMBL252568 | 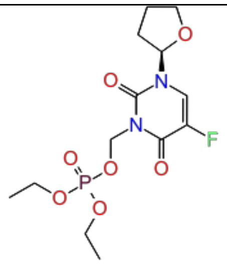 | 4.14 |
| 3. | CHEMBL452926 | 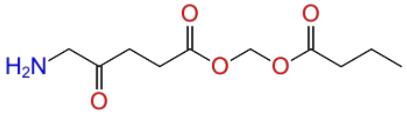 | 4.23 |
| 4. | CHEMBL467590 | 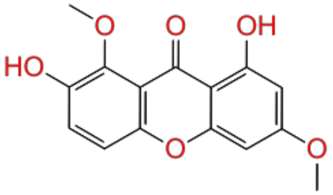 | 4.27 |
| 5. | CHEMBL3357146 | 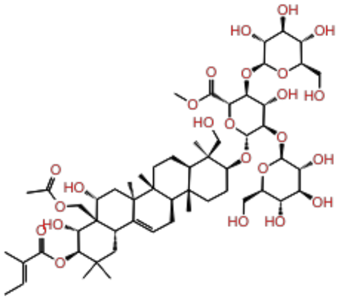 | 4.32 |

| | | | |
|---|---|---|---|
| 6. | CHEMBL364713 | 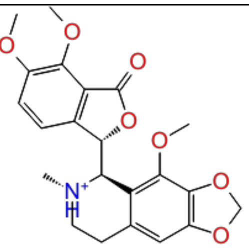 | 4.40 |
| 7. | CHEMBL3357167 | 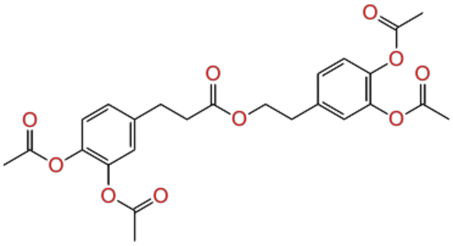 | 4.43 |
| 8. | CHEMBL45240 | 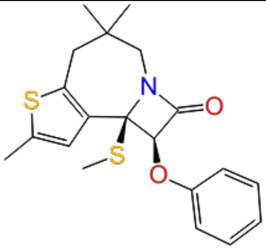 | 4.48 |
| 9. | CHEMBL1784163 | 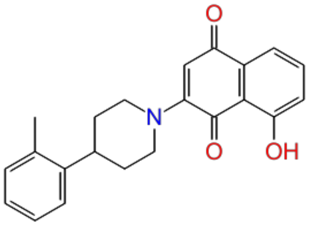 | 4.52 |
| 10. | CHEMBL178297 | 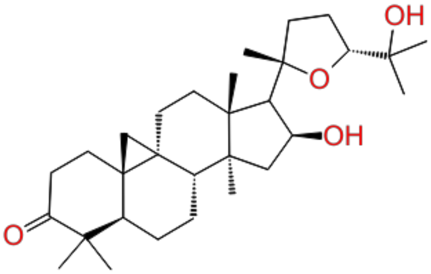 | 4.56 |
| 11. | CHEMBL3357166 | 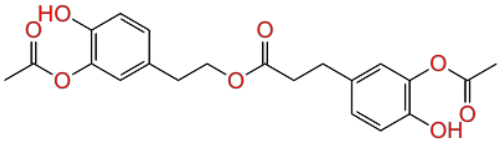 | 4.57 |

| | | | |
|---|---|---|---|
| 12. | CHEMBL1958316 | 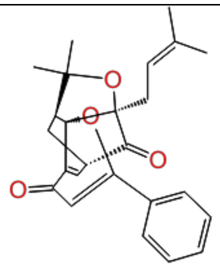 | 4.65 |
| 13. | CHEMBL180924 | 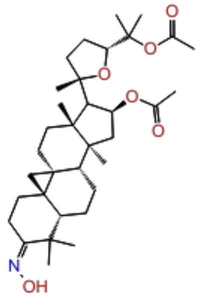 | 4.67 |
| 14. | CHEMBL179443 | 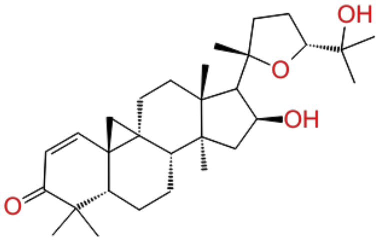 | 4.72 |
| 15. | CHEMBL3331345 | 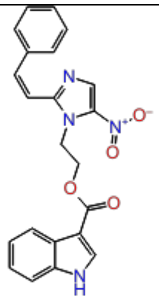 | 4.77 |
| 16. | CHEMBL3357147 | 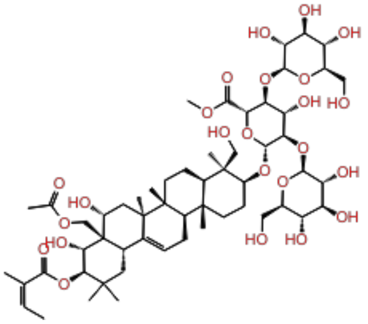 | 4.81 |

| | | | |
|---|---|---|---|
| 17. | CHEMBL178305 | 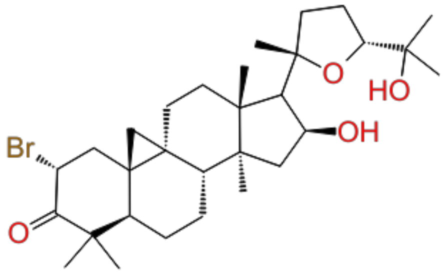 | 4.86 |
| 18. | CHEMBL3588937 | 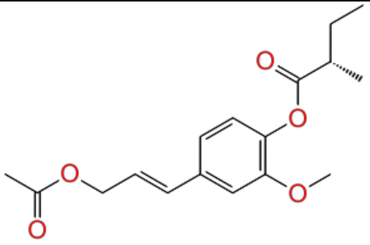 | 4.89 |
| 19. | CHEMBL1784159 | 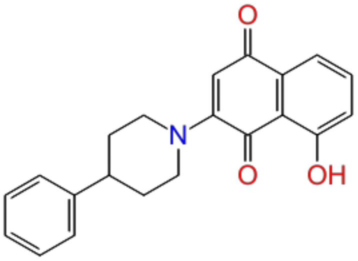 | 4.97 |
| 20. | CHEMBL478578 | 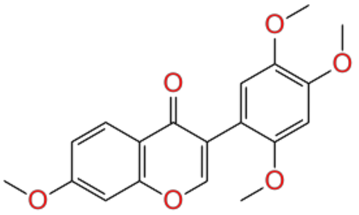 | 5.03 |
| 21. | CHEMBL1958320 | 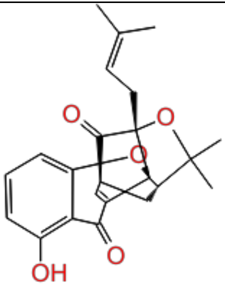 | 5.04 |

| | | | |
|---|---|---|---|
| 22. | CHEMBL1958314 | 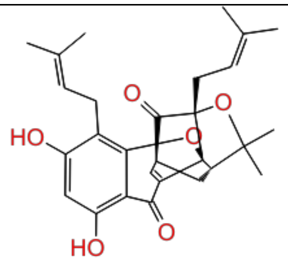 | 5.07 |
| 23. | CHEMBL4065827 | 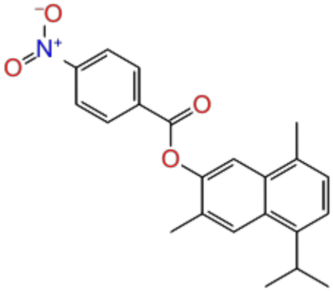 | 5.11 |
| 24. | CHEMBL498233 | 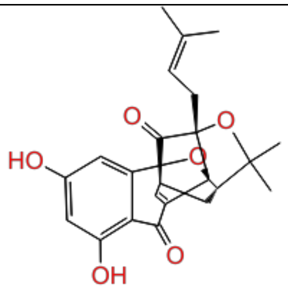 | 5.13 |
| 25. | CHEMBL1784157 | 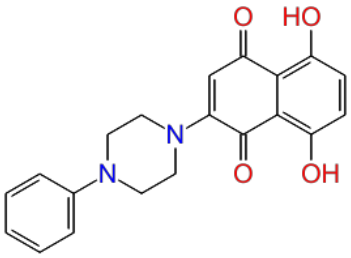 | 5.21 |
| 26. | CHEMBL3311053 | 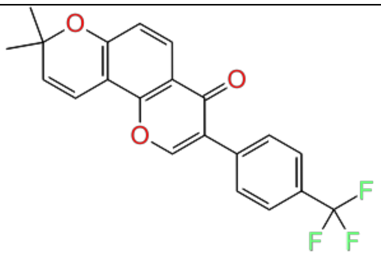 | 5.23 |

| | | | |
|---|---|---|---|
| 27. | CHEMBL1784155 | 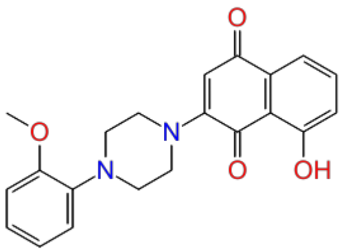 | 5.36 |
| 28. | CHEMBL3331008 | 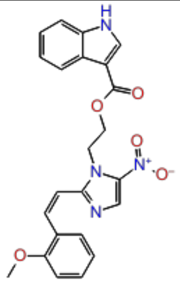 | 5.39 |
| 29. | CHEMBL3331000 | 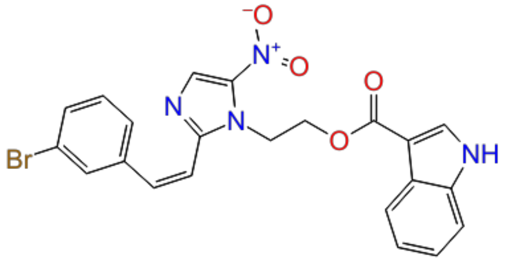 | 5.41 |
| 30. | CHEMBL3311165 | 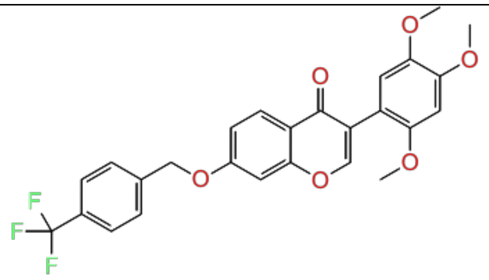 | 5.43 |
| 31. | CHEMBL3311159 | 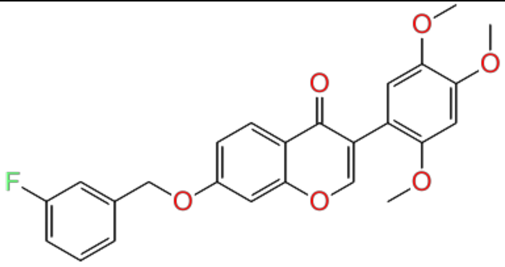 | 5.46 |

| | | | |
|---|---|---|---|
| 32. | CHEMBL3331007 | 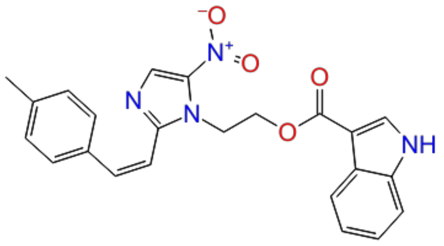 | 5.51 |
| 33. | CHEMBL3331006 | 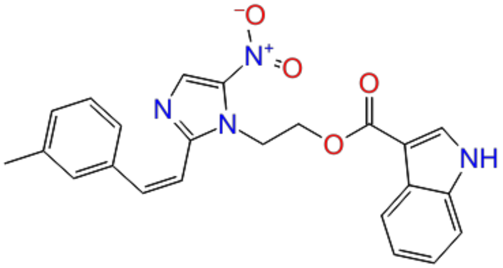 | 5.52 |
| 34. | CHEMBL3311150 | 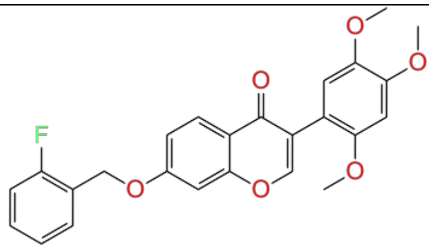 | 5.54 |
| 35. | CHEMBL3331005 | 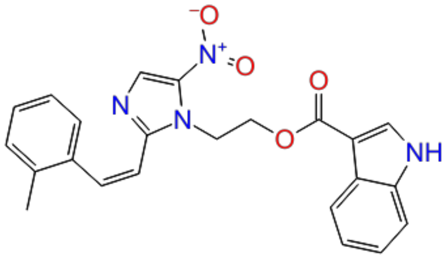 | 5.55 |
| 36. | CHEMBL3331009 | 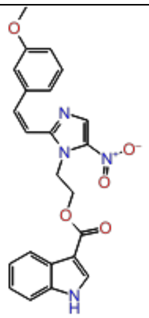 | 5.56 |

| | | | |
|---|---|---|---|
| 37. | CHEMBL555017 | 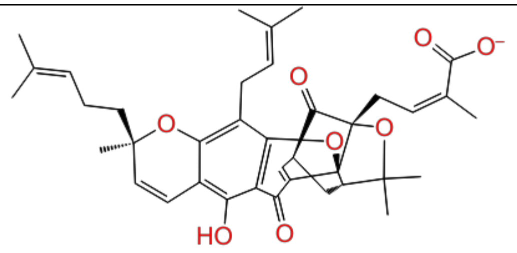 | 5.59 |
| 38. | CHEMBL3400386 | 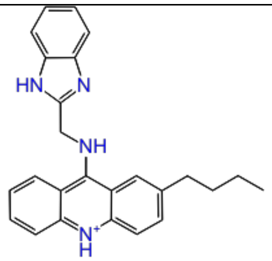 | 5.62 |
| 39. | CHEMBL3311153 | 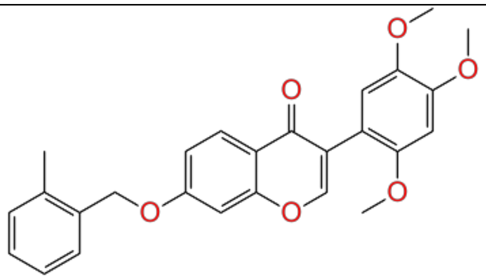 | 5.70 |
| 40. | CHEMBL61619 | 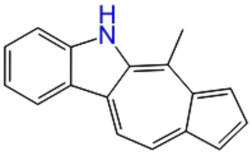 | 5.83 |
| 41. | CHEMBL3311158 | 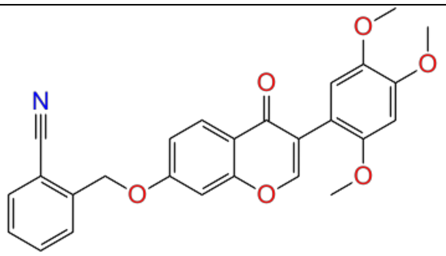 | 6.01 |

| | | | |
|---|---|---|---|
| 42. | CHEMBL2064455 | 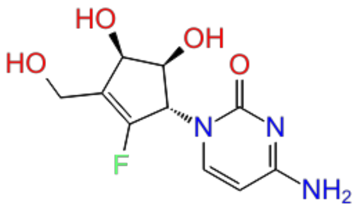 | 6.08 |
| 43. | CHEMBL3408446 | 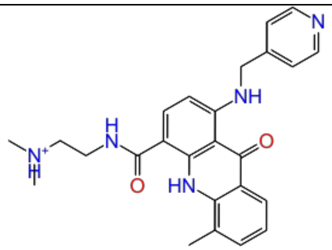 | 6.10 |
| 44. | CHEMBL67 | 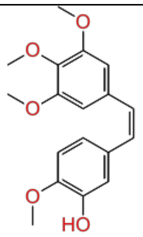 | 6.27 |
| 45. | CHEMBL4066486 | 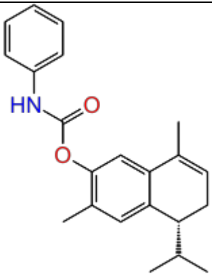 | 6.34 |
| 46. | CHEMBL1278206 | 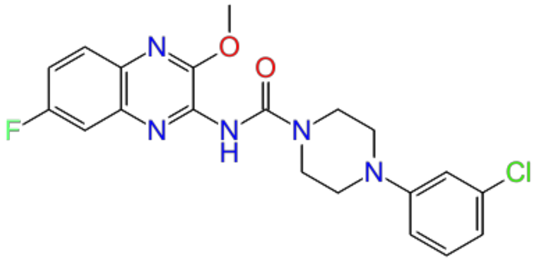 | 7.10 |

| | | | |
|---|---|---|---|
| 47. | CHEMBL1278025 | 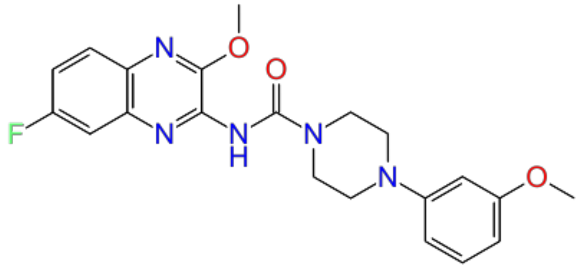 | 7.16 |
| 48. | CHEMBL1277938 | 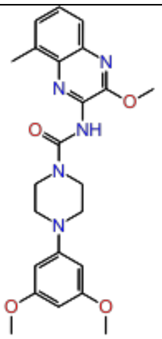 | 7.27 |
| 49. | CHEMBL1277042 | 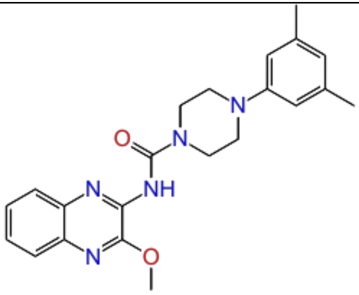 | 7.52 |
| 50. | CHEMBL107 | 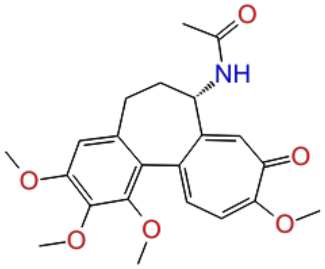 | 7.70 |
| 51. | CHEMBL1683547 | 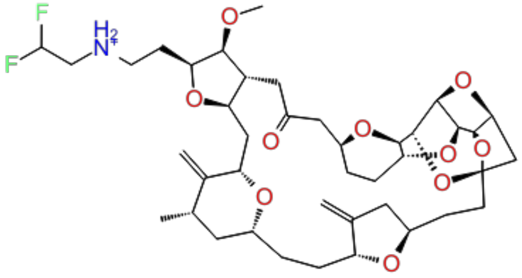 | 8.39 |

| | | | |
|---|---|---|---|
| 52. | CHEMBL1683554 | 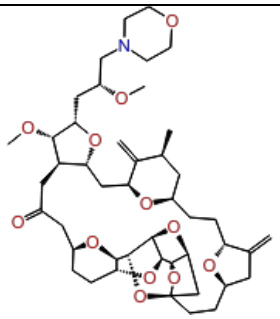 | 8.71 |
| 53. | CHEMBL1683548 | 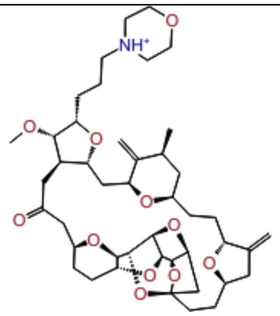 | 8.84 |
| 54. | CHEMBL1683551 | 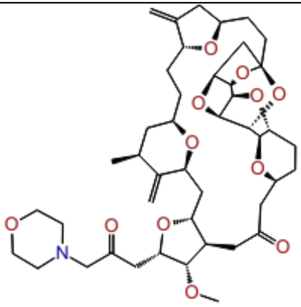 | 9.06 |
| 55. | CHEMBL1683545 | 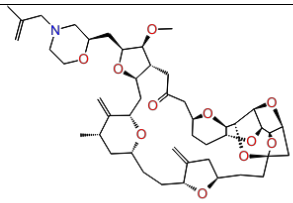 | 9.30 |
| 56. | CHEMBL1683555 | 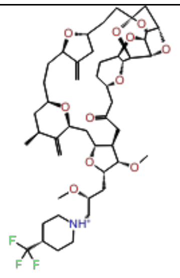 | 9.48 |

**Table S6:** Remaining 8 models constructed in ML-based U87 cell line models from U-87 inhibitors.

| Model # | Model code | Score | S.D. | $R^2$ | RMSE | $Q^2$ |
|---|---|---|---|---|---|---|
| 1 | kpls_radial_24 | 0.70 | 0.40 | 0.88 | 0.52 | 0.76 |
| 2 | kpls_dendritic_45 | 0.70 | 0.63 | 0.70 | 0.61 | 0.67 |
| 3 | kpls_linear_45 | 0.70 | 0.63 | 0.70 | 0.61 | 0.66 |
| 4 | kpls_radial_46 | 0.67 | 0.36 | 0.89 | 0.54 | 0.77 |
| 5 | kpls_dendtritic_40 | 0.66 | 0.66 | 0.66 | 0.64 | 0.64 |
| 6 | kpls_linear_40 | 0.66 | 0.67 | 0.66 | 0.65 | 0.63 |
| 7 | kpls_dendritic_24 | 0.66 | 0.43 | 0.86 | 0.56 | 0.72 |
| 8 | kpls_molprint2D_45 | 0.64 | 0.69 | 0.63 | 0.64 | 0.63 |

**Table S7:** Remaining 9 models constructed in ML-based U251 cell line models from U-251inhibitors.

| Model # | Model code | Score | S.D. | $R^2$ | RMSE | $Q^2$ |
|---|---|---|---|---|---|---|
| 1 | kpls_radial_16 | 0.88 | 0.53 | 0.88 | 0.48 | 0.87 |
| 2 | kpls_radial_36 | 0.88 | 0.51 | 0.89 | 0.50 | 0.87 |
| 3 | kpls_radial_24 | 0.87 | 0.55 | 0.87 | 0.41 | 0.91 |
| 4 | kpls_molprint2D_15 | 0.86 | 0.48 | 0.89 | 0.50 | 0.88 |
| 5 | kpls_molprint2D_16 | 0.83 | 0.62 | 0.83 | 0.52 | 0.84 |
| 6 | kpls_desc_16 | 0.81 | 0.68 | 0.81 | 0.60 | 0.80 |
| 7 | kpls_molprint2D_34 | 0.81 | 0.67 | 0.80 | 0.59 | 0.81 |
| 8 | kpls_linear_3 | 0.81 | 0.67 | 0.81 | 0.62 | 0.79 |
| 9 | kpls_radial_3 | 0.80 | 0.69 | 0.79 | 0.60 | 0.81 |

**Table S8:** ML-based models generated from IDH1 mutant inhibitors (training and test set) IDH1 ligands (173 compounds) along with $pIC_{50}$ values.

| Sr.No | Compound ID | 2D structure | $pIC_{50}$ |
|---|---|---|---|
| 1. | CHEMBL4217951 | 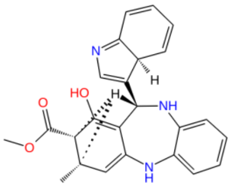 | 4.3 |
| 2. | CHEMBL4580009 | 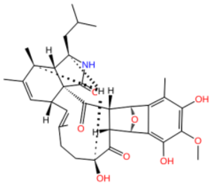 | 4.36 |
| 3. | CHEMBL167779 | 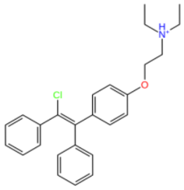 | 4.42 |
| 4. | CHEMBL4278116 | 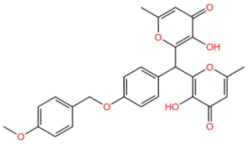 | 4.5 |
| 5. | CHEMBL4575450 | 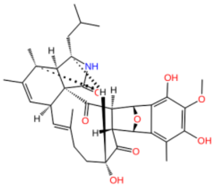 | 4.76 |

| | | | |
|---|---|---|---|
| 6. | CHEMBL4556155 | 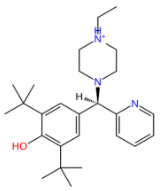 | 4.92 |
| 7. | CHEMBL4590233 | 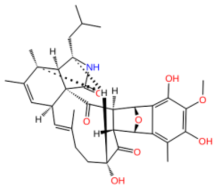 | 4.93 |
| 8. | CHEMBL4434758 | 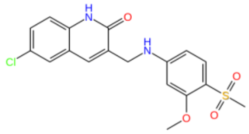 | 5.01 |
| 9. | CHEMBL3970770 | 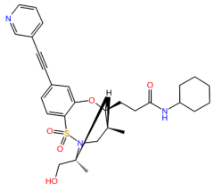 | 5.04 |
| 10. | CHEMBL3917452 | 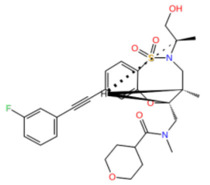 | 5.1 |

| | | | |
|---|---|---|---|
| 11. | CHEMBL2180743 | 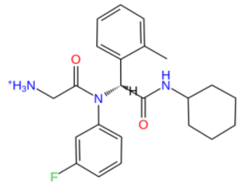 | 5.11 |
| 12. | CHEMBL3945329 | 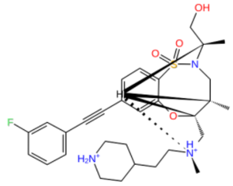 | 5.13 |
| 13. | CHEMBL2180397 | 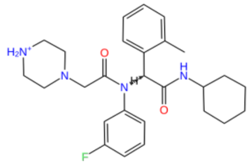 | 5.25 |
| 14. | CHEMBL4446593 | 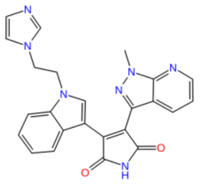 | 5.36 |
| 15. | CHEMBL4534551 | 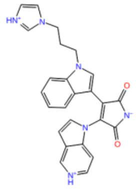 | 5.43 |

| | | | |
|---|---|---|---|
| 16. | CHEMBL2180751 | 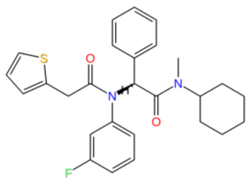 | 5.44 |
| 17. | CHEMBL4560003 | 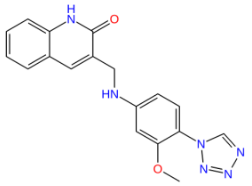 | 5.48 |
| 18. | CHEMBL4574211 | 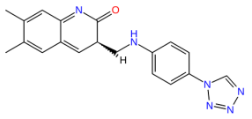 | 5.51 |
| 19. | CHEMBL4574726 | 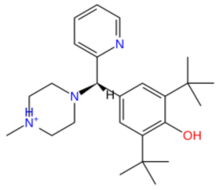 | 5.52 |
| 20. | CHEMBL4216167 | 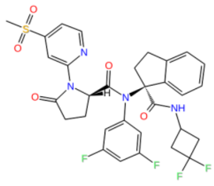 | 5.53 |

| | | | |
|---|---|---|---|
| 21. | CHEMBL2180733 | 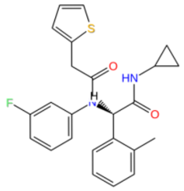 | 5.54 |
| 22. | CHEMBL4459022 | 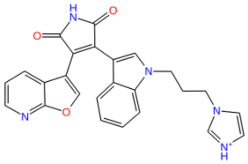 | 5.65 |
| 23. | CHEMBL2180735 | 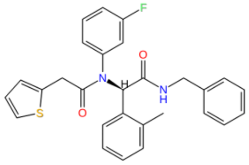 | 5.75 |
| 24. | CHEMBL4520403 | 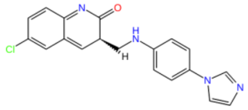 | 5.77 |
| 25. | CHEMBL2180395 | 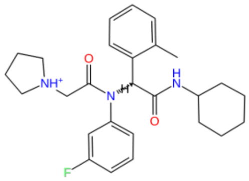 | 5.79 |

| | | | |
|---|---|---|---|
| 26. | CHEMBL3926431 | 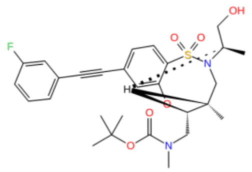 | 5.8 |
| 27. | CHEMBL3972520 | 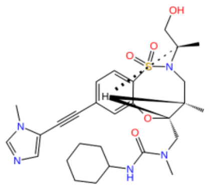 | 5.8 |
| 28. | CHEMBL4548634 | 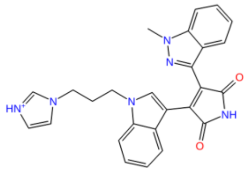 | 5.81 |
| 29. | CHEMBL4473155 | 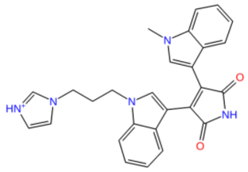 | 5.83 |
| 30. | CHEMBL4463255 | 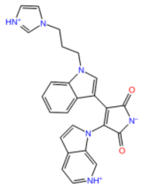 | 5.84 |

| | | | |
|---|---|---|---|
| 31. | CHEMBL2180750 | 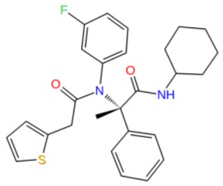 | 5.85 |
| 32. | CHEMBL3953491 | 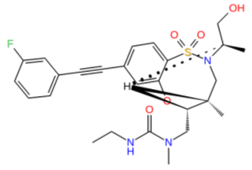 | 5.85 |
| 33. | CHEMBL550052 | 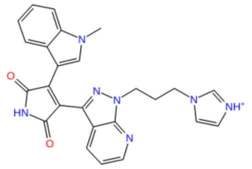 | 5.89 |
| 34. | CHEMBL2180741 | 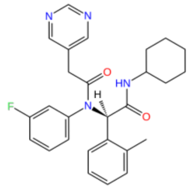 | 5.96 |
| 35. | CHEMBL4473004 | 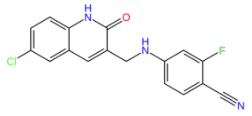 | 6 |

| | | | |
|---|---|---|---|
| 36. | CHEMBL3926888 | 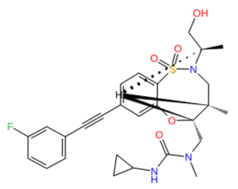 | 6 |
| 37. | CHEMBL4557985 | 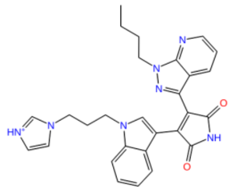 | 6.02 |
| 38. | CHEMBL4524313 | 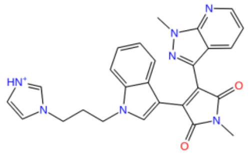 | 6.03 |
| 39. | CHEMBL2180742 | 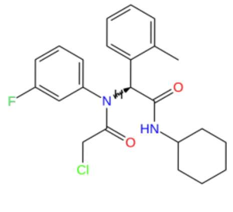 | 6.05 |
| 40. | CHEMBL2180732 | 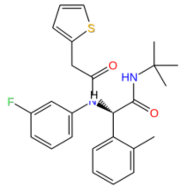 | 6.07 |

| | | | |
|---|---|---|---|
| 41. | CHEMBL4517014 | 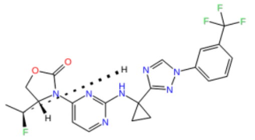 | 6.08 |
| 42. | CHEMBL3682033 | 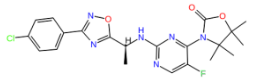 | 6.09 |
| 43. | CHEMBL4576620 | 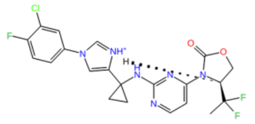 | 6.14 |
| 44. | CHEMBL4530068 | 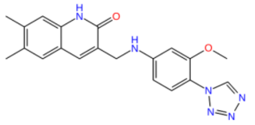 | 6.15 |
| 45. | CHEMBL4476486 | 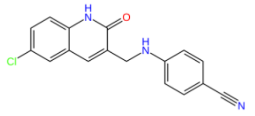 | 6.17 |

| | | | |
|---|---|---|---|
| 46. | CHEMBL2180736 | 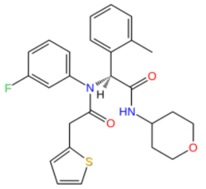 | 6.19 |
| 47. | CHEMBL4211712 | 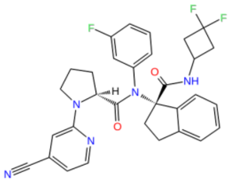 | 6.2 |
| 48. | CHEMBL3979949 | 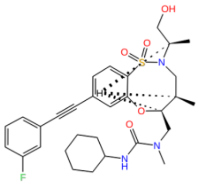 | 6.22 |
| 49. | CHEMBL2180734 | 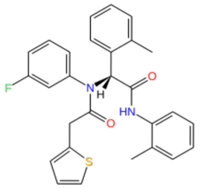 | 6.24 |
| 50. | CHEMBL3982213 | 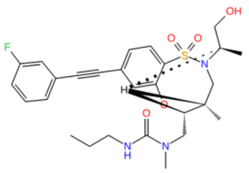 | 6.26 |

| | | | |
|---|---|---|---|
| 51. | CHEMBL4465347 | 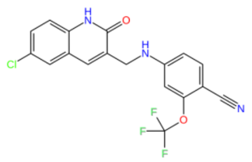 | 6.31 |
| 52. | CHEMBL4542800 | 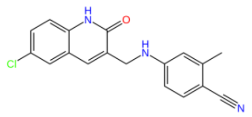 | 6.32 |
| 53. | CHEMBL4447496 | 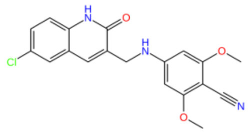 | 6.33 |
| 54. | CHEMBL4549366 | 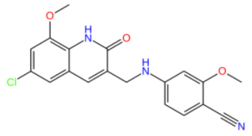 | 6.34 |
| 55. | CHEMBL2180745 | 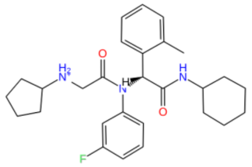 | 6.35 |

| | | | |
|---|---|---|---|
| 56. | CHEMBL3962802 | 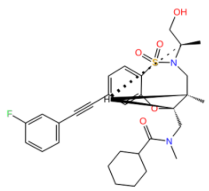 | 6.36 |
| 57. | CHEMBL2180399 | 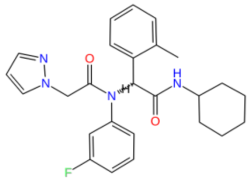 | 6.38 |
| 58. | CHEMBL2095003 | 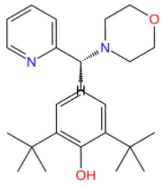 | 6.39 |
| 59. | CHEMBL4550645 | 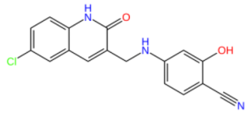 | 6.42 |
| 60. | CHEMBL4520492 | 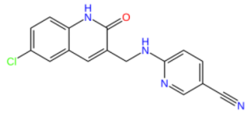 | 6.44 |

| | | | |
|---|---|---|---|
| 61. | CHEMBL4438247 | 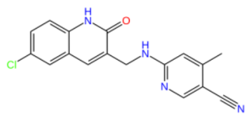 | 6.46 |
| 62. | CHEMBL4203130 | 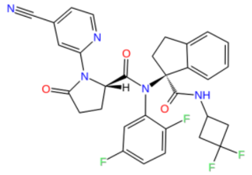 | 6.47 |
| 63. | CHEMBL3891327 | 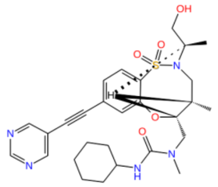 | 6.5 |
| 64. | CHEMBL4547765 | 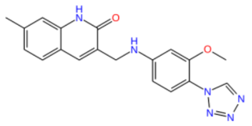 | 6.52 |
| 65. | CHEMBL4444111 | 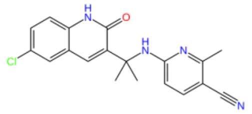 | 6.53 |

| | | | |
|---|---|---|---|
| 66. | CHEMBL4457293 | 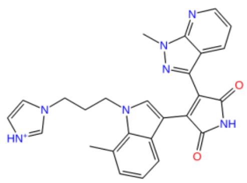 | 6.54 |
| 67. | CHEMBL4215610 | 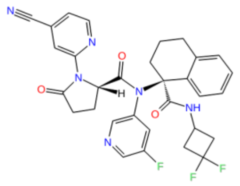 | 6.56 |
| 68. | CHEMBL4203416 | 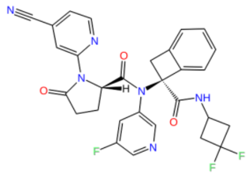 | 6.57 |
| 69. | CHEMBL4216287 | 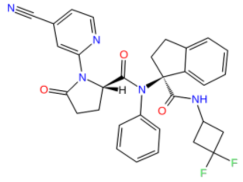 | 6.58 |
| 70. | CHEMBL4561231 | 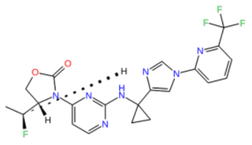 | 6.58 |

| | | | |
|---|---|---|---|
| 71. | CHEMBL4214783 | 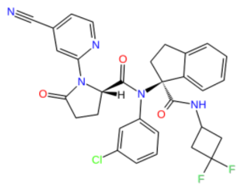 | 6.59 |
| 72. | CHEMBL4483760 | 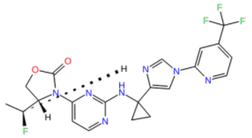 | 6.6 |
| 73. | CHEMBL4566994 | 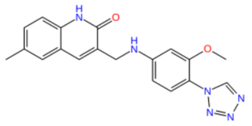 | 6.62 |
| 74. | CHEMBL2180396 | 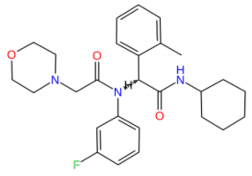 | 6.62 |
| 75. | CHEMBL4453798 | 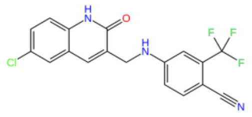 | 6.64 |

| | | | |
|---|---|---|---|
| 76. | CHEMBL4544382 | 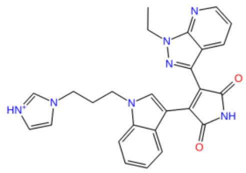 | 6.64 |
| 77. | CHEMBL4588007 | 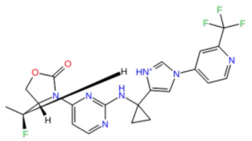 | 6.66 |
| 78. | CHEMBL4467855 | 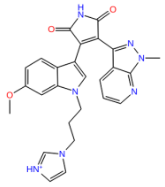 | 6.72 |
| 79. | CHEMBL3939537 | 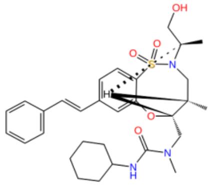 | 6.72 |
| 80. | CHEMBL4465669 | 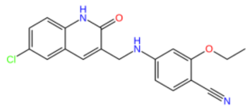 | 6.73 |

| | | | |
|---|---|---|---|
| 81. | CHEMBL4215069 | 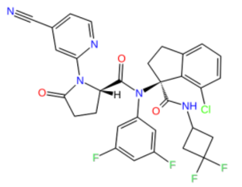 | 6.75 |
| 82. | CHEMBL4212804 | 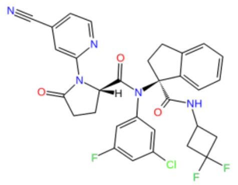 | 6.76 |
| 83. | CHEMBL3944296 | 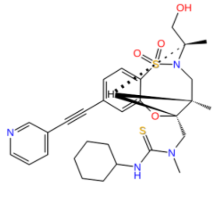 | 6.8 |
| 84. | CHEMBL3974203 | 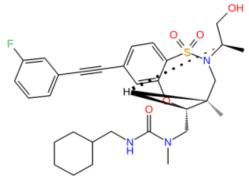 | 6.8 |
| 85. | CHEMBL4558914 | 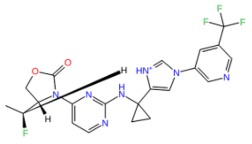 | 6.82 |

| | | | |
|---|---|---|---|
| 86. | CHEMBL4551306 | 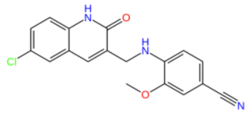 | 6.83 |
| 87. | CHEMBL4514241 | 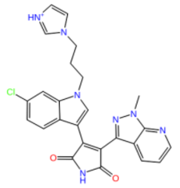 | 6.85 |
| 88. | CHEMBL3917616 | 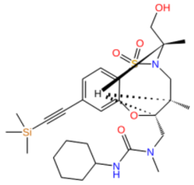 | 6.85 |
| 89. | CHEMBL2180400 | 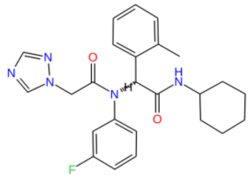 | 6.85 |
| 90. | CHEMBL4215481 | 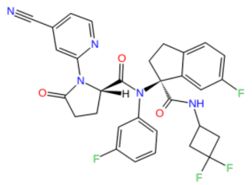 | 6.86 |

| | | | |
|---|---|---|---|
| 91. | CHEMBL4546161 | 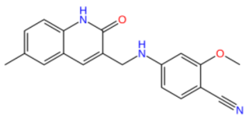 | 6.88 |
| 92. | CHEMBL4203955 | 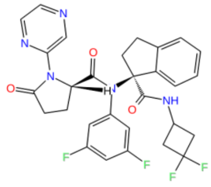 | 6.89 |
| 93. | CHEMBL4441769 | 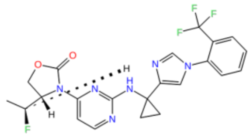 | 6.95 |
| 94. | CHEMBL4203806 | 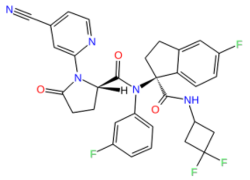 | 6.97 |
| 95. | CHEMBL4469450 | 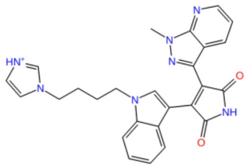 | 7 |

| | | | |
|---|---|---|---|
| 96. | CHEMBL3950295 | 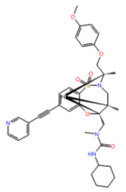 | 7.02 |
| 97. | CHEMBL4216343 | 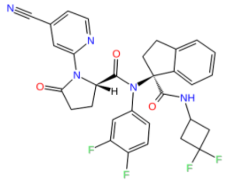 | 7.02 |
| 98. | CHEMBL4215084 | 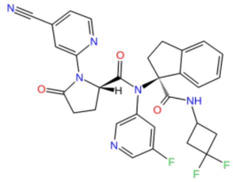 | 7.03 |
| 99. | CHEMBL3960382 | 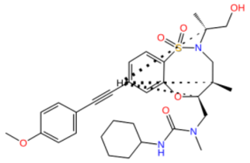 | 7.04 |
| 100. | CHEMBL3972787 | 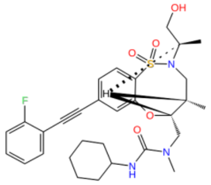 | 7.04 |

| | | | |
|---|---|---|---|
| 101. | CHEMBL4436439 | 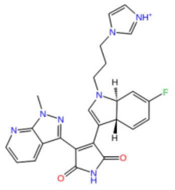 | 7.05 |
| 102. | CHEMBL4210654 | 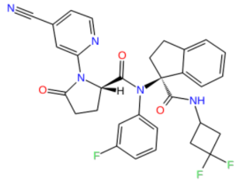 | 7.07 |
| 103. | CHEMBL4483502 | 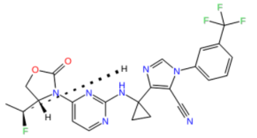 | 7.07 |
| 104. | CHEMBL4456915 | 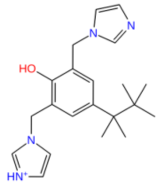 | 7.09 |
| 105. | CHEMBL2180394 | 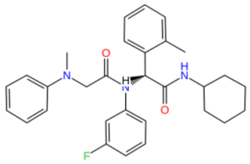 | 7.1 |

| | | | |
|---|---|---|---|
| 106. | CHEMBL4212118 | 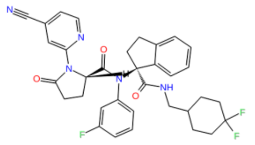 | 7.1 |
| 107. | CHEMBL4463451 | 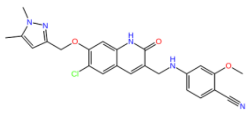 | 7.11 |
| 108. | CHEMBL4289678 | 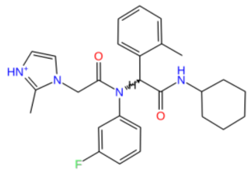 | 7.16 |
| 109. | CHEMBL4211065 | 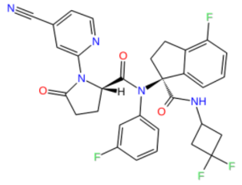 | 7.17 |
| 110. | CHEMBL4468093 | 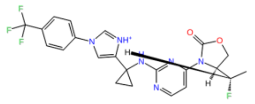 | 7.19 |

| | | | |
|---|---|---|---|
| 111. | CHEMBL4476190 | 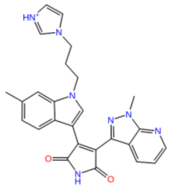 | 7.22 |
| 112. | CHEMBL4556914 | 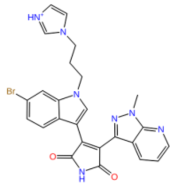 | 7.22 |
| 113. | CHEMBL4207520 | 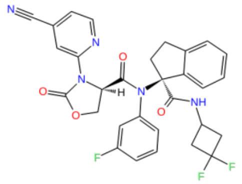 | 7.24 |
| 114. | CHEMBL3960589 | 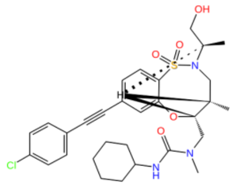 | 7.25 |
| 115. | CHEMBL4541453 | 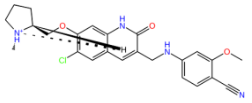 | 7.25 |

| | | | |
|---|---|---|---|
| 116. | CHEMBL4210785 | 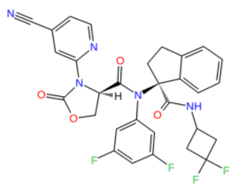 | 7.27 |
| 117. | CHEMBL4442007 | 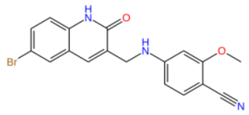 | 7.27 |
| 118. | CHEMBL4207639 | 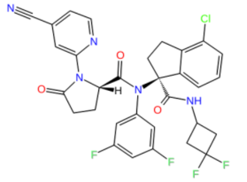 | 7.28 |
| 119. | CHEMBL4541801 | 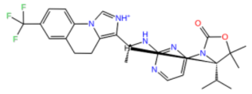 | 7.29 |
| 120. | CHEMBL4212544 | 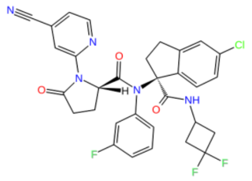 | 7.31 |

| | | | |
|---|---|---|---|
| 121. | CHEMBL4217038 | 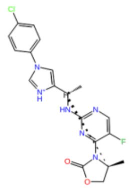 | 7.33 |
| 122. | CHEMBL4282455 | 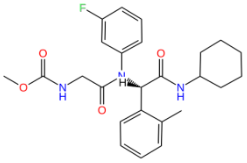 | 7.34 |
| 123. | CHEMBL4285677 | 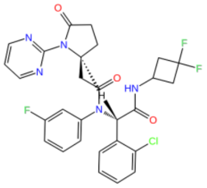 | 7.35 |
| 124. | CHEMBL4207897 | 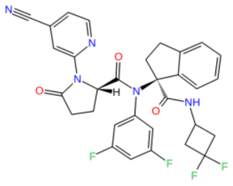 | 7.35 |
| 125. | CHEMBL4285854 | 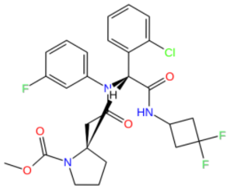 | 7.36 |

| | | | |
|---|---|---|---|
| 126. | CHEMBL4537907 | 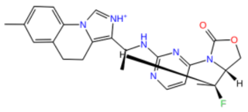 | 7.37 |
| 127. | CHEMBL4444615 | 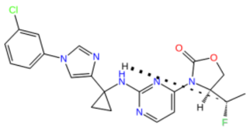 | 7.38 |
| 128. | CHEMBL4215739 | 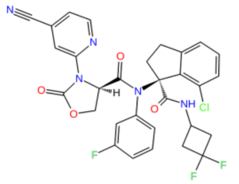 | 7.38 |
| 129. | CHEMBL4555124 | 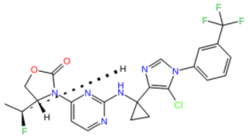 | 7.39 |
| 130. | CHEMBL4444255 | 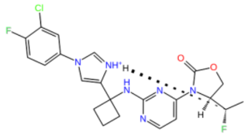 | 7.4 |

| | | | |
|---|---|---|---|
| 131. | CHEMBL4216196 | 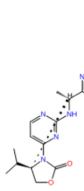 | 7.41 |
| 132. | CHEMBL2359350 | 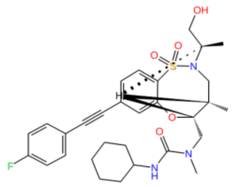 | 7.43 |
| 133. | CHEMBL4214694 | 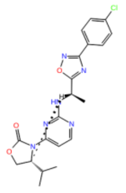 | 7.44 |
| 134. | CHEMBL4475577 | 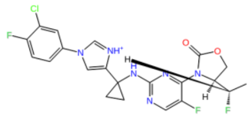 | 7.46 |
| 135. | CHEMBL4567884 | 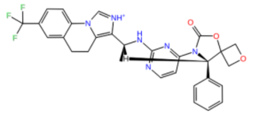 | 7.47 |

| | | | |
|---|---|---|---|
| 136. | CHEMBL4445771 | 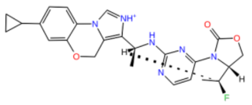 | 7.48 |
| 137. | CHEMBL4455045 | 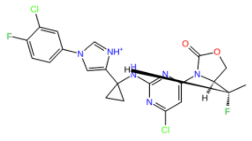 | 7.5 |
| 138. | CHEMBL4446109 | 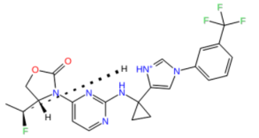 | 7.5 |
| 139. | CHEMBL3923861 | 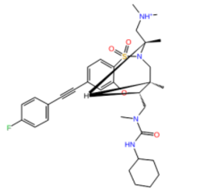 | 7.5 |
| 140. | CHEMBL3953686 | 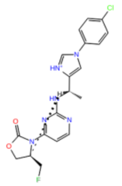 | 7.52 |

| | | | |
|---|---|---|---|
| 141. | CHEMBL3987085 | 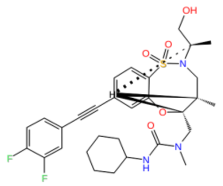 | 7.55 |
| 142. | CHEMBL4469739 | 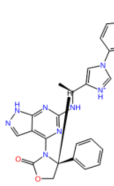 | 7.55 |
| 143. | CHEMBL4545655 | 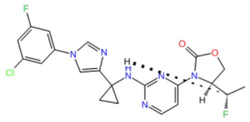 | 7.57 |
| 144. | CHEMBL4440585 | 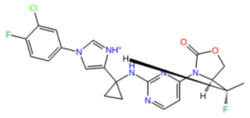 | 7.6 |
| 145. | CHEMBL3894881 | 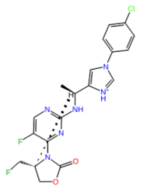 | 7.62 |

| | | | |
|---|---|---|---|
| 146. | CHEMBL4553096 | 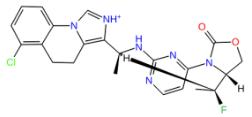 | 7.62 |
| 147. | CHEMBL4455559 | 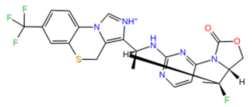 | 7.62 |
| 148. | CHEMBL4563967 | 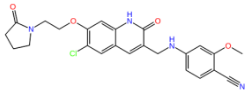 | 7.64 |
| 149. | CHEMBL3959391 | 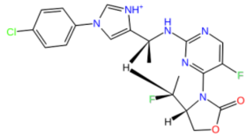 | 7.66 |
| 150. | CHEMBL4216785 | 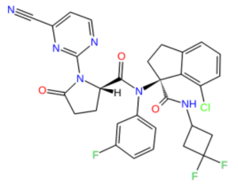 | 7.66 |

| | | | |
|---|---|---|---|
| 151. | CHEMBL4204501 | 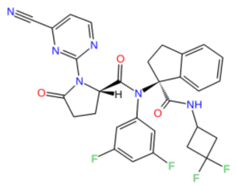 | 7.66 |
| 152. | CHEMBL4476025 | 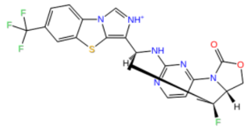 | 7.72 |
| 153. | CHEMBL4474186 | 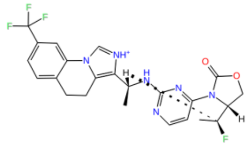 | 7.74 |
| 154. | CHEMBL4437186 | 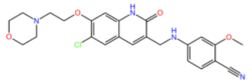 | 7.8 |
| 155. | CHEMBL4451801 | 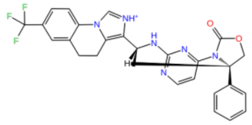 | 7.81 |

| | | | |
|---|---|---|---|
| 156. | CHEMBL4208648 | 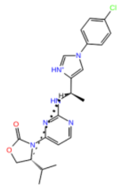 | 7.82 |
| 157. | CHEMBL4216510 | 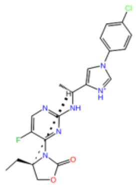 | 7.89 |
| 158. | CHEMBL4282267 | 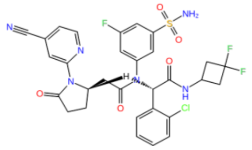 | 7.96 |
| 159. | CHEMBL4464313 | 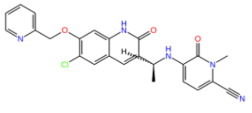 | 7.96 |
| 160. | CHEMBL4278793 | 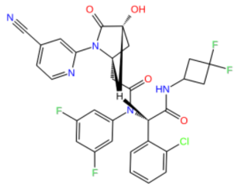 | 8 |

| | | | |
|---|---|---|---|
| 161. | CHEMBL4289061 | 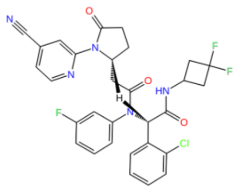 | 8.05 |
| 162. | CHEMBL4469055 | 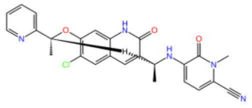 | 8.1 |
| 163. | CHEMBL4289692 | 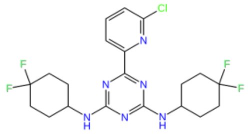 | 8.15 |
| 164. | CHEMBL4214223 | 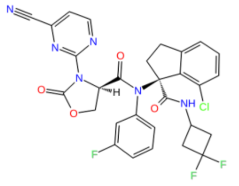 | 8.25 |
| 165. | CHEMBL4568616 | 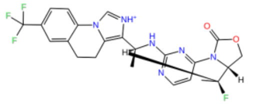 | 8.4 |

| | | | |
|---|---|---|---|
| 166. | CHEMBL4288840 | 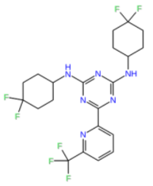 | 8.4 |
| 167. | CHEMBL4475715 | 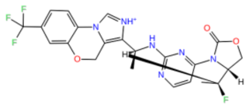 | 8.57 |
| 168. | CHEMBL4467991 | 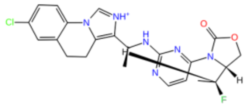 | 8.64 |
| 169. | CHEMBL4529476 | 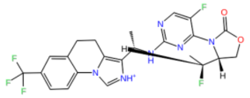 | 8.68 |
| 170. | CHEMBL4439421 | 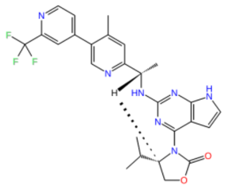 | 8.7 |

| | | | |
|---|---|---|---|
| 171. | CHEMBL4280772 | 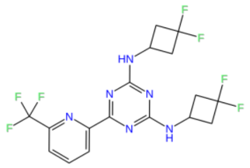 | 9 |
| 172. | CHEMBL4444067 | 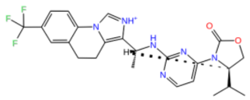 | 9 |
| 173. | CHEMBL4278845 | 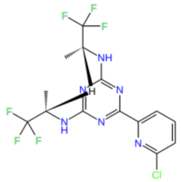 | 9.6 |

**Table S9:** External test set compounds (85 compounds) and their $pIC_{50}$ values used in the validation of ML-models generated from IDH1 mutant inhibitors.

| Sr.No | Compound ID | 2D structure | $pIC_{50}$ |
|---|---|---|---|
| 1. | CHEMBL4450414 | 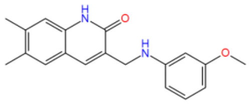 | 4.29 |
| 2. | CHEMBL4579725 | 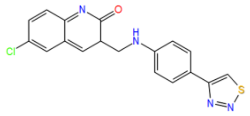 | 4.34 |

| | | | |
|---|---|---|---|
| 3. | CHEMBL4286032 | 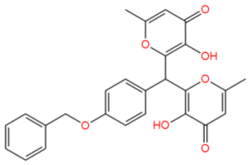 | 4.64 |
| 4. | CHEMBL3895369 | 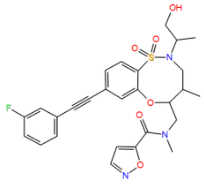 | 4.72 |
| 5. | CHEMBL4562901 | 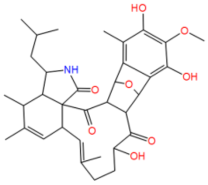 | 4.88 |
| 6. | CHEMBL4453488 | 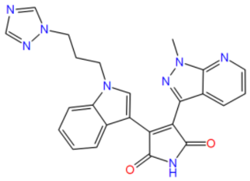 | 4.93 |
| 7. | CHEMBL4467139 | 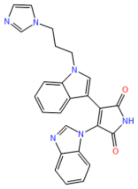 | 5.1 |

| | | | |
|---|---|---|---|
| 8. | CHEMBL3979015 | 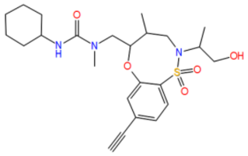 | 5.11 |
| 9. | CHEMBL4469507 | 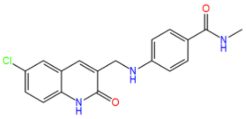 | 5.2 |
| 10. | CHEMBL3973763 | 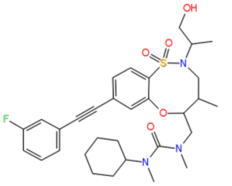 | 5.24 |
| 11. | CHEMBL4446452 | 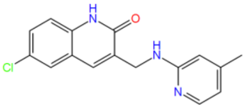 | 5.28 |
| 12. | CHEMBL4451545 | 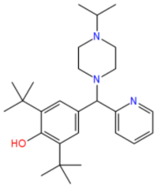 | 5.32 |

| | | | |
|---|---|---|---|
| 13. | CHEMBL3976839 | 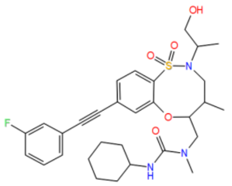 | 5.41 |
| 14. | CHEMBL4471075 | 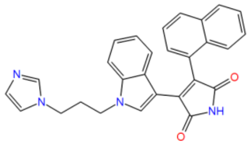 | 5.52 |
| 15. | CHEMBL4207353 | 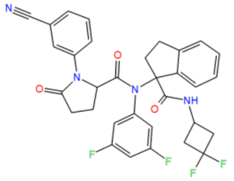 | 5.58 |
| 16. | CHEMBL4442650 | 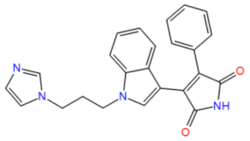 | 5.7 |
| 17. | CHEMBL3950996 | 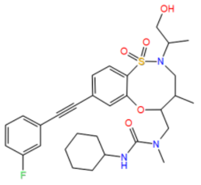 | 5.77 |

| | | | |
|---|---|---|---|
| 18. | CHEMBL4560109 | 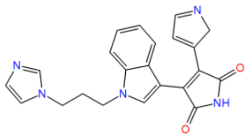 | 5.8 |
| 19. | CHEMBL4454383 | 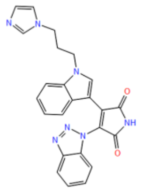 | 5.85 |
| 20. | CHEMBL4449515 | 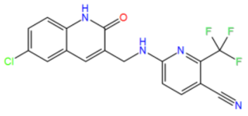 | 5.9 |
| 21. | CHEMBL4444846 | 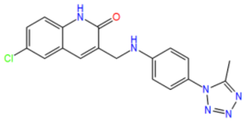 | 6 |
| 22. | CHEMBL2357019 | 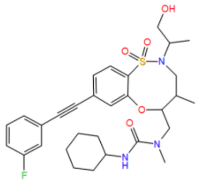 | 6 |

| | | | |
|---|---|---|---|
| 23. | CHEMBL4568252 | 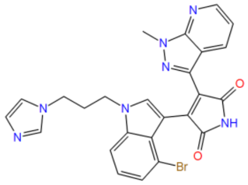 | 6.03 |
| 24. | CHEMBL4513764 | 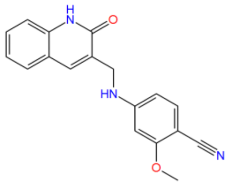 | 6.04 |
| 25. | CHEMBL4471506 | 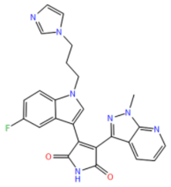 | 6.09 |
| 26. | CHEMBL4207387 | 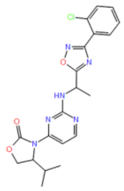 | 6.21 |
| 27. | CHEMBL4212061 | 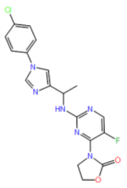 | 6.25 |

| | | | |
|---|---|---|---|
| 28. | CHEMBL4575435 | 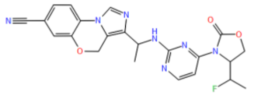 | 6.32 |
| 29. | CHEMBL3935326 | 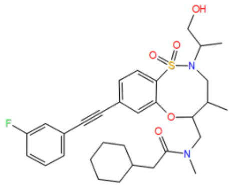 | 6.44 |
| 30. | CHEMBL4460692 | 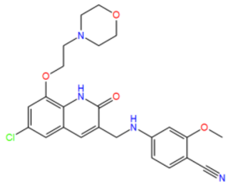 | 6.47 |
| 31. | CHEMBL2180731 | 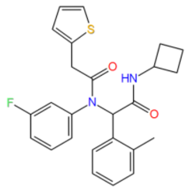 | 6.54 |
| 32. | CHEMBL4558677 | 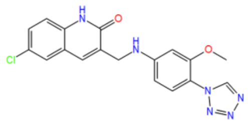 | 6.57 |

| | | | |
|---|---|---|---|
| 33. | CHEMBL3904242 | 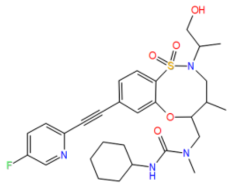 | 6.58 |
| 34. | CHEMBL4214891 | 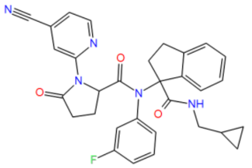 | 6.58 |
| 35. | CHEMBL4114410 | 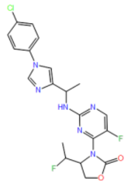 | 6.62 |
| 36. | CHEMBL4293065 | 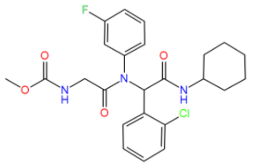 | 6.64 |
| 37. | CHEMBL4547484 | 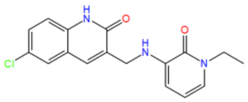 | 6.65 |

| | | | |
|---|---|---|---|
| 38. | CHEMBL2180744 | 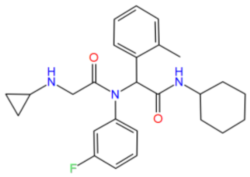 | 6.72 |
| 39. | CHEMBL4216027 | 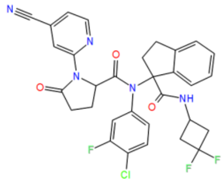 | 6.73 |
| 40. | CHEMBL4461178 | 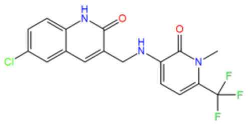 | 6.8 |
| 41. | CHEMBL4442250 | 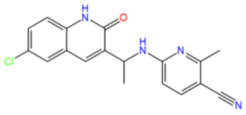 | 6.83 |
| 42. | CHEMBL4202591 | 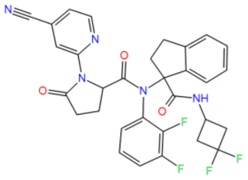 | 6.83 |

| | | | |
|---|---|---|---|
| 43. | CHEMBL4568861 | 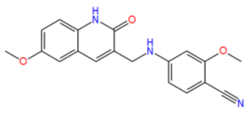 | 6.85 |
| 44. | CHEMBL3926310 | 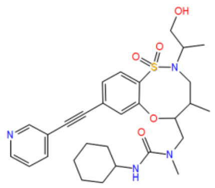 | 6.85 |
| 45. | CHEMBL4435874 | 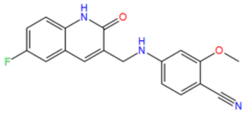 | 6.86 |
| 46. | CHEMBL4532585 | 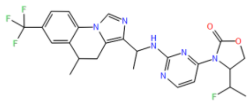 | 6.89 |
| 47. | CHEMBL4461972 | 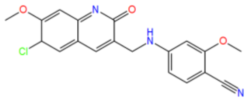 | 6.9 |

| | | | |
|---|---|---|---|
| 48. | CHEMBL4440134 | 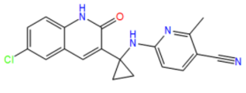 | 7 |
| 49. | CHEMBL4465287 | 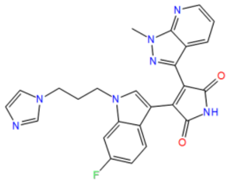 | 7.05 |
| 50. | CHEMBL3891685 | 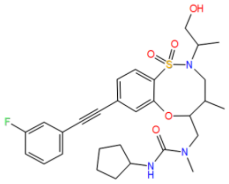 | 7.1 |
| 51. | CHEMBL4448857 | 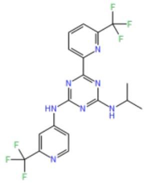 | 7.11 |
| 52. | CHEMBL4520104 | 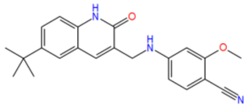 | 7.12 |

| | | | |
|---|---|---|---|
| 53. | CHEMBL4445691 | 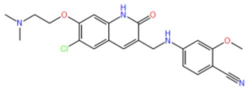 | 7.16 |
| 54. | CHEMBL4203298 | 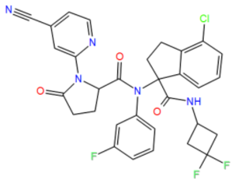 | 7.17 |
| 55. | CHEMBL4470532 | 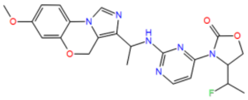 | 7.21 |
| 56. | CHEMBL2180748 | 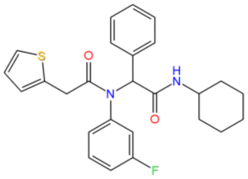 | 7.22 |
| 57. | CHEMBL3909367 | 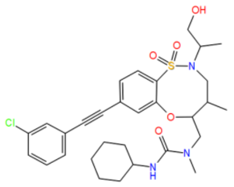 | 7.22 |

| | | | |
|---|---|---|---|
| 58. | CHEMBL4460895 | 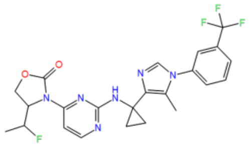 | 7.25 |
| 59. | CHEMBL4471133 | 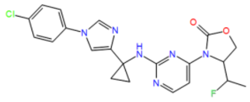 | 7.27 |
| 60. | CHEMBL4513760 | 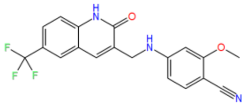 | 7.28 |
| 61. | CHEMBL4208169 | 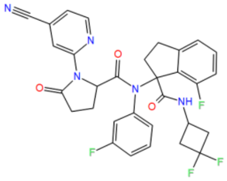 | 7.29 |
| 62. | CHEMBL3947537 | 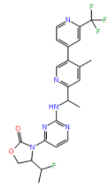 | 7.3 |

| | | | |
|---|---|---|---|
| 63. | CHEMBL4546041 | 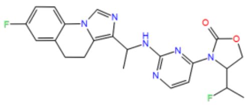 | 7.32 |
| 64. | CHEMBL4592898 | 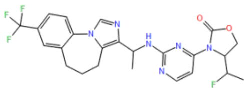 | 7.35 |
| 65. | CHEMBL4570265 | 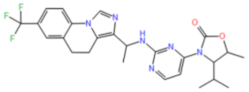 | 7.38 |
| 66. | CHEMBL4547257 | 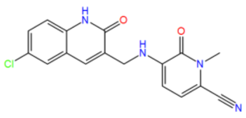 | 7.39 |
| 67. | CHEMBL4281172 | 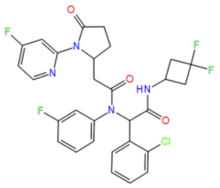 | 7.4 |

| | | | |
|---|---|---|---|
| 68. | CHEMBL4202711 | 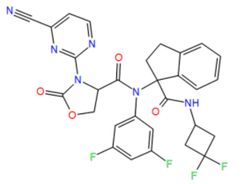 | 7.4 |
| 69. | CHEMBL4206945 | 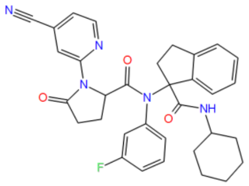 | 7.43 |
| 70. | CHEMBL4466857 | 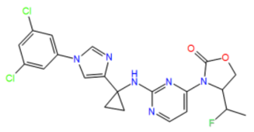 | 7.44 |
| 71. | CHEMBL4277931 | 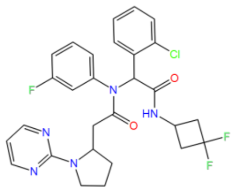 | 7.47 |
| 72. | CHEMBL4515664 | 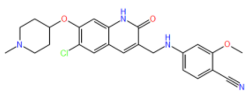 | 7.5 |

| | | | |
|---|---|---|---|
| 73. | CHEMBL4449563 | 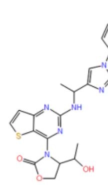 | 7.55 |
| 74. | CHEMBL4476345 | 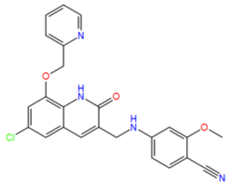 | 7.66 |
| 75. | CHEMBL3909586 | 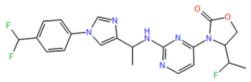 | 7.82 |
| 76. | CHEMBL4567882 | 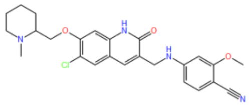 | 7.89 |
| 77. | CHEMBL4586919 | 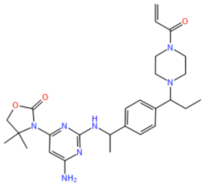 | 7.96 |

| | | | |
|---|---|---|---|
| 78. | CHEMBL4292864 | 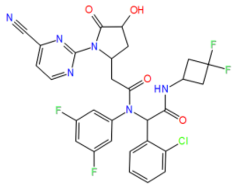 | 8.15 |
| 79. | CHEMBL4289465 | 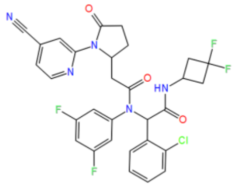 | 8.22 |
| 80. | CHEMBL4170686 | 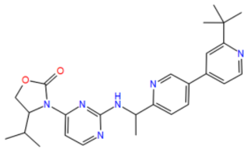 | 8.4 |
| 81. | CHEMBL4282989 | 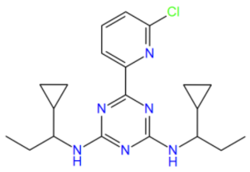 | 8.4 |
| 82. | CHEMBL4548637 | 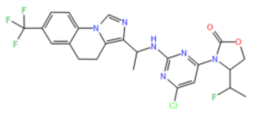 | 8.67 |

| | | | |
|---|---|---|---|
| 83. | CHEMBL4463644 | 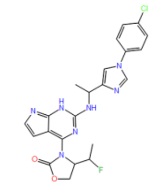 | 8.7 |
| 84. | CHEMBL4277352 | 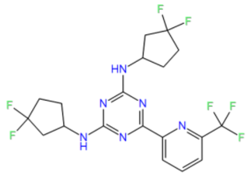 | 9 |
| 85. | CHEMBL4283785 | 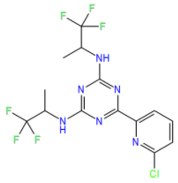 | 9.52 |

**Table S10:** Dataset of inhibitors against U87 cell line for pharmacophore hypothesis and 3D QSAR model generation (PHASE U87)

| Sr. No | Compound ID | 2D structure | $pIC_{50}$ | Dataset Division (Train & Test) |
|---|---|---|---|---|
| 1. | CHEMBL266090 | 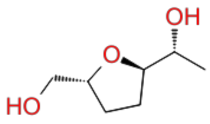 | 3.60 | test |
| 2. | CHEMBL7701 | 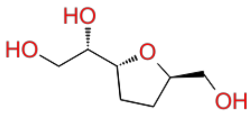 | 3.60 | train |
| 3. | CHEMBL589122 | 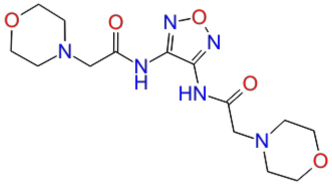 | 3.70 | test |

| | | | | |
|---|---|---|---|---|
| 4. | CHEMBL590093 | 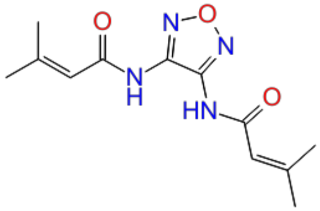 | 3.70 | train |
| 5. | CHEMBL600046 | 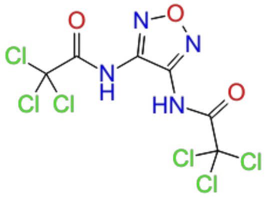 | 3.70 | train |
| 6. | CHEMBL4293736 | 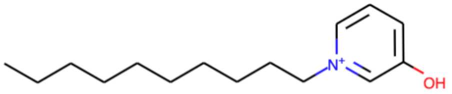 | 3.80 | train |
| 7. | CHEMBL590720 | 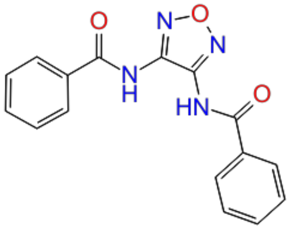 | 3.92 | Test |
| 8. | CHEMBL7476 | 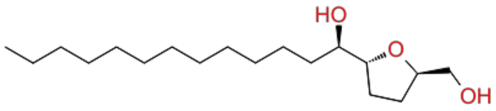 | 3.97 | Train |

| | | | | |
|---|---|---|---|---|
| 9. | CHEMBL120862 | 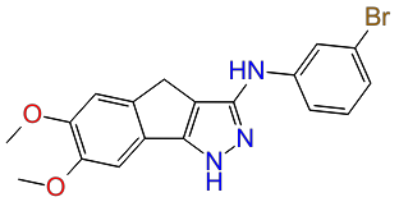 | 4.00 | train |
| 10. | CHEMBL184948 | 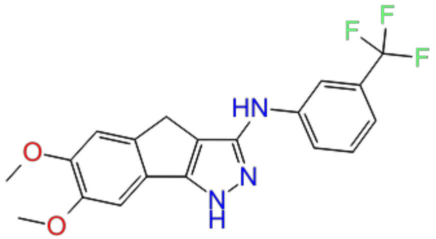 | 4.00 | test |
| 11. | CHEMBL371910 | 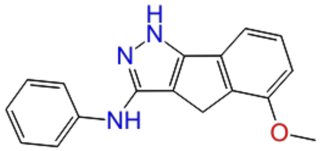 | 4.00 | train |
| 12. | CHEMBL371949 | 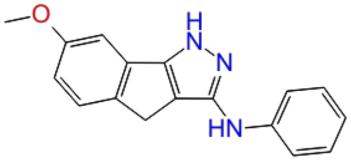 | 4.00 | train |

| | | | | |
|---|---|---|---|---|
| 13. | CHEMBL212037 | 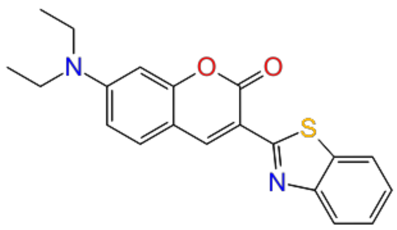 | 4.16 | test |
| 14. | CHEMBL212070 | 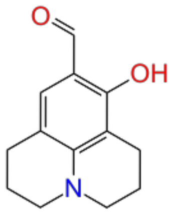 | 4.16 | train |
| 15. | CHEMBL213438 | 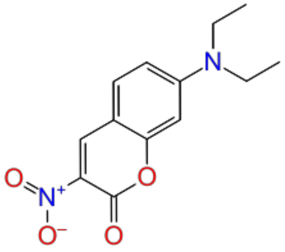 | 4.16 | train |
| 16. | CHEMBL379350 | 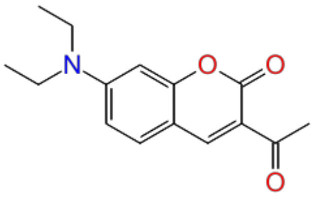 | 4.16 | test |

| | | | | |
|---|---|---|---|---|
| 17. | CHEMBL4287207 | 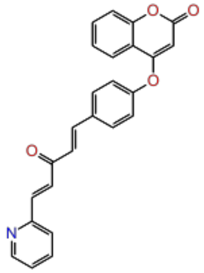 | 4.29 | train |
| 18. | CHEMBL4100030 | 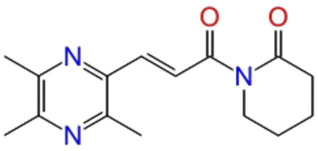 | 4.30 | train |
| 19. | CHEMBL4280398 | 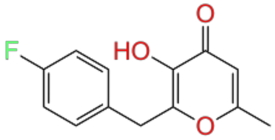 | 4.30 | train |
| 20. | CHEMBL4282653 | 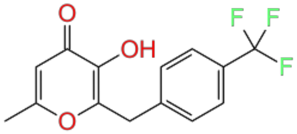 | 4.30 | train |

| | | | | |
|---|---|---|---|---|
| 21. | CHEMBL4283735 | 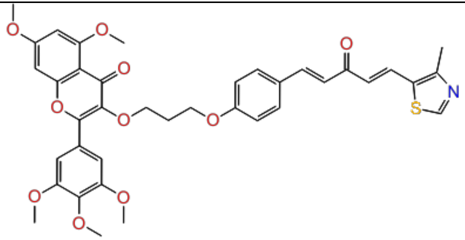 | 4.39 | test |
| 22. | CHEMBL589107 | 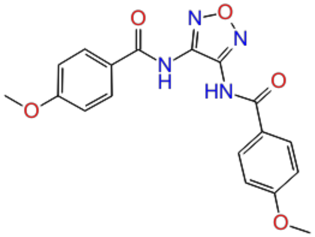 | 4.40 | train |
| 23. | CHEMBL212242 | 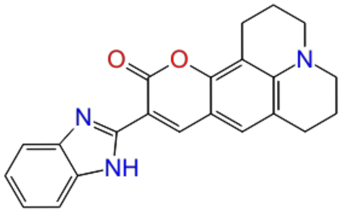 | 4.48 | train |
| 24. | CHEMBL4080967 | 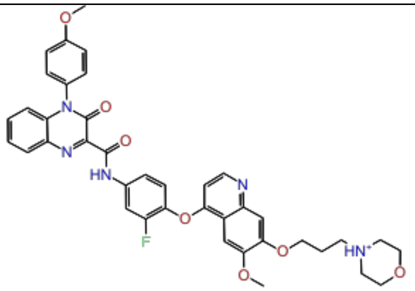 | 4.50 | train |

| | | | | |
|---|---|---|---|---|
| 25. | CHEMBL3588940 | 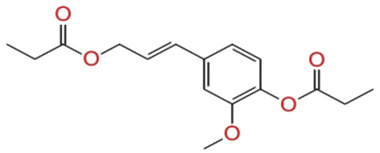 | 4.54 | train |
| 26. | CHEMBL4287242 | 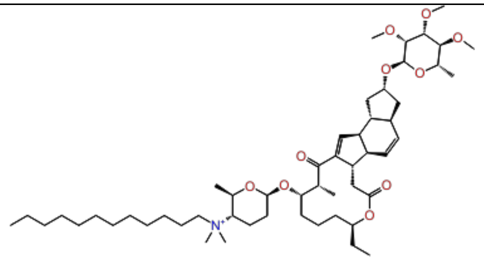 | 4.55 | test |
| 27. | CHEMBL4286444 | 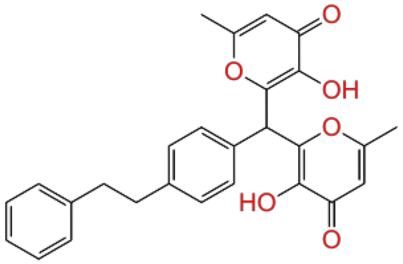 | 4.57 | test |
| 28. | CHEMBL591985 | 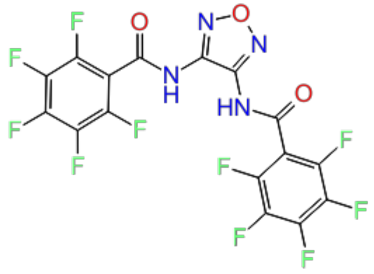 | 4.59 | train |

| | | | | |
|---|---|---|---|---|
| 29. | CHEMBL4060108 | 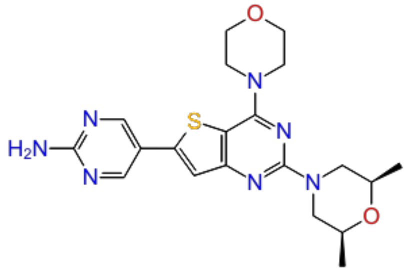 | 4.60 | train |
| 30. | CHEMBL4065513 | 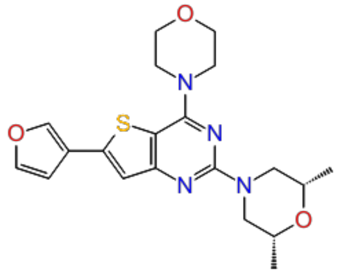 | 4.60 | train |
| 31. | CHEMBL4068220 | 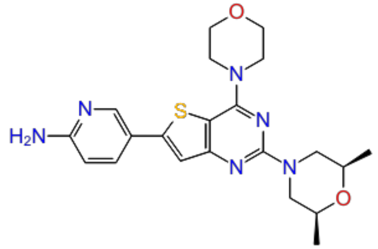 | 4.60 | test |
| 32. | CHEMBL4092492 | 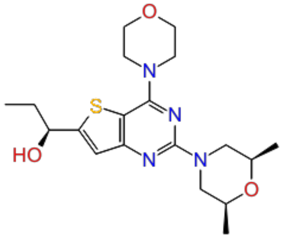 | 4.60 | train |

| | | | | |
|---|---|---|---|---|
| 33. | CHEMBL4100237 | 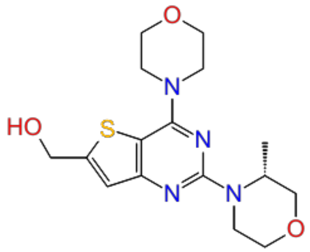 | 4.60 | train |
| 34. | CHEMBL432219 | 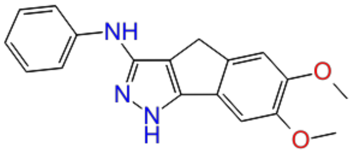 | 4.68 | train |
| 35. | CHEMBL4284792 | 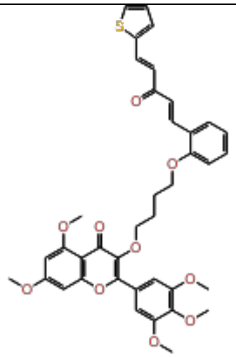 | 4.68 | train |
| 36. | CHEMBL4276849 | 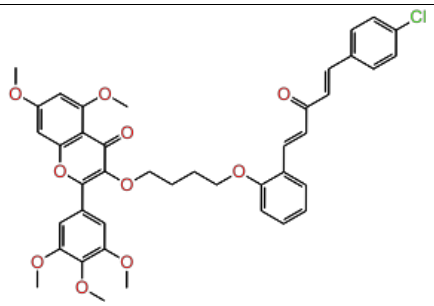 | 4.68 | test |

| | | | | |
|---|---|---|---|---|
| 37. | CHEMBL590061 | 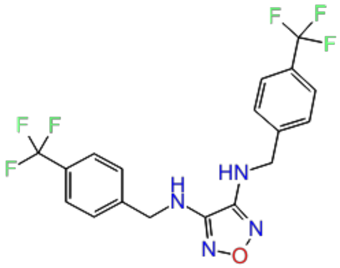 | 4.69 | train |
| 38. | CHEMBL4283971 | 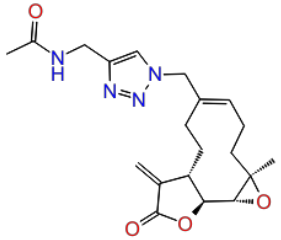 | 4.70 | test |
| 39. | CHEMBL3588941 | 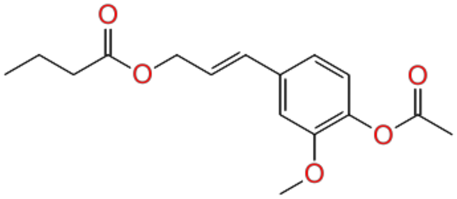 | 4.73 | train |
| 40. | CHEMBL590060 | 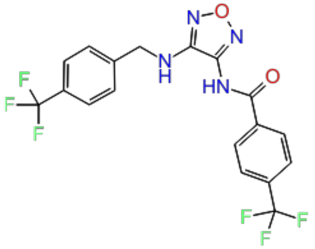 | 4.74 | train |

| | | | | |
|---|---|---|---|---|
| 41. | CHEMBL591752 | 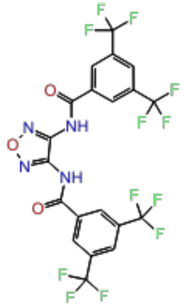 | 4.75 | test |
| 42. | CHEMBL600041 | 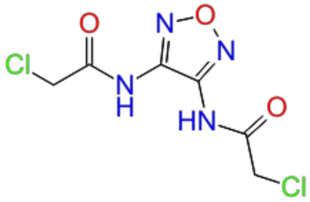 | 4.77 | train |
| 43. | CHEMBL465843 | 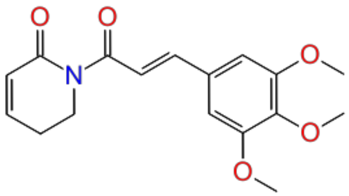 | 4.79 | train |
| 44. | CHEMBL303697 | 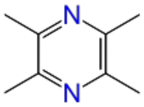 | 4.79 | train |

| | | | | |
|---|---|---|---|---|
| 45. | CHEMBL4227076 | 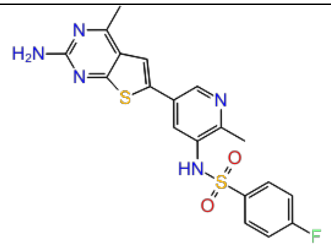 | 4.82 | test |
| 46. | CHEMBL3588943 | 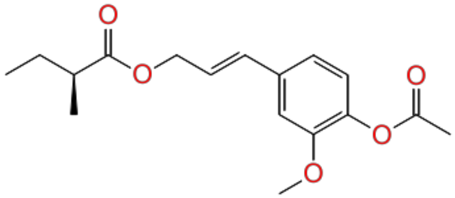 | 4.82 | train |
| 47. | CHEMBL591033 | 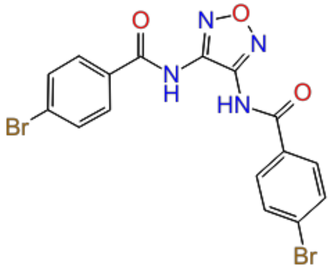 | 4.85 | test |
| 48. | CHEMBL4294821 | 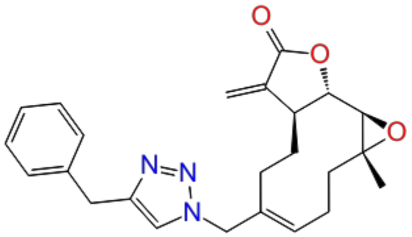 | 4.88 | train |

| | | | | |
|---|---|---|---|---|
| 49. | CHEMBL4086949 | 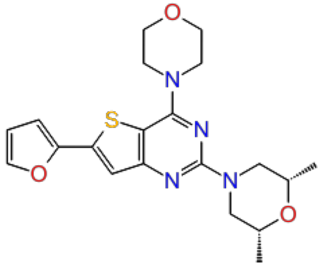 | 4.89 | train |
| 50. | CHEMBL4293259 | 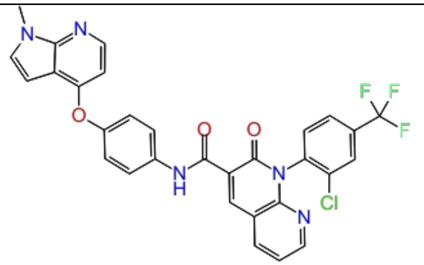 | 4.90 | train |
| 51. | CHEMBL4280764 | 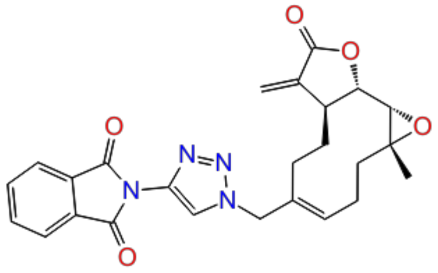 | 4.91 | train |
| 52. | CHEMBL4293238 | 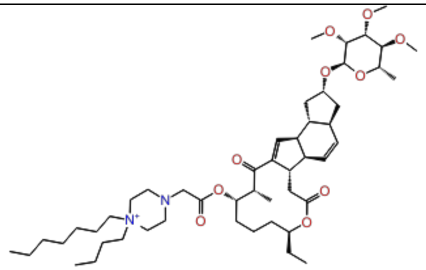 | 4.92 | test |

| | | | | |
|---|---|---|---|---|
| 53. | CHEMBL4214300 | 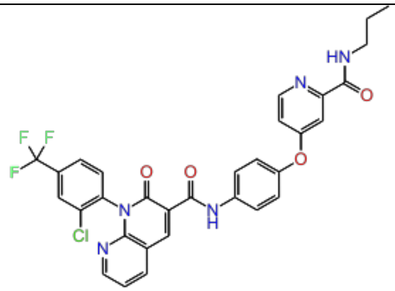 | 4.96 | train |
| 54. | CHEMBL4100335 | 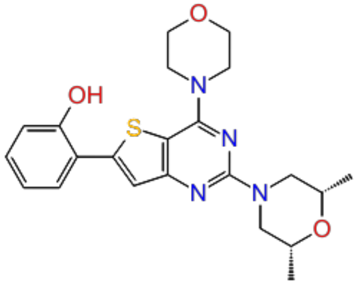 | 5.00 | train |
| 55. | CHEMBL4279750 | 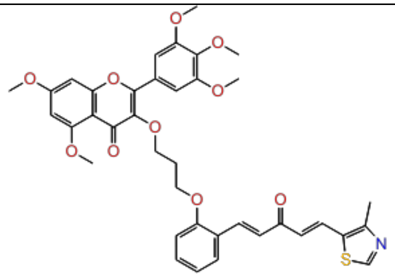 | 5.02 | train |
| 56. | CHEMBL4228112 | 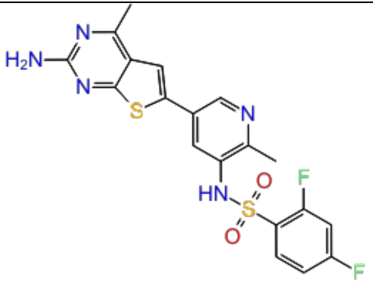 | 5.06 | test |

| | | | | |
|---|---|---|---|---|
| 57. | CHEMBL3612318 | 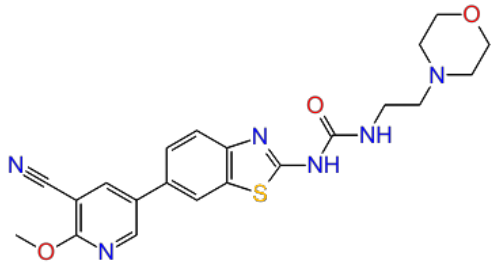 | 5.08 | train |
| 58. | CHEMBL590806 | 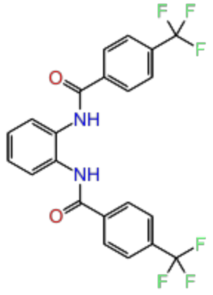 | 5.10 | test |
| 59. | CHEMBL118201 | 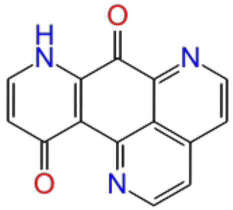 | 5.10 | train |
| 60. | CHEMBL1644111 | 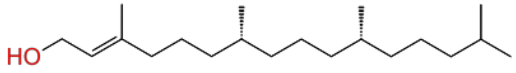 | 5.10 | train |

| | | | | |
|---|---|---|---|---|
| 61. | CHEMBL4215210 | 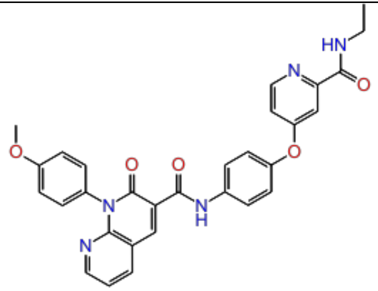 | 5.14 | train |
| 62. | CHEMBL4078390 | 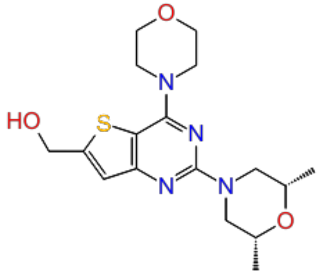 | 5.18 | test |
| 63. | CHEMBL563939 | 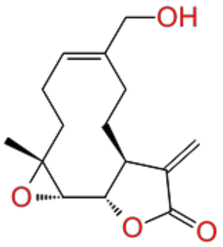 | 5.20 | train |
| 64. | CHEMBL4279674 | 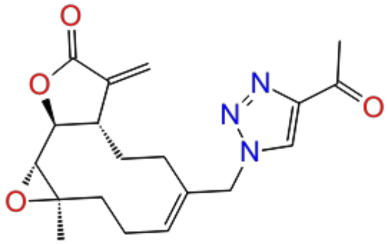 | 5.21 | train |

| | | | | |
|---|---|---|---|---|
| 65. | CHEMBL118845 | 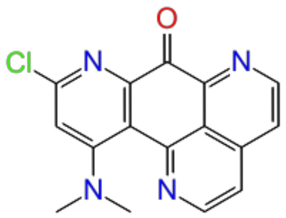 | 5.22 | train |
| 66. | CHEMBL4291410 | 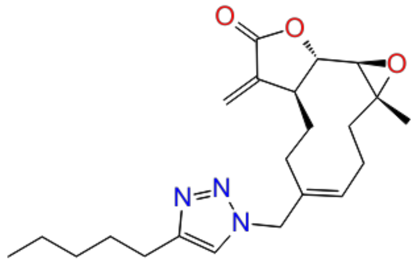 | 5.24 | test |
| 67. | CHEMBL4278116 | 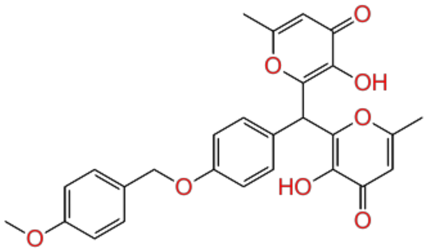 | 5.28 | test |
| 68. | CHEMBL331562 | 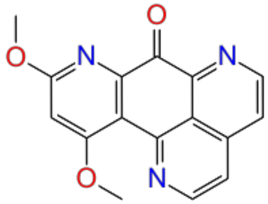 | 5.30 | train |

| | | | | |
|---|---|---|---|---|
| 69. | CHEMBL4280439 | 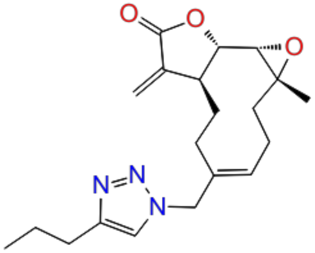 | 5.30 | train |
| 70. | CHEMBL4286032 | 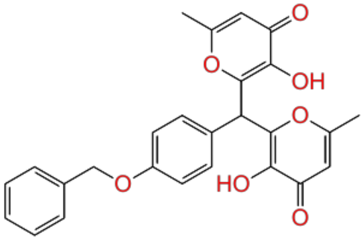 | 5.34 | train |
| 71. | CHEMBL4098035 | 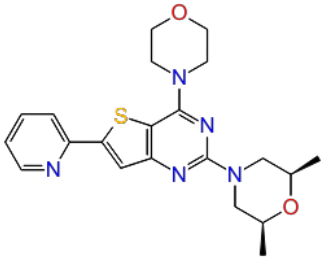 | 5.35 | test |
| 72. | CHEMBL4294566 | 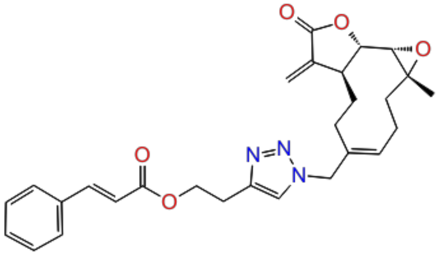 | 5.36 | train |

| | | | | |
|---|---|---|---|---|
| 73. | CHEMBL122557 | 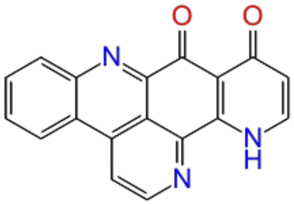 | 5.40 | train |
| 74. | CHEMBL2152488 | 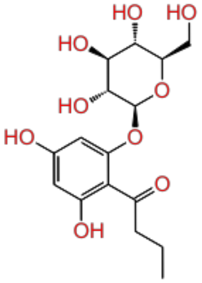 | 5.41 | train |
| 75. | CHEMBL385786 | 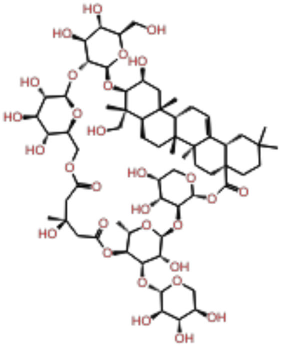 | 5.44 | train |

| | | | | |
|---|---|---|---|---|
| 76. | CHEMBL4281496 | 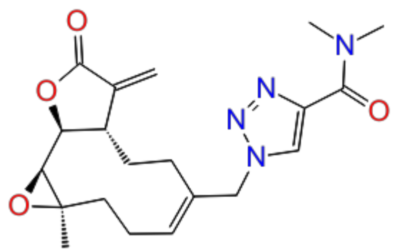 | 5.45 | test |
| 77. | CHEMBL4204238 | 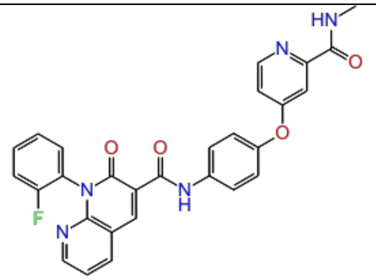 | 5.47 | train |
| 78. | CHEMBL4064087 | 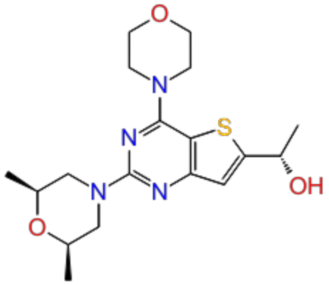 | 5.48 | train |
| 79. | CHEMBL4209984 | 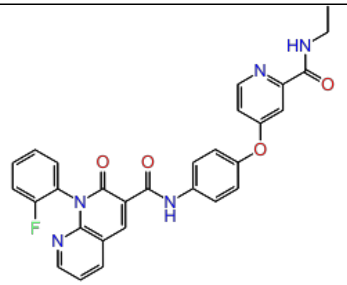 | 5.49 | train |

| | | | | |
|---|---|---|---|---|
| 80. | CHEMBL4225944 | 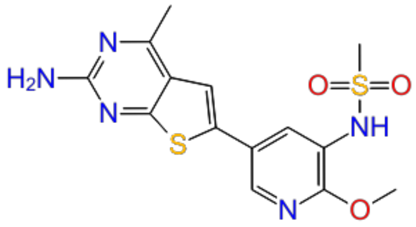 | 5.52 | test |
| 81. | CHEMBL4282461 | 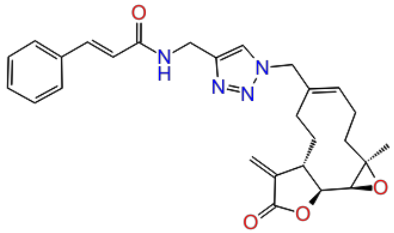 | 5.53 | train |
| 82. | CHEMBL4280635 | 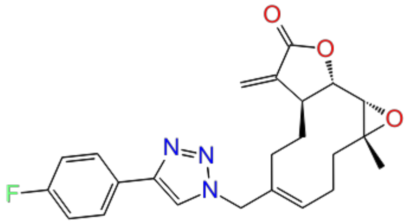 | 5.55 | train |
| 83. | CHEMBL3659991 | 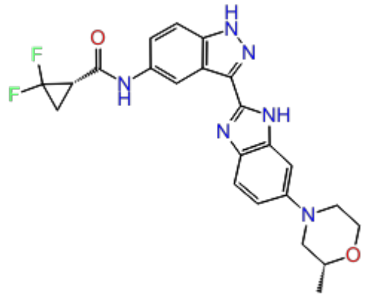 | 5.57 | test |

| | | | | |
|---|---|---|---|---|
| 84. | CHEMBL4066144 | 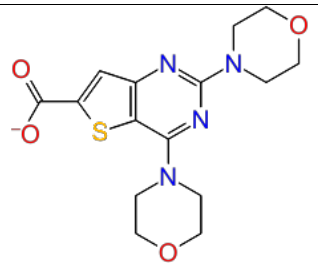 | 5.61 | train |
| 85. | CHEMBL4226082 | 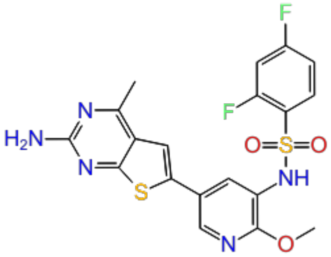 | 5.62 | train |
| 86. | CHEMBL4285130 | 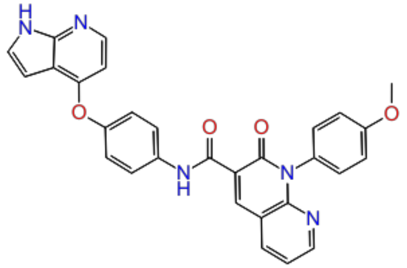 | 5.63 | test |
| 87. | CHEMBL4293054 | 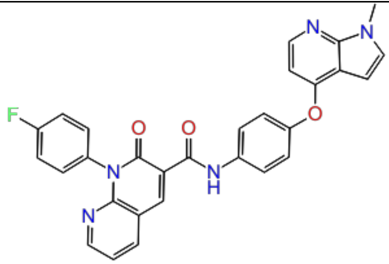 | 5.67 | test |

| | | | | |
|---|---|---|---|---|
| 88. | CHEMBL4225722 | 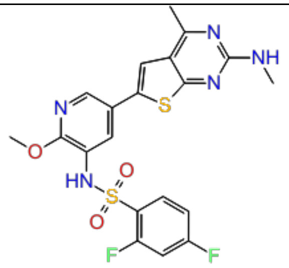 | 5.68 | train |
| 89. | CHEMBL111249 | 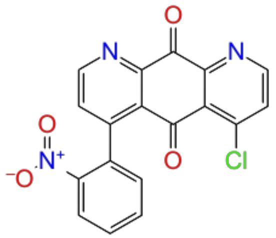 | 5.70 | train |
| 90. | CHEMBL4288866 | 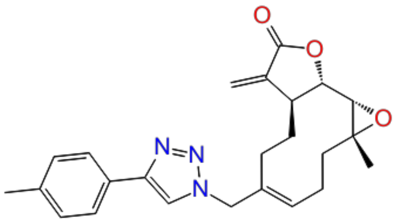 | 5.72 | train |
| 91. | CHEMBL4072336 | 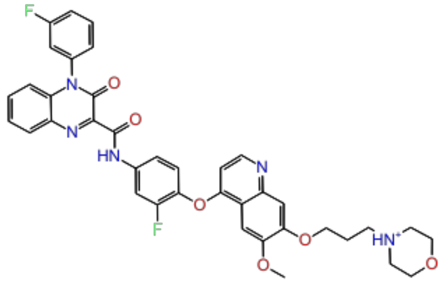 | 5.73 | test |

| | | | | |
|---|---|---|---|---|
| 92. | CHEMBL373587 | 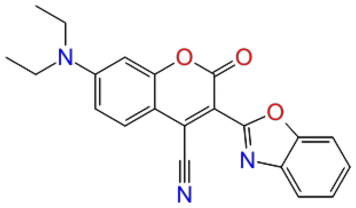 | 5.75 | train |
| 93. | CHEMBL4074866 | 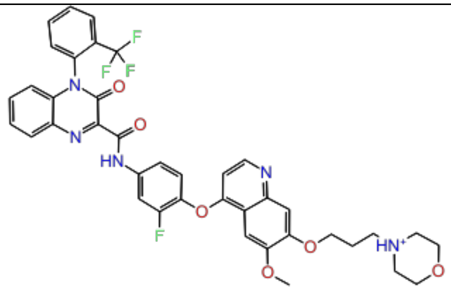 | 5.75 | train |
| 94. | CHEMBL3612319 | 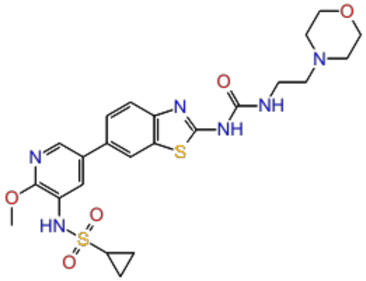 | 5.78 | train |
| 95. | CHEMBL4084146 | 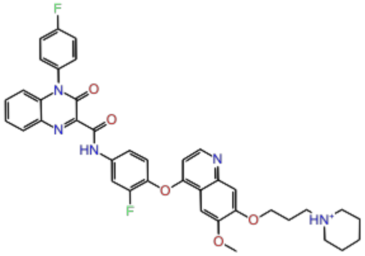 | 5.79 | train |

| | | | | |
|---|---|---|---|---|
| 96. | CHEMBL4097578 | 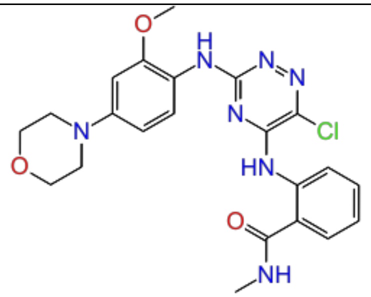 | 5.82 | test |
| 97. | CHEMBL4207773 | 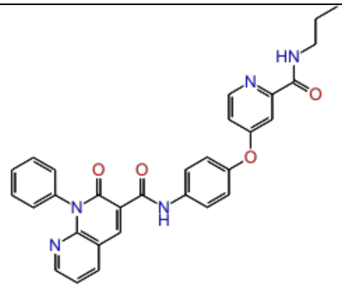 | 5.82 | train |
| 98. | CHEMBL3612194 | 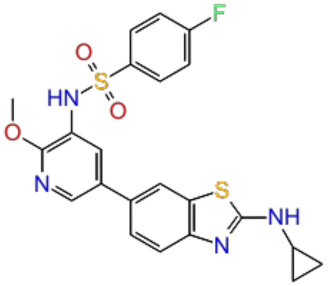 | 5.82 | train |

| | | | | |
|---|---|---|---|---|
| 99. | CHEMBL4288118 | 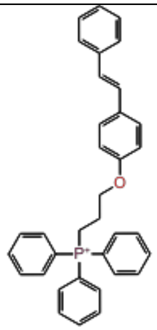 | 5.85 | train |
| 100. | CHEMBL4075562 | 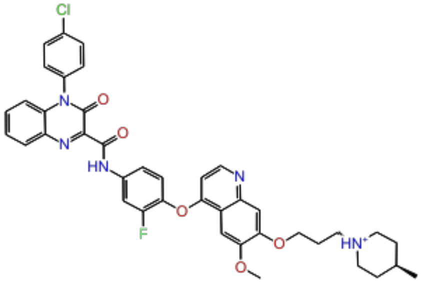 | 5.85 | test |
| 101. | CHEMBL3612197 | 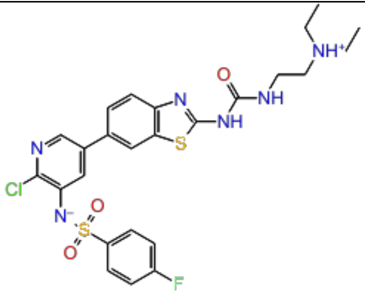 | 5.86 | test |
| 102. | CHEMBL4289873 | 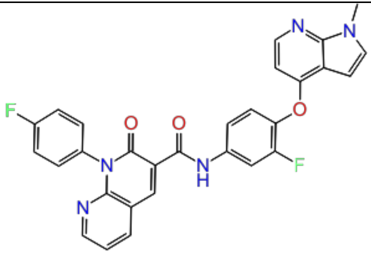 | 5.88 | train |

| | | | | |
|---|---|---|---|---|
| 103. | CHEMBL4277119 | 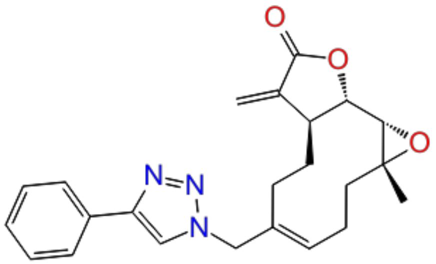 | 5.90 | train |
| 104. | CHEMBL4278233 | 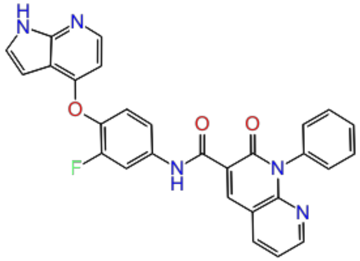 | 5.90 | test |
| 105. | CHEMBL4285131 | 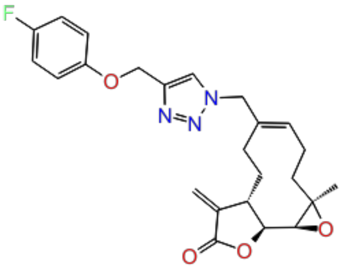 | 5.90 | train |
| 106. | CHEMBL4294165 | 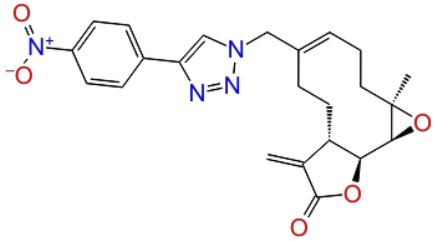 | 5.91 | train |

| | | | | |
|---|---|---|---|---|
| 107. | CHEMBL4059832 | 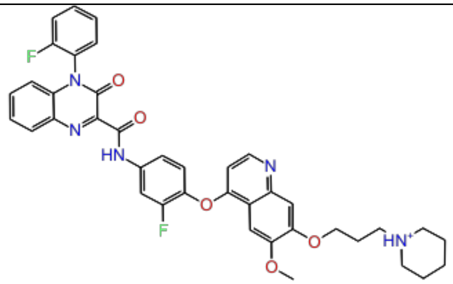 | 5.93 | train |
| 108. | CHEMBL4214671 | 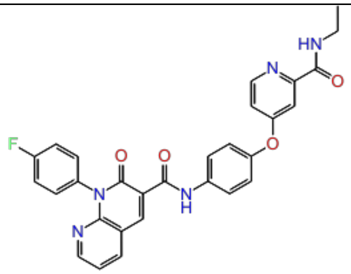 | 5.95 | train |
| 109. | CHEMBL3612315 | 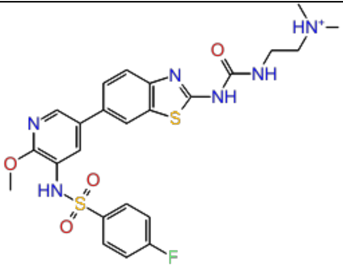 | 5.99 | test |
| 110. | CHEMBL4244147 | 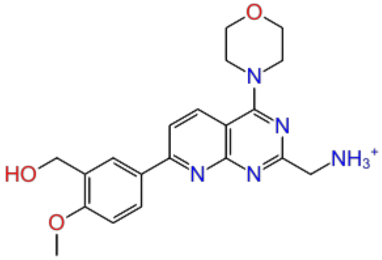 | 6.00 | train |

| | | | | |
|---|---|---|---|---|
| 111. | CHEMBL4242668 | 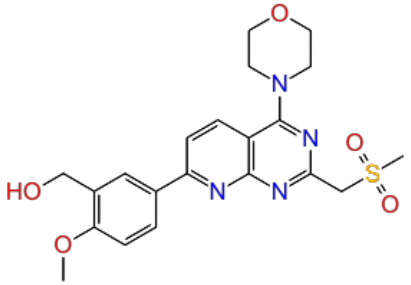 | 6.00 | train |
| 112. | CHEMBL4225819 | 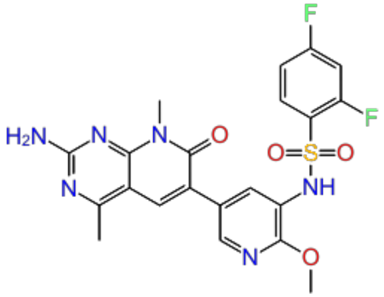 | 6.02 | test |
| 113. | CHEMBL4204920 | 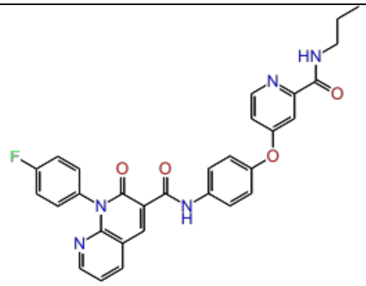 | 6.02 | train |
| 114. | CHEMBL1230609 | 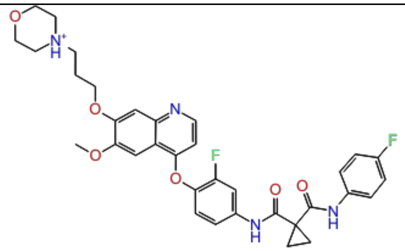 | 6.02 | train |

| | | | | |
|---|---|---|---|---|
| 115. | CHEMBL4286698 | 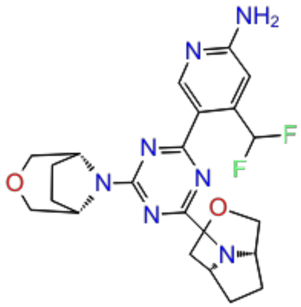 | 6.04 | train |
| 116. | CHEMBL4239569 | 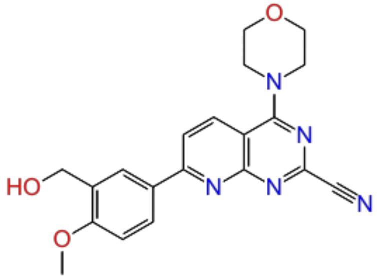 | 6.04 | test |
| 117. | CHEMBL4098250 | 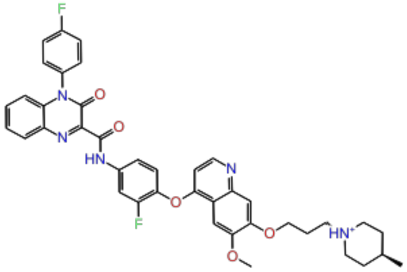 | 6.06 | train |
| 118. | CHEMBL117616 | 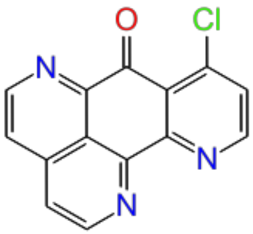 | 6.10 | train |

| | | | | |
|---|---|---|---|---|
| 119. | CHEMBL118709 | 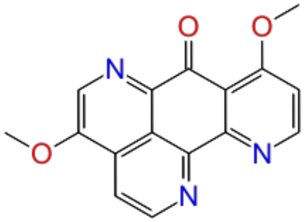 | 6.10 | train |
| 120. | CHEMBL111300 | 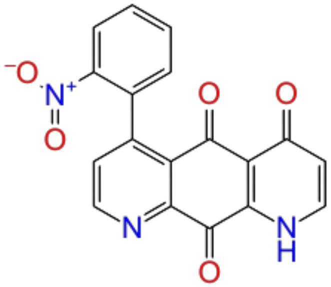 | 6.10 | test |
| 121. | CHEMBL111383 | 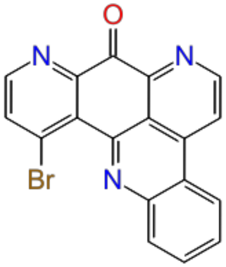 | 6.16 | train |

| | | | | |
|---|---|---|---|---|
| 122. | CHEMBL4065141 | 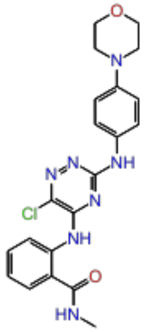 | 6.16 | train |
| 123. | CHEMBL4225483 | 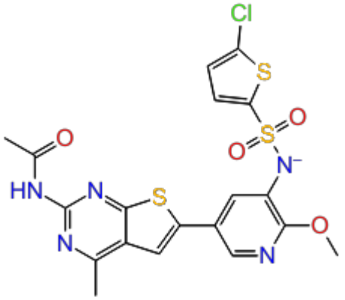 | 6.16 | test |
| 124. | CHEMBL4092858 | 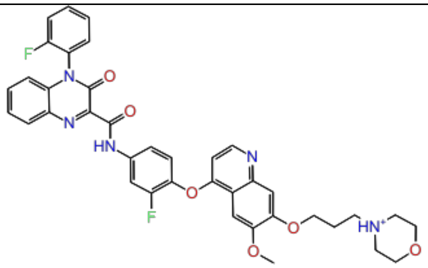 | 6.19 | train |
| 125. | CHEMBL4280452 | 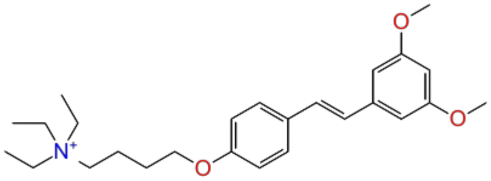 | 6.24 | test |

| | | | | |
|---|---|---|---|---|
| 126. | CHEMBL521851 | 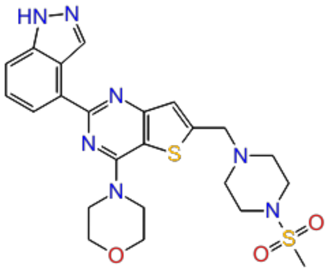 | 6.26 | train |
| 127. | CHEMBL3612321 | 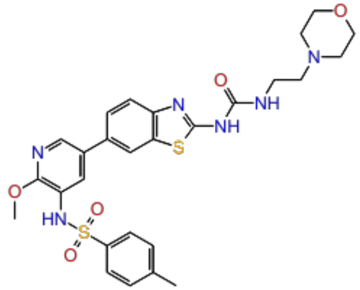 | 6.28 | train |
| 128. | CHEMBL325202 | 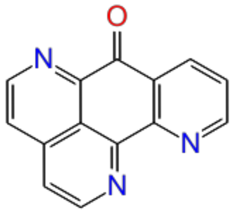 | 6.30 | test |
| 129. | CHEMBL4092812 | 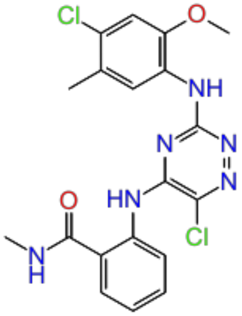 | 6.30 | train |

| | | | | |
|---|---|---|---|---|
| 130. | CHEMBL3612322 | 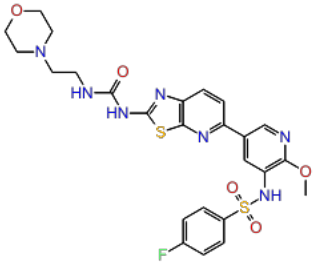 | 6.32 | train |
| 131. | CHEMBL4284998 | 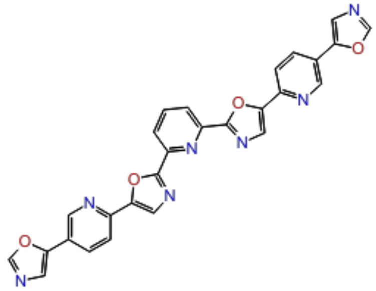 | 6.36 | test |
| 132. | CHEMBL3612314 | 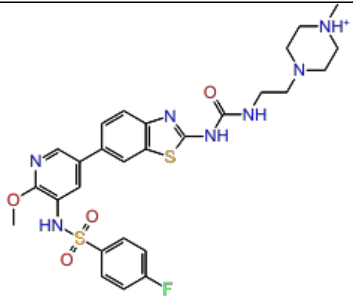 | 6.36 | train |
| 133. | CHEMBL4071083 | 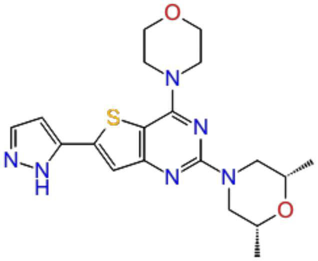 | 6.37 | train |

| | | | | |
|---|---|---|---|---|
| 134. | CHEMBL4291927 | 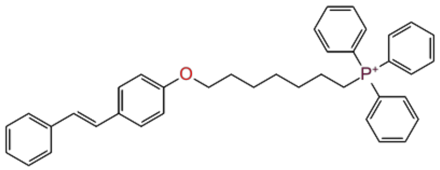 | 6.44 | train |
| 135. | CHEMBL3612312 | 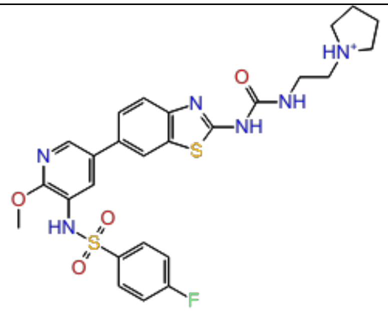 | 6.44 | train |
| 136. | CHEMBL1234354 | 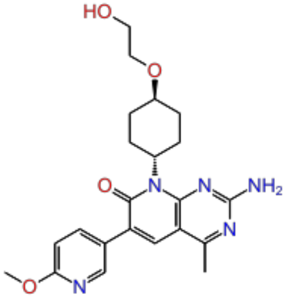 | 6.47 | test |
| 137. | CHEMBL112938 | 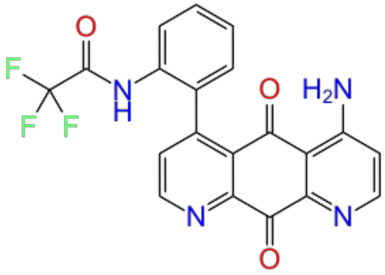 | 6.52 | train |

| | | | | |
|---|---|---|---|---|
| 138. | CHEMBL4284890 | 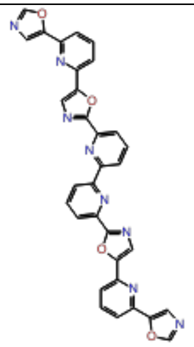 | 6.61 | train |
| 139. | CHEMBL4159104 | 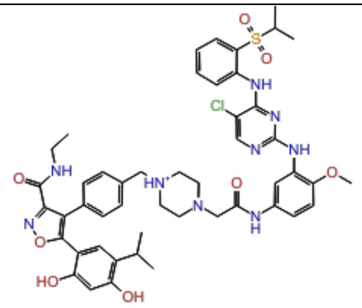 | 6.62 | train |
| 140. | CHEMBL359744 | 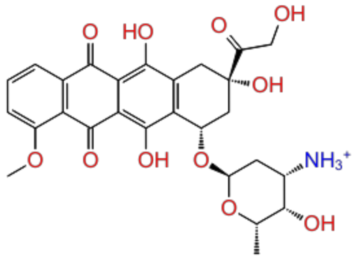 | 6.80 | test |
| 141. | CHEMBL4072042 | 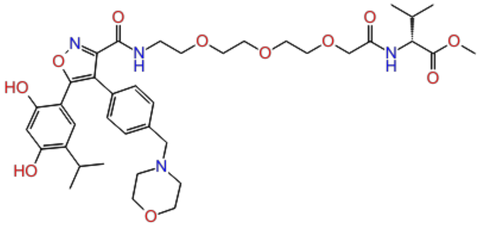 | 6.96 | train |

| | | | | |
|---|---|---|---|---|
| 142. | CHEMBL327101 | 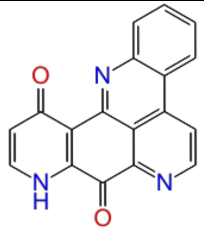 | 7.00 | train |
| 143. | CHEMBL117784 | 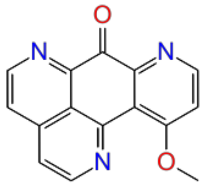 | 7.00 | train |
| 144. | CHEMBL4280053 | 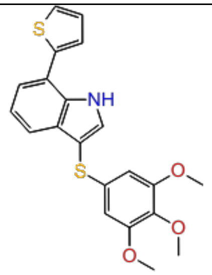 | 7.02 | test |
| 145. | CHEMBL4290427 | 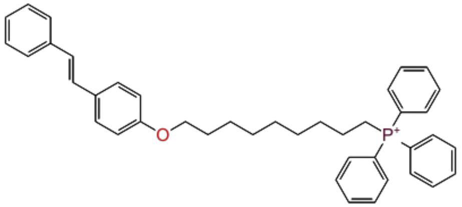 | 7.05 | test |

| | | | | |
|---|---|---|---|---|
| 146. | CHEMBL4246481 | 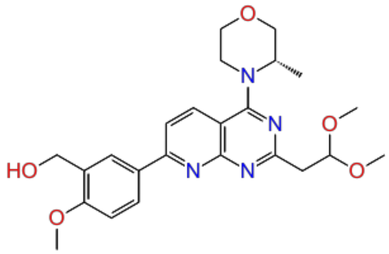 | 7.05 | train |
| 147. | CHEMBL4246121 | 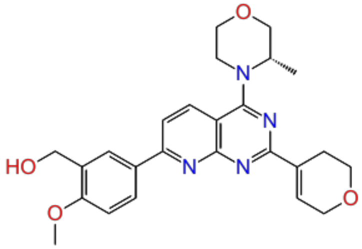 | 7.07 | train |
| 148. | CHEMBL122076 | 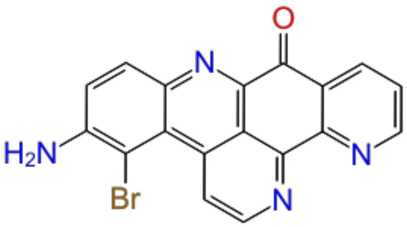 | 7.10 | train |
| 149. | CHEMBL125395 | 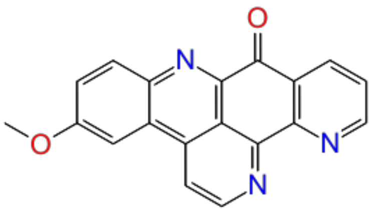 | 7.16 | train |

| | | | | |
|---|---|---|---|---|
| 150. | CHEMBL121911 | 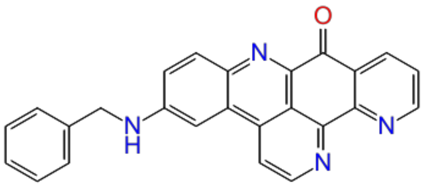 | 7.16 | train |
| 151. | CHEMBL340160 | 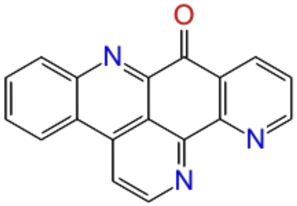 | 7.16 | test |
| 152. | CHEMBL4060227 | 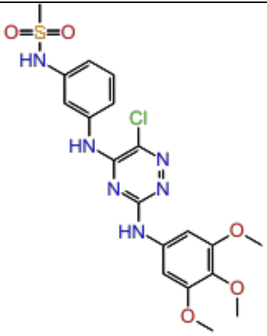 | 7.16 | train |
| 153. | CHEMBL4281312 | 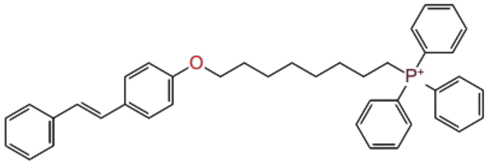 | 7.16 | test |

| | | | | |
|---|---|---|---|---|
| 154. | CHEMBL323280 | 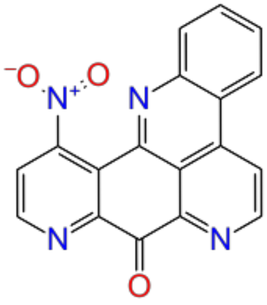 | 7.16 | train |
| 155. | CHEMBL324780 | 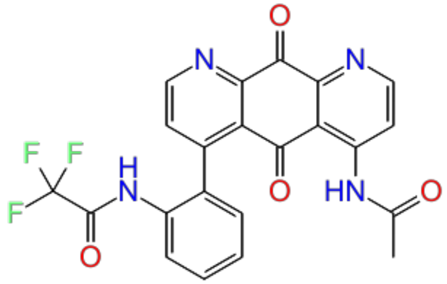 | 7.22 | test |
| 156. | CHEMBL324884 | 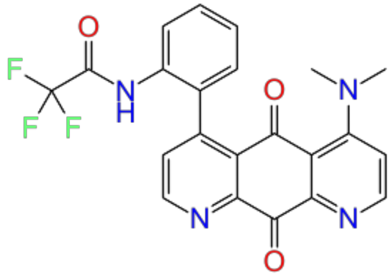 | 7.22 | train |
| 157. | CHEMBL4280206 | 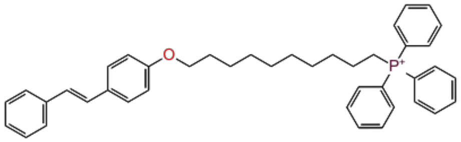 | 7.22 | train |

| | | | | |
|---|---|---|---|---|
| 158. | CHEMBL333669 | 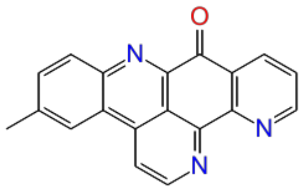 | 7.22 | test |
| 159. | CHEMBL109479 | 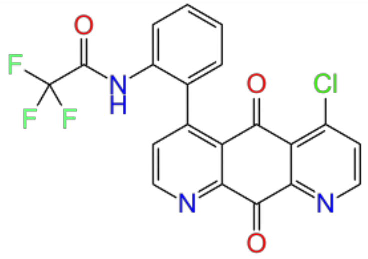 | 7.30 | train |
| 160. | CHEMBL3301610 | 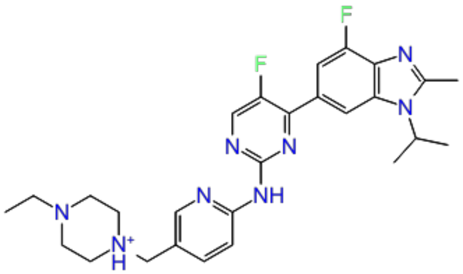 | 7.32 | train |
| 161. | CHEMBL4239712 | 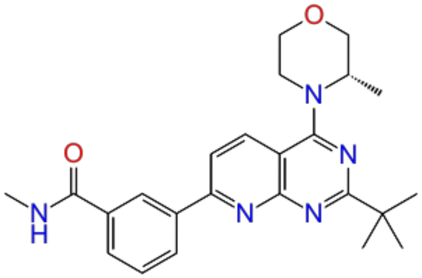 | 7.49 | train |

| | | | | |
|---|---|---|---|---|
| 162. | CHEMBL1801204 | 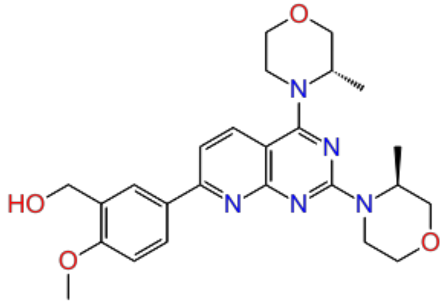 | 7.67 | train |
| 163. | CHEMBL4278554 | 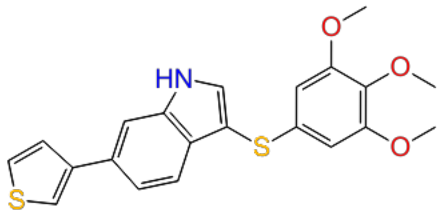 | 7.80 | train |
| 164. | CHEMBL4098626 | 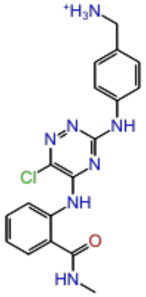 | 7.85 | test |
| 165. | CHEMBL498271 | 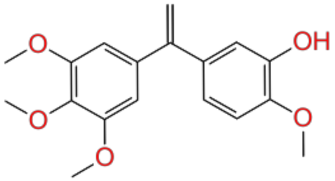 | 8.10 | test |

| | | | | |
|---|---|---|---|---|
| 166. | CHEMBL4162270 | 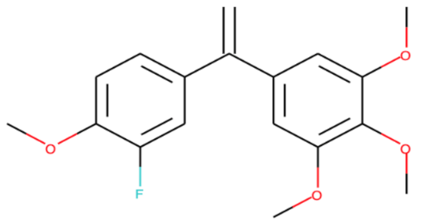 | 8.10 | train |
| 167. | CHEMBL4238656 | 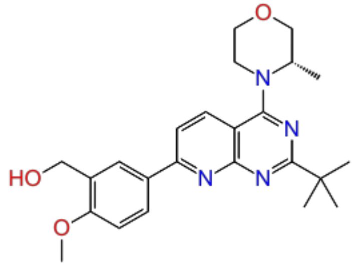 | 8.49 | train |
| 168. | CHEMBL4172942 | 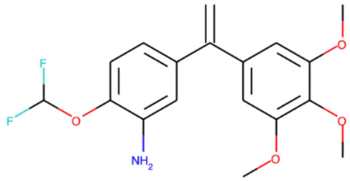 | 9.22 | train |
| 169. | CHEMBL508617 | 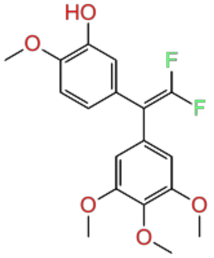 | 9.52 | train |

**Table S11:** Dataset from U251 cell line for pharmacophore hypothesis and 3D QSAR model generation.

| Sr. No | Compound ID | 2D structure | Pharm Set | #Conformations | QSAR set | Activity (Observed) | Activity (Predicted) | Fitness score |
|---|---|---|---|---|---|---|---|---|
| 1. | CHEMBL20883 | 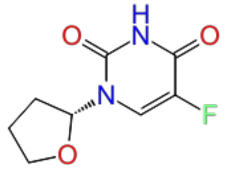 | inactive | 5 | training | 3.42 | 4.94 | 1.51 |
| 2. | CHEMBL253353 | 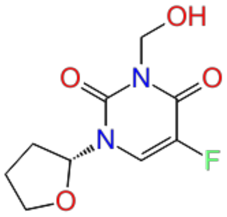 | inactive | 10 | test | 3.99 | 4.95 | 1.48 |
| 3. | CHEMBL500473 | 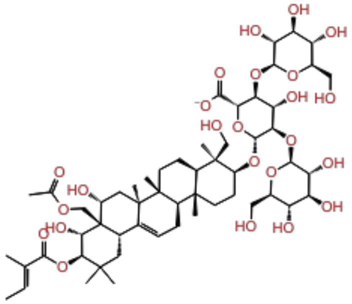 | inactive | 153 | training | 4.09 | 5.37 | 1.32 |

| | | | | | | | | |
|---|---|---|---|---|---|---|---|---|
| 4. | CHEMBL50 | 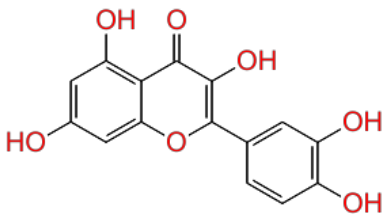 | inactive | 20 | training | 4.11 | 4.72 | 1.52 |
| 5. | CHEMBL180828 | 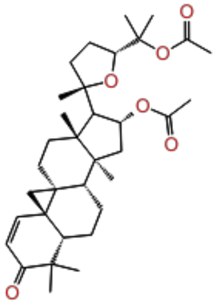 | inactive | 55 | test | 4.23 | 5.37 | 1.18 |
| 6. | CHEMBL252368 | 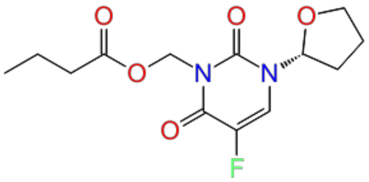 | inactive | 130 | training | 4.26 | 4.39 | 1.54 |
| 7. | CHEMBL3588940 | 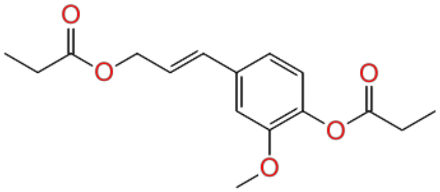 | inactive | 188 | training | 4.28 | 4.33 | 1.42 |

| | | | | | | | | |
|---|---|---|---|---|---|---|---|---|
| 8. | CHEMBL75267 | 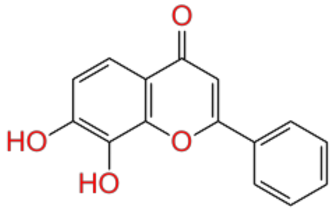 | inactive | 4 | test | 4.30 | 5.4 | 1.7 |
| 9. | CHEMBL432108 | 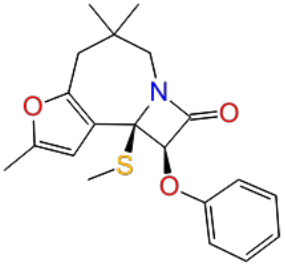 | inactive | 21 | training | 4.33 | 5.82 | 0.64 |
| 10. | CHEMBL1958318 | 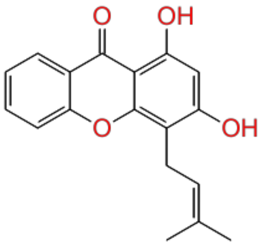 | inactive | 10 | training | 4.33 | 4.63 | 1.57 |
| 11. | CHEMBL3588942 | 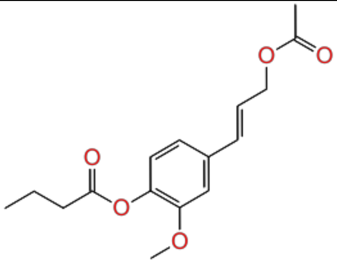 | inactive | 248 | test | 4.34 | 5.36 | 1.64 |

| | | | | | | | | |
|---|---|---|---|---|---|---|---|---|
| 12. | CHEMBL412883 | 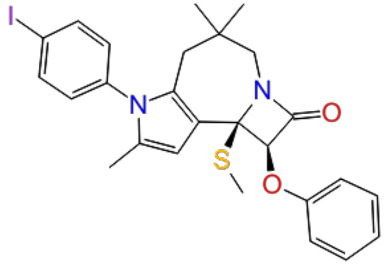 | inactive | 26 | training | 4.40 | 5.38 | 1.5 |
| 13. | CHEMBL233930 | 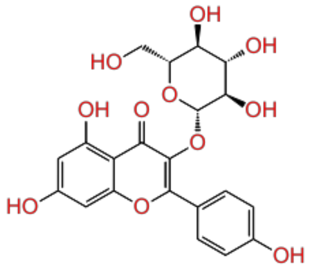 | inactive | 61 | training | 4.42 | 4.72 | 1.55 |
| 14. | CHEMBL4093795 | 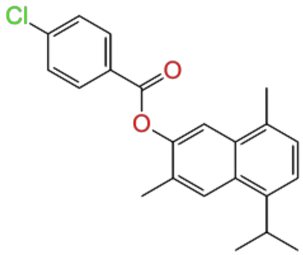 | inactive | 4 | test | 4.45 | 5.09 | 1.22 |
| 15. | CHEMBL3588941 | 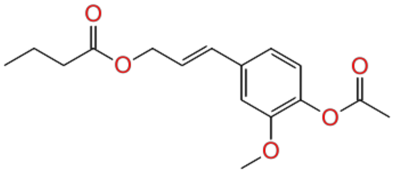 | inactive | 232 | training | 4.48 | 4.85 | 1.38 |

| | | | | | | | | |
|---|---|---|---|---|---|---|---|---|
| 16. | CHEMBL1784141 | 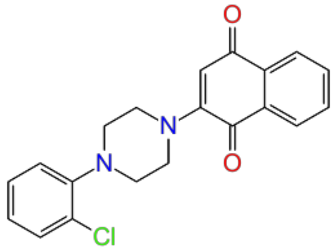 | Moderate | 50 | training | 4.51 | 5.04 | 1.47 |
| 17. | CHEMBL359600 | 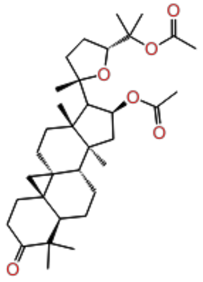 | Moderate | 68 | test | 4.52 | 4.94 | 1.22 |
| 18. | CHEMBL3588943 | 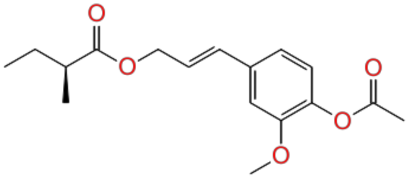 | Moderate | 250 | training | 4.55 | 4.56 | 1.38 |
| 19. | CHEMBL187044 | 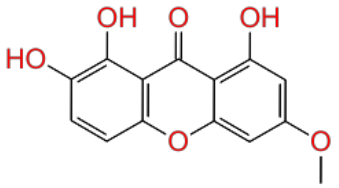 | Moderate | 5 | test | 4.56 | 4.62 | 1.39 |

| | | | | | | | | |
|---|---|---|---|---|---|---|---|---|
| 20. | CHEMBL1644111 | 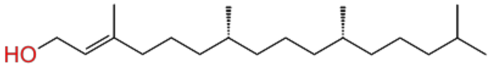 | Moderate | 512 | training | 4.56 | 4.07 | 1.28 |
| 21. | CHEMBL182093 | 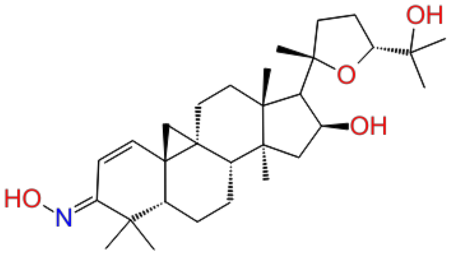 | Moderate | 65 | training | 4.57 | 6.6 | 1.24 |
| 22. | CHEMBL3588938 | 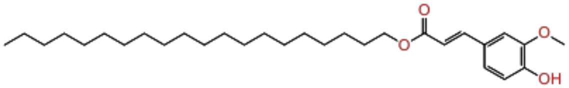 | Moderate | 203 | training | 4.57 | 4.4 | 1.1 |
| 23. | CHEMBL298047 | 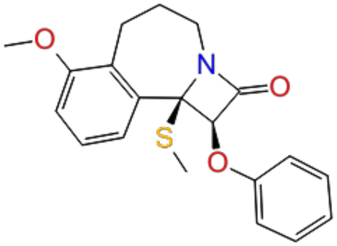 | Moderate | 17 | test | 4.60 | 6.28 | 1.49 |

| | | | | | | | | |
|---|---|---|---|---|---|---|---|---|
| 24. | CHEMBL1958315 | 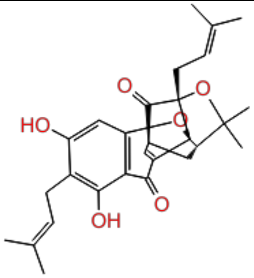 | Moderate | 132 | training | 4.65 | 3.65 | 1.23 |
| 25. | CHEMBL4103953 | 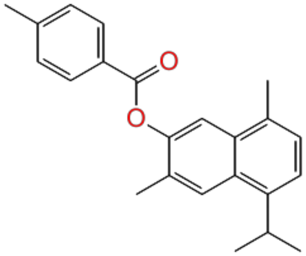 | Moderate | 4 | training | 4.65 | 5.09 | 1.22 |
| 26. | CHEMBL2152488 | 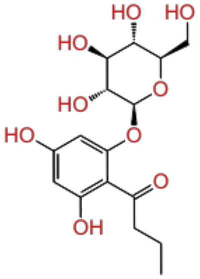 | Moderate | 141 | test | 4.71 | 5.79 | 1.49 |
| 27. | CHEMBL1958317 | 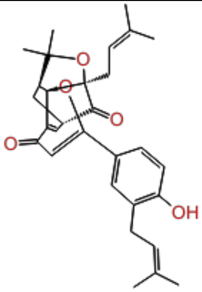 | Moderate | 194 | training | 4.73 | 3.97 | 1.26 |

| | | | | | | | | |
|---|---|---|---|---|---|---|---|---|
| 28. | CHEMBL1958319 | 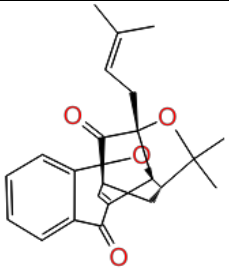 | Moderate | 11 | training | 4.73 | 3.74 | 1.21 |
| 29. | CHEMBL4087813 | 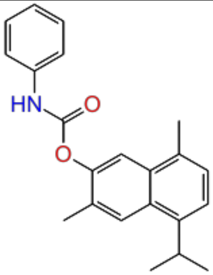 | Moderate | 16 | test | 4.80 | 4.06 | 1.17 |
| 30. | CHEMBL4172865 | 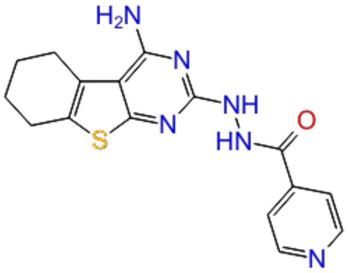 | Moderate | 18 | training | 4.88 | 5.49 | 1.55 |
| 31. | CHEMBL3331344 | 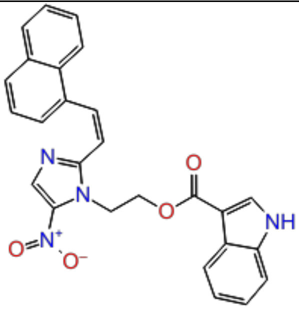 | Moderate | 183 | training | 4.89 | 5.92 | 1.24 |

| | | | | | | | | |
|---|---|---|---|---|---|---|---|---|
| 32. | CHEMBL178232 | 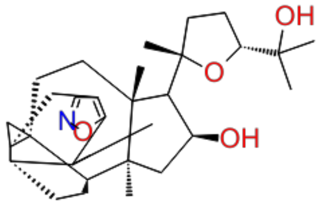 | Moderate | 51 | test | 4.95 | 4.69 | 0.44 |
| 33. | CHEMBL1784156 | 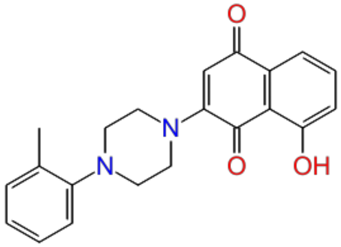 | Moderate | 51 | training | 4.99 | 4.65 | 1.59 |
| 34. | CHEMBL3311152 | 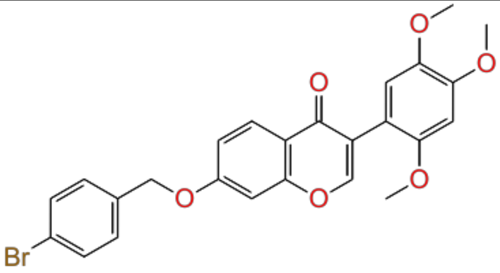 | Moderate | 81 | training | 5.00 | 5.02 | 1.09 |
| 35. | CHEMBL3588939 | 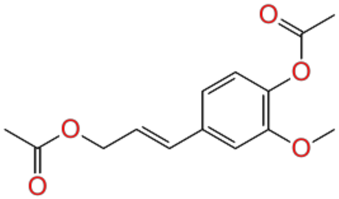 | Moderate | 83 | test | 5.01 | 3.78 | 1.48 |

| | | | | | | | | |
|---|---|---|---|---|---|---|---|---|
| 36. | CHEMBL3311154 | 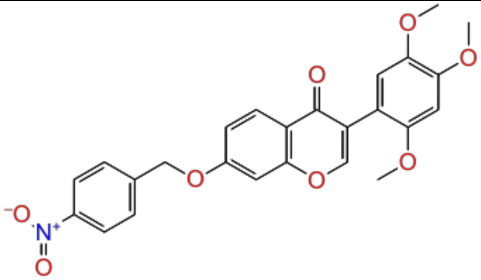 | Moderate | 81 | training | 5.03 | 4.91 | 1.19 |
| 37. | CHEMBL1784154 | 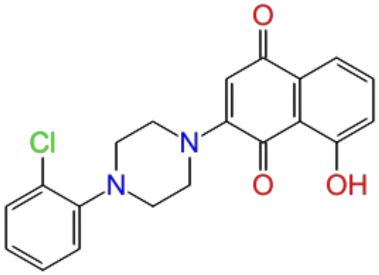 | Moderate | 51 | training | 5.10 | 4.58 | 1.57 |
| 38. | CHEMBL3311147 | 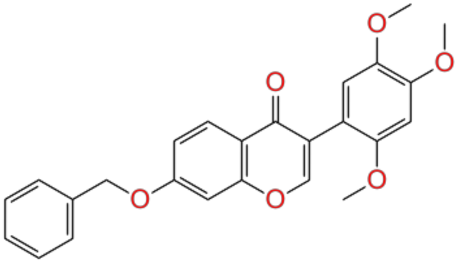 | Moderate | 83 | test | 5.17 | 4.91 | 1.2 |
| 39. | CHEMBL3311146 | 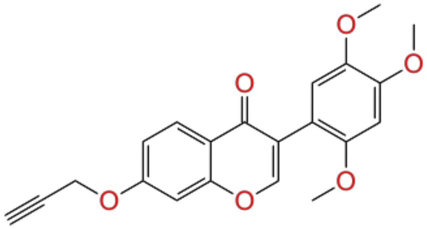 | Moderate | 75 | training | 5.19 | 5.49 | 1.24 |

| | | | | | | | | |
|---|---|---|---|---|---|---|---|---|
| 40. | CHEMBL1784150 | 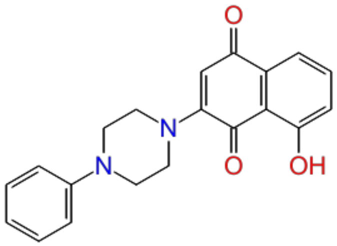 | Moderate | 29 | test | 5.21 | 5.92 | 1.19 |
| 41. | CHEMBL1784161 | 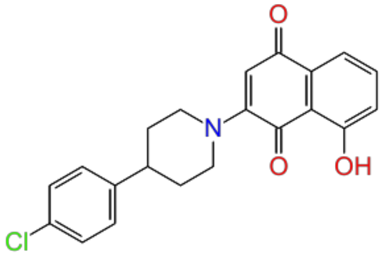 | Moderate | 28 | training | 5.21 | 6.26 | 1.48 |
| 42. | CHEMBL3311157 | 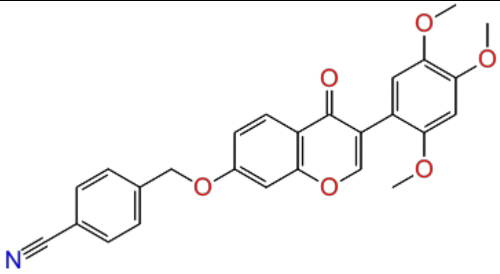 | Moderate | 78 | training | 5.21 | 4.97 | 1.22 |
| 43. | CHEMBL3311145 | 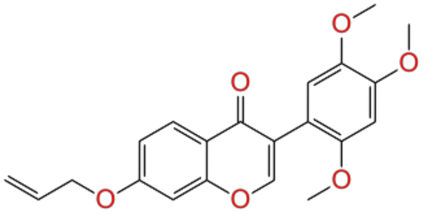 | Moderate | 110 | training | 5.23 | 5.2 | 1.28 |

| | | | | | | | | |
|---|---|---|---|---|---|---|---|---|
| 44. | CHEMBL446125 | 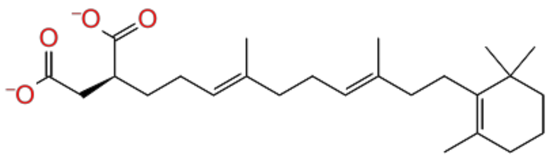 | Moderate | 515 | test | 5.39 | - | - |
| 45. | CHEMBL3311038 | 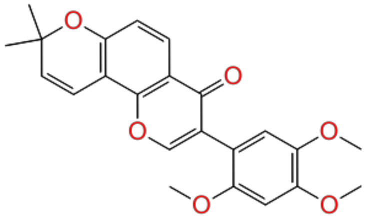 | Moderate | 27 | training | 5.39 | 4.9 | 1.66 |
| 46. | CHEMBL3330998 | 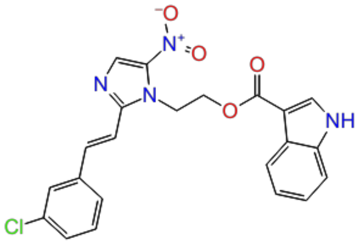 | Moderate | 90 | training | 5.39 | 5.53 | 1.27 |
| 47. | CHEMBL3331001 | 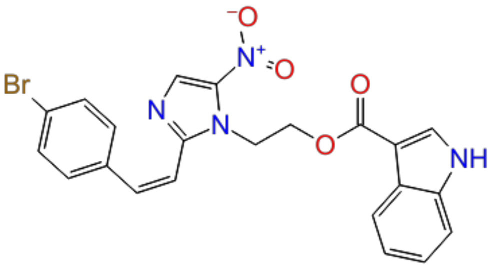 | Moderate | 115 | test | 5.41 | 4.43 | 1.45 |

| | | | | | | | | |
|---|---|---|---|---|---|---|---|---|
| 48. | CHEMBL258765 | 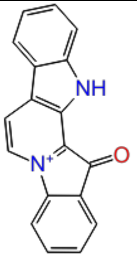 | Moderate | 1 | training | 5.44 | 5.08 | 1.43 |
| 49. | CHEMBL3311061 | 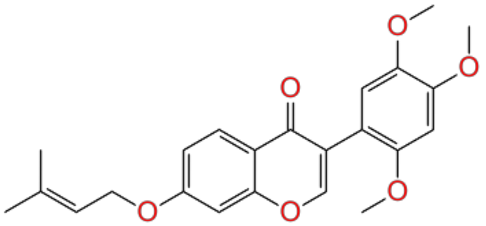 | Moderate | 162 | training | 5.46 | 5.54 | 1.22 |
| 50. | CHEMBL3329220 | 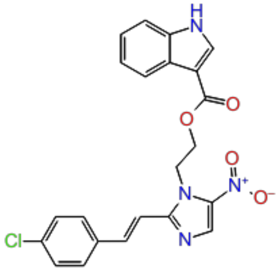 | Moderate | 50 | test | 5.49 | 5.13 | 1.53 |
| 51. | CHEMBL3311151 | 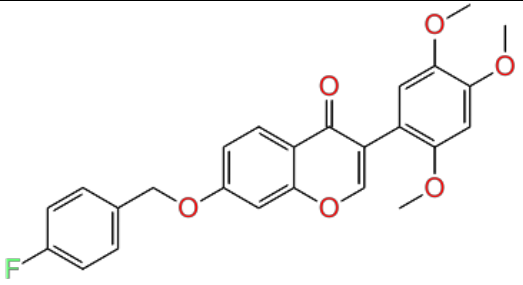 | Moderate | 91 | training | 5.50 | 5.03 | 1.09 |

| | | | | | | | | |
|---|---|---|---|---|---|---|---|---|
| 52. | CHEMBL3330999 | 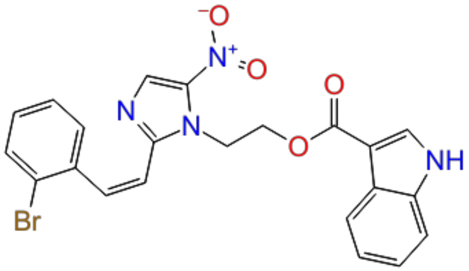 | Moderate | 151 | test | 5.51 | 6.09 | 1.87 |
| 53. | CHEMBL3331010 | 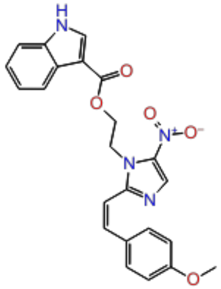 | Moderate | 107 | training | 5.51 | 5.75 | 1.33 |
| 54. | CHEMBL3330997 | 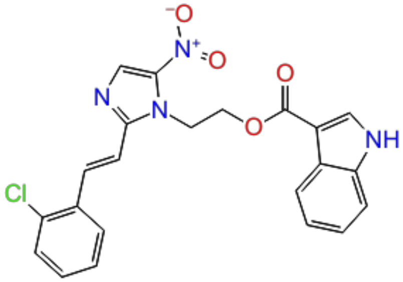 | Moderate | 84 | training | 5.53 | 5.03 | 1.07 |
| 55. | CHEMBL3331003 | 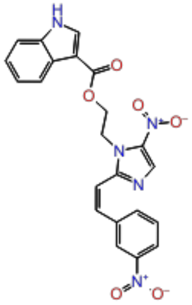 | Moderate | 182 | training | 5.55 | 5.2 | 1.16 |

| | | | | | | | | |
|---|---|---|---|---|---|---|---|---|
| 56. | CHEMBL3311052 | 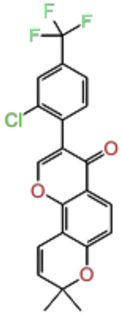 | Moderate | 4 | test | 5.60 | 4.93 | 1.15 |
| 57. | CHEMBL3331004 | 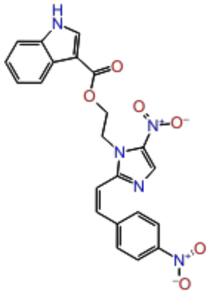 | Moderate | 114 | training | 5.69 | 5.62 | 1.15 |
| 58. | CHEMBL1164847 | 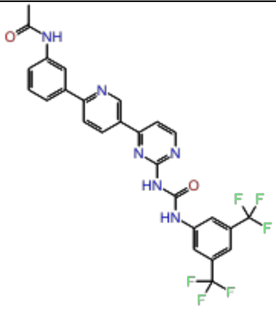 | Moderate | 255 | training | 5.70 | 4.47 | 0.92 |

| | | | | | | | | |
|---|---|---|---|---|---|---|---|---|
| 59. | CHEMBL1271866 | 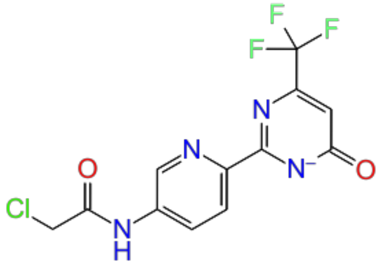 | Moderate | 12 | test | 5.82 | 5.33 | 0.54 |
| 60. | CHEMBL3331002 | 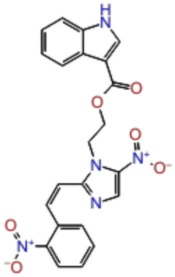 | Moderate | 185 | training | 5.96 | 5.61 | 1.15 |
| 61. | CHEMBL3331011 | 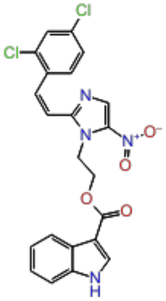 | Moderate | 137 | training | 6.06 | 6.58 | 1.78 |

| | | | | | | | | |
|---|---|---|---|---|---|---|---|---|
| 62. | CHEMBL1956190 | 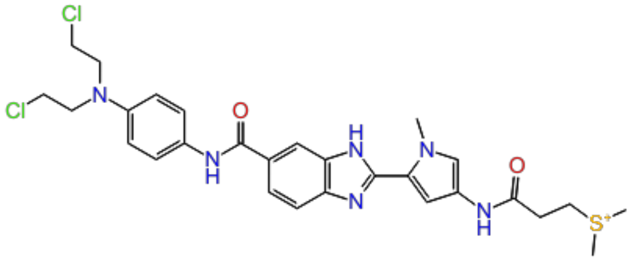 | Moderate | 607 | test | 6.15 | 5.93 | 1.25 |
| 63. | CHEMBL53463 | 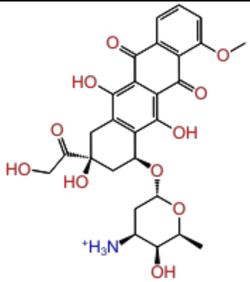 | Moderate | 26 | training | 6.22 | 5.08 | 0.96 |
| 64. | CHEMBL84 | 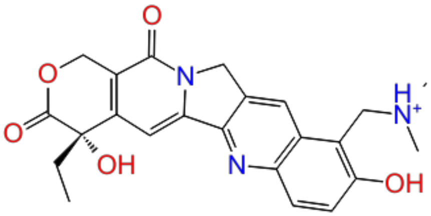 | Moderate | 18 | training | 6.30 | 6.34 | 0.88 |
| 65. | CHEMBL3311160 | 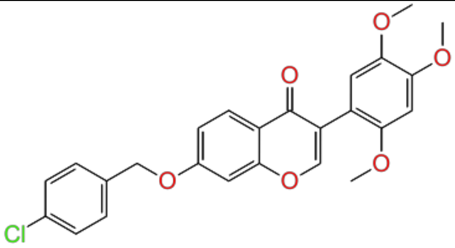 | Moderate | 87 | test | 6.33 | 4.89 | 1.1 |

| | | | | | | | | |
|---|---|---|---|---|---|---|---|---|
| 66. | CHEMBL1278024 | 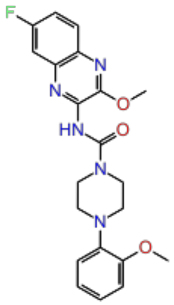 | Moderate | 133 | training | 6.85 | 7.02 | 1.89 |
| 67. | CHEMBL3758224 | 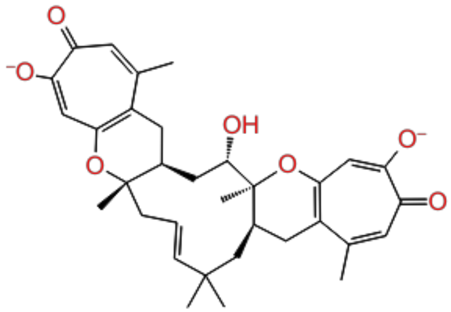 | active | 7 | training | 7.00 | 7.27 | 1.18 |
| 68. | CHEMBL1276954 | 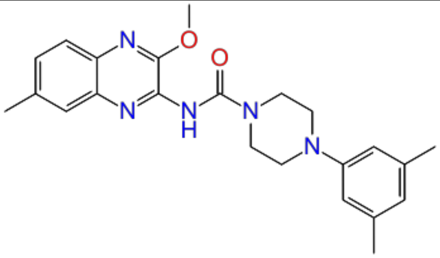 | active | 96 | test | 7.07 | 7.69 | 2.94 |
| 69. | CHEMBL1277041 | 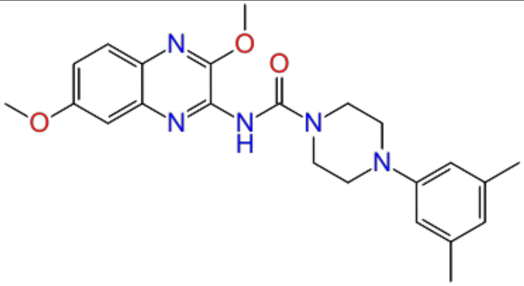 | active | 128 | training | 7.12 | 7.6 | 3 |

| | | | | | | | | |
|---|---|---|---|---|---|---|---|---|
| 70. | CHEMBL1276866 | 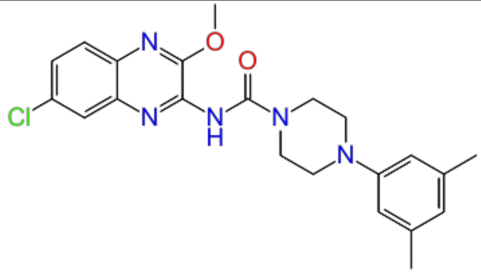 | active | 96 | training | 7.16 | 7.81 | 2.93 |
| 71. | CHEMBL1277939 | 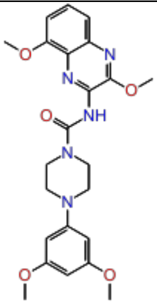 | active | 132 | test | 7.24 | 7.22 | 2.69 |
| 72. | CHEMBL1276865 | 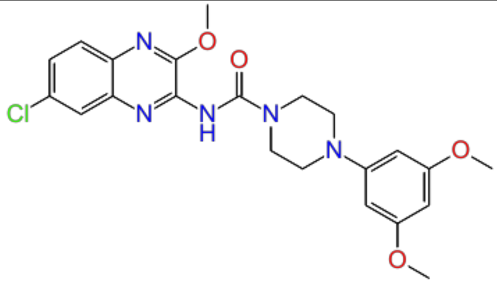 | active | 152 | training | 7.25 | 7.25 | 2.76 |
| 73. | CHEMBL1278119 | 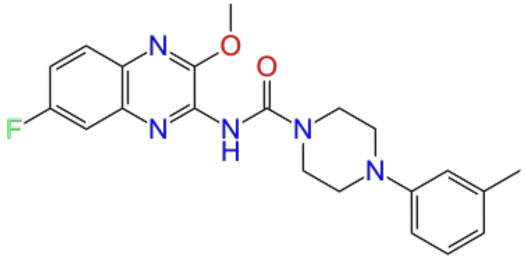 | active | 159 | training | 7.30 | 7.79 | 2.86 |

| | | | | | | | | |
|---|---|---|---|---|---|---|---|---|
| 74. | CHEMBL1276953 | 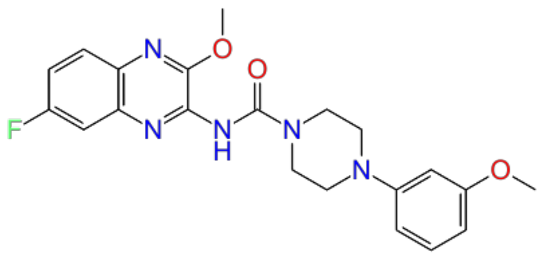 | active | 174 | test | 7.51 | 7.46 | 2.74 |
| 75. | CHEMBL3338195 | 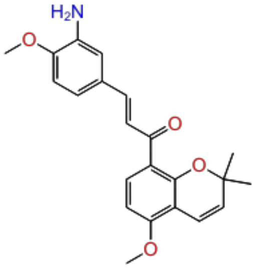 | active | 59 | training | 7.62 | 6.73 | 1.61 |
| 76. | CHEMBL428647 | 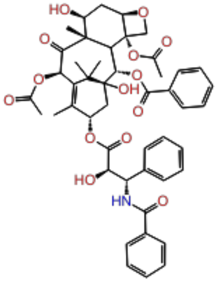 | active | 75 | training | 7.64 | 7.18 | 0.69 |
| 77. | CHEMBL1278205 | 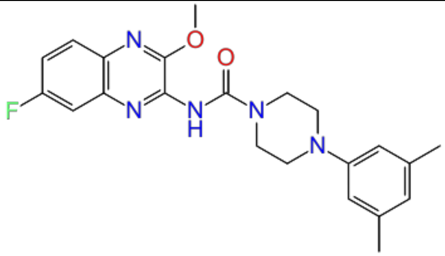 | active | 98 | test | 7.72 | 7.6 | 2.96 |

| | | | | | | | | |
|---|---|---|---|---|---|---|---|---|
| 78. | CHEMBL1278118 | 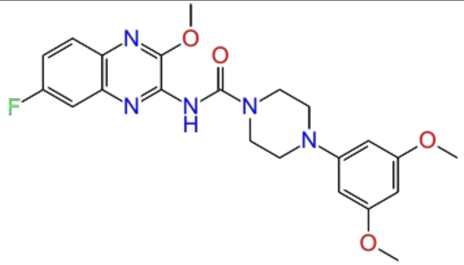 | active | 161 | training | 7.82 | 7.77 | 2.75 |
| 79. | CHEMBL1683544 | 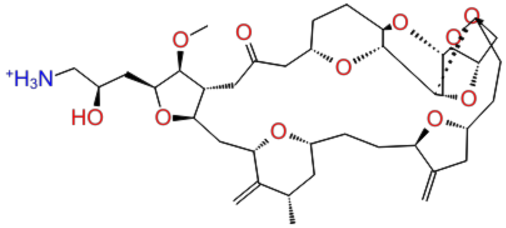 | active | 139 | training | 8.23 | 8 | 1.4 |
| 80. | CHEMBL1683553 | 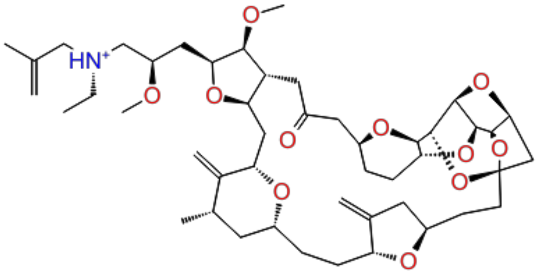 | active | 130 | test | 8.43 | 7.67 | 1.34 |
| 81. | CHEMBL1683552 | 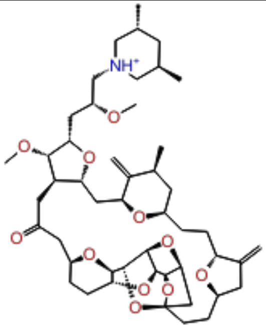 | active | 158 | training | 8.59 | 7.81 | 1.32 |

| | | | | | | | | |
|---|---|---|---|---|---|---|---|---|
| 82. | CHEMBL2420629 | 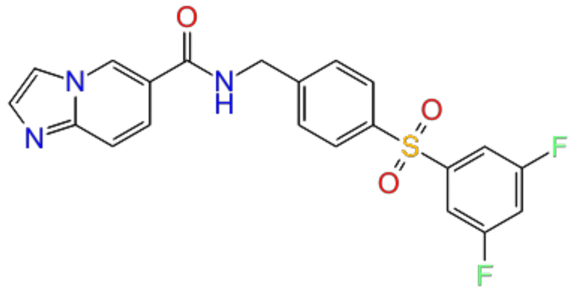 | active | 41 | training | 8.74 | 7.85 | 1.44 |
| 83. | CHEMBL1683546 | 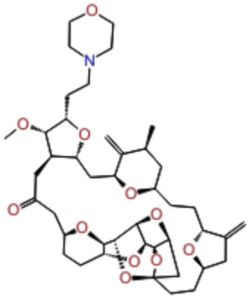 | active | 108 | test | 8.90 | 7.82 | 1.43 |
| 84. | CHEMBL1683523 | 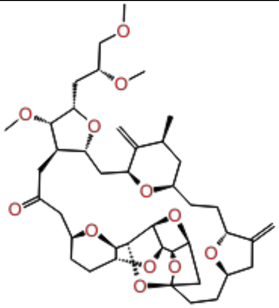 | active | 107 | training | 9.13 | 7.94 | 1.35 |

| | | | | | | | | |
|---|---|---|---|---|---|---|---|---|
| 85. | CHEMBL1683549 | 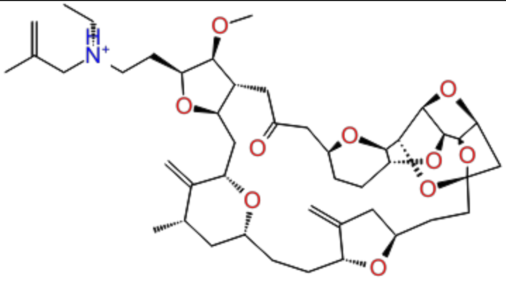 | active | 227 | training | 9.21 | 8.44 | 1.31 |
| 86. | CHEMBL1683550 | 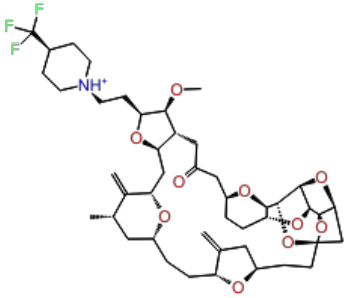 | active | 103 | test | 9.55 | 8.18 | 1.27 |
| 87. | CHEMBL1683556 | 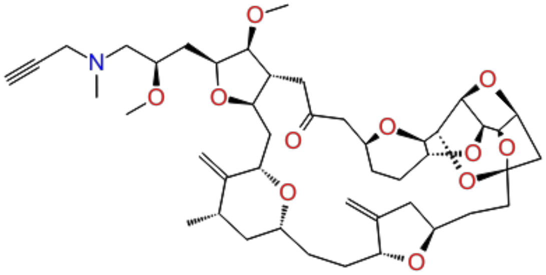 | active | 115 | training | 9.74 | 8.5 | 1.38 |

**Table S12:** Inhibitors against U251 cell line used as an external test set for the generated PHASE model.

| Sr. No | Compound ID | 2D structure | $pIC_{50}$ |
|---|---|---|---|
| 1. | CHEMBL1958321 | 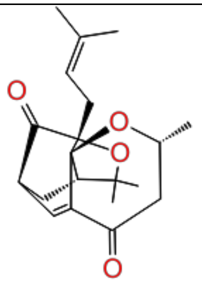 | 4.08 |
| 2. | CHEMBL252568 | 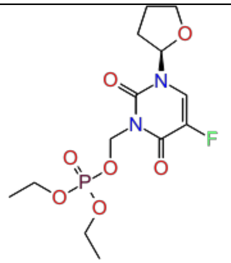 | 4.14 |
| 3. | CHEMBL452926 | 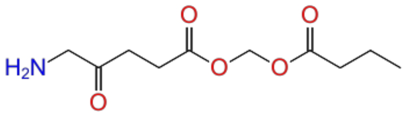 | 4.23 |
| 4. | CHEMBL467590 | 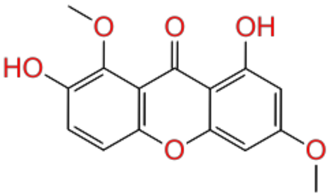 | 4.27 |

| | | | |
|---|---|---|---|
| 5. | CHEMBL3357146 | 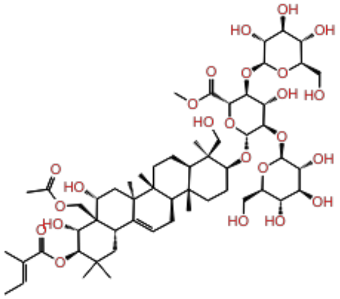 | 4.32 |
| 6. | CHEMBL364713 | 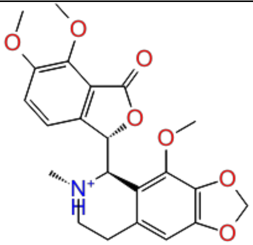 | 4.40 |
| 7. | CHEMBL3357167 | 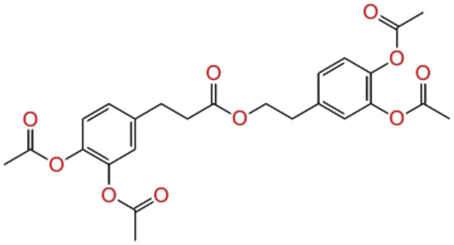 | 4.43 |
| 8. | CHEMBL45240 | 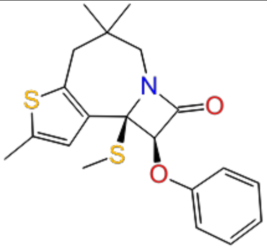 | 4.48 |

| | | | |
|---|---|---|---|
| 9. | CHEMBL1784163 | 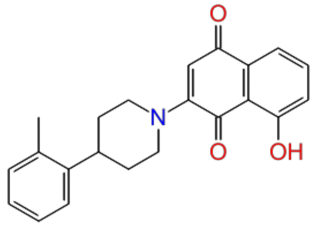 | 4.52 |
| 10. | CHEMBL178297 | 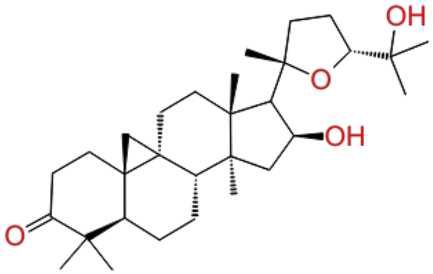 | 4.56 |
| 11. | CHEMBL3357166 | 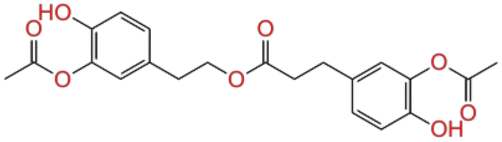 | 4.57 |
| 12. | CHEMBL1958316 | 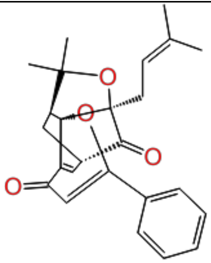 | 4.65 |

| | | | |
|---|---|---|---|
| 13. | CHEMBL180924 | 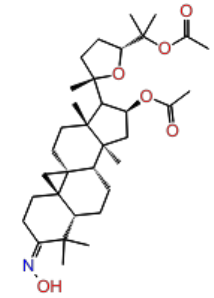 | 4.67 |
| 14. | CHEMBL179443 | 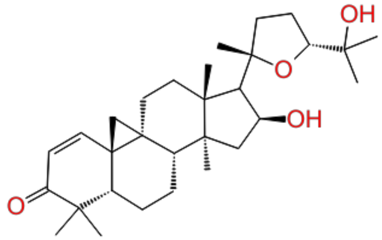 | 4.72 |
| 15. | CHEMBL3331345 | 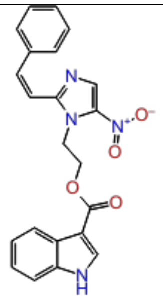 | 4.77 |
| 16. | CHEMBL3357147 | 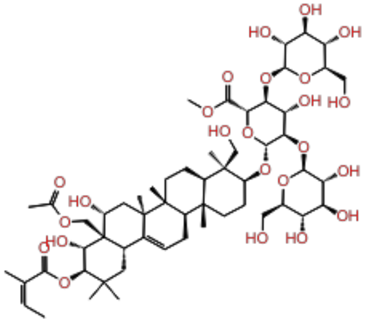 | 4.81 |

| | | | |
|---|---|---|---|
| 17. | CHEMBL178305 | 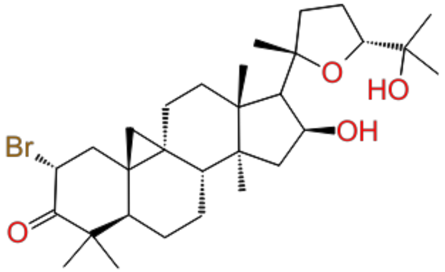 | 4.86 |
| 18. | CHEMBL3588937 | 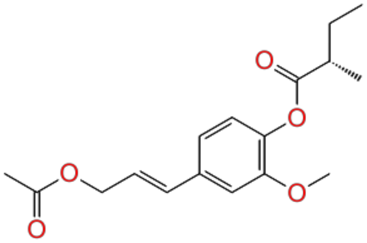 | 4.89 |
| 19. | CHEMBL1784159 | 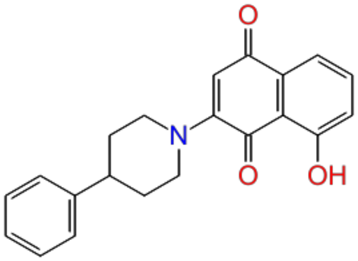 | 4.97 |
| 20. | CHEMBL478578 | 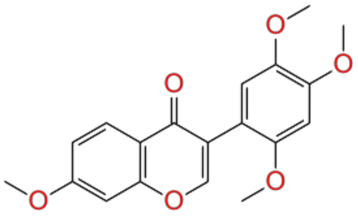 | 5.03 |

| | | | |
|---|---|---|---|
| 21. | CHEMBL1958320 | 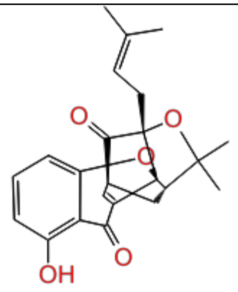 | 5.04 |
| 22. | CHEMBL1958314 | 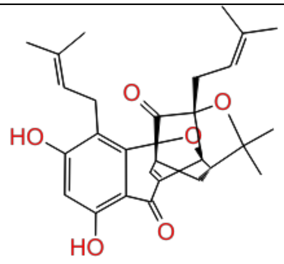 | 5.07 |
| 23. | CHEMBL4065827 | 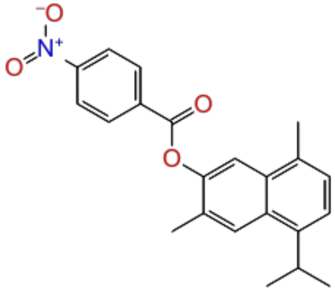 | 5.11 |
| 24. | CHEMBL498233 | 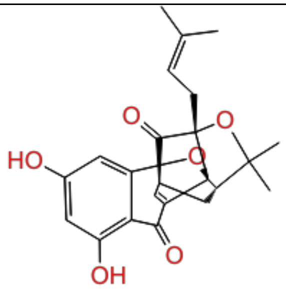 | 5.13 |

| | | | |
|---|---|---|---|
| 25. | CHEMBL1784157 | 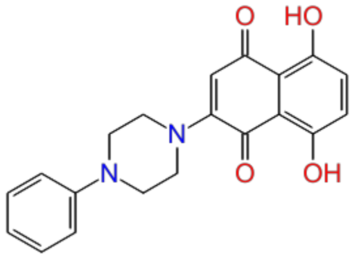 | 5.21 |
| 26. | CHEMBL3311053 | 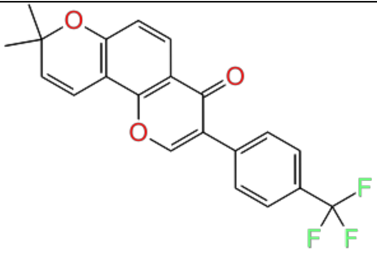 | 5.23 |
| 27. | CHEMBL1784155 | 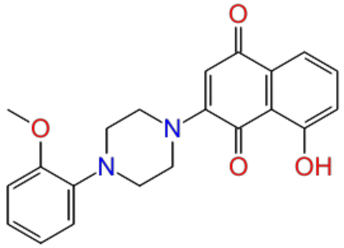 | 5.36 |
| 28. | CHEMBL3331008 | 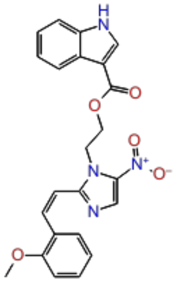 | 5.39 |

| | | | |
|---|---|---|---|
| 29. | CHEMBL3331000 | 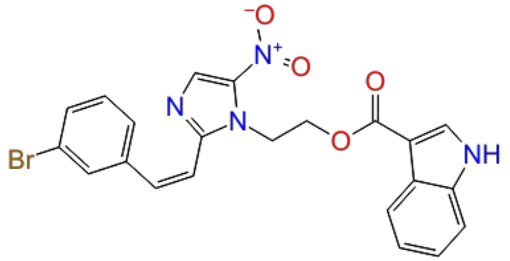 | 5.41 |
| 30. | CHEMBL3311165 | 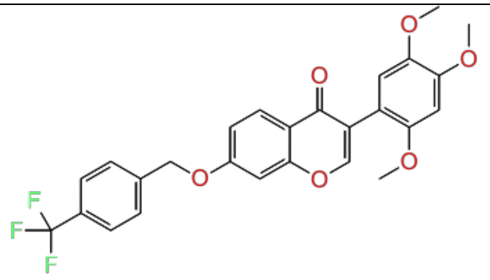 | 5.43 |
| 31. | CHEMBL3311159 | 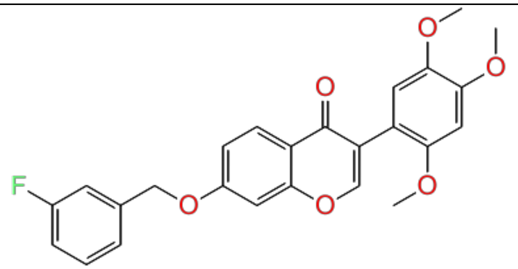 | 5.46 |
| 32. | CHEMBL3331007 | 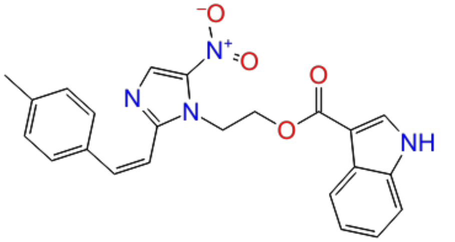 | 5.51 |

| | | | |
|---|---|---|---|
| 33. | CHEMBL3331006 | 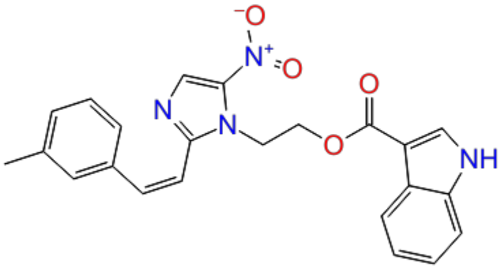 | 5.52 |
| 34. | CHEMBL3311150 | 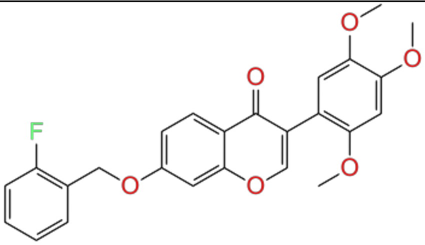 | 5.54 |
| 35. | CHEMBL3331005 | 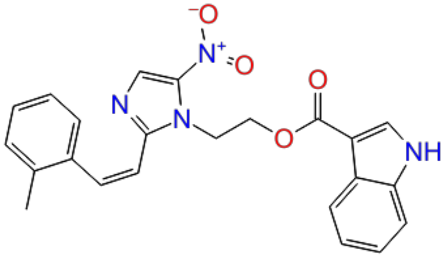 | 5.55 |
| 36. | CHEMBL3331009 | 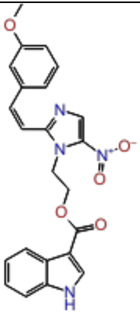 | 5.56 |

| | | | |
|---|---|---|---|
| 37. | CHEMBL555017 | 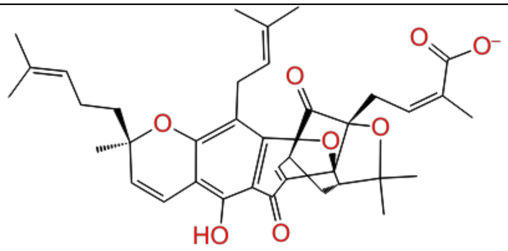 | 5.59 |
| 38. | CHEMBL3400386 | 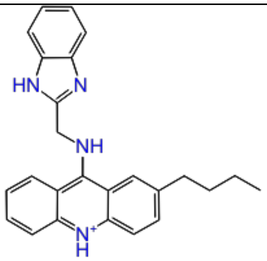 | 5.62 |
| 39. | CHEMBL3311153 | 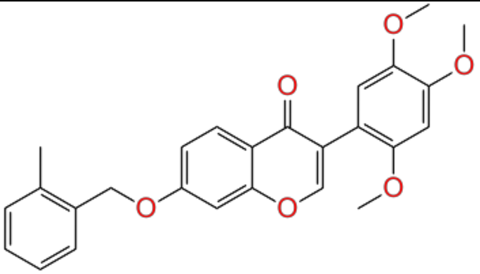 | 5.70 |
| 40. | CHEMBL61619 | 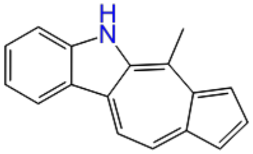 | 5.83 |

| | | | |
|---|---|---|---|
| 41. | CHEMBL3311158 | 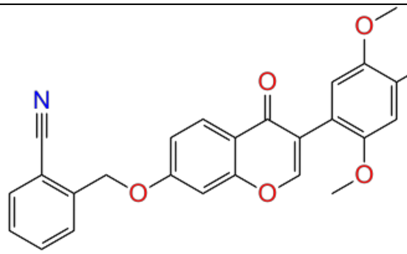 | 6.01 |
| 42. | CHEMBL2064455 | 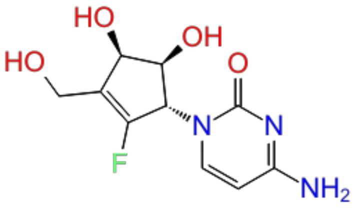 | 6.08 |
| 43. | CHEMBL3408446 | 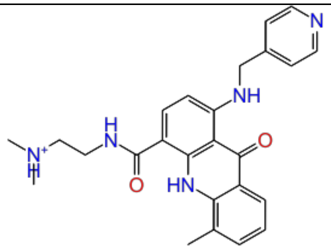 | 6.10 |
| 44. | CHEMBL67 | 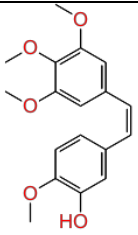 | 6.27 |

| | | | |
|---|---|---|---|
| 45. | CHEMBL4066486 | 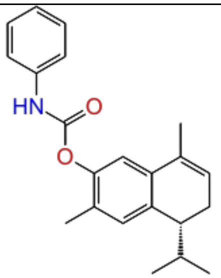 | 6.34 |
| 46. | CHEMBL1278206 | 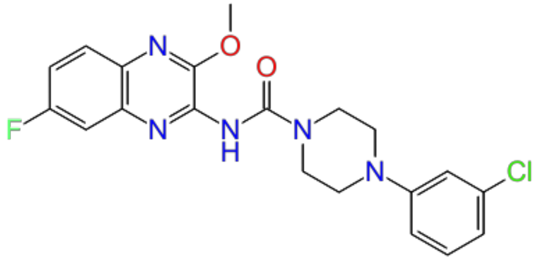 | 7.10 |
| 47. | CHEMBL1278025 | 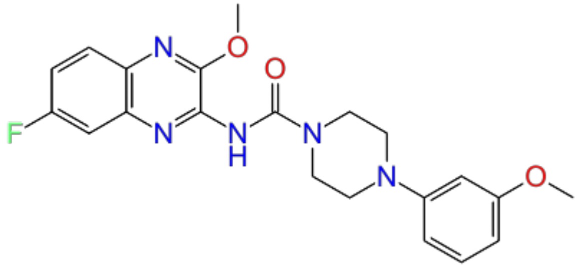 | 7.16 |
| 48. | CHEMBL1277938 | 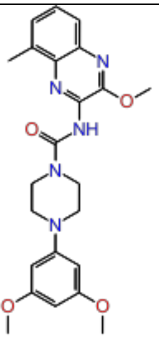 | 7.27 |

| | | | |
|---|---|---|---|
| 49. | CHEMBL1277042 | 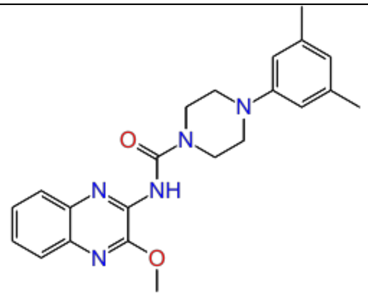 | 7.52 |
| 50. | CHEMBL107 | 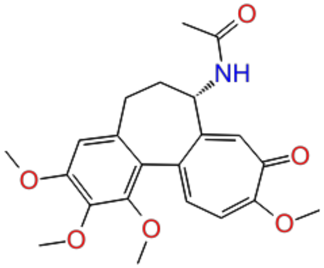 | 7.70 |
| 51. | CHEMBL1683547 | 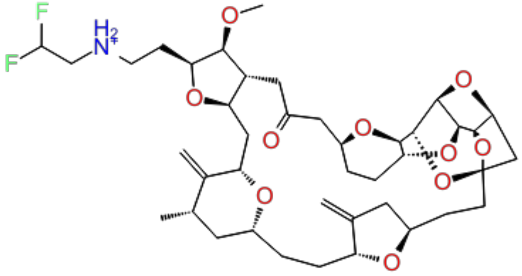 | 8.39 |
| 52. | CHEMBL1683554 | 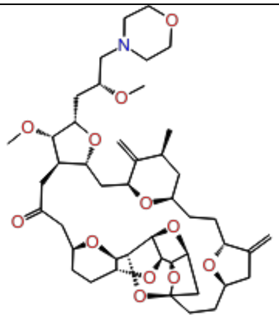 | 8.71 |

| | | | |
|---|---|---|---|
| 53. | CHEMBL1683548 | 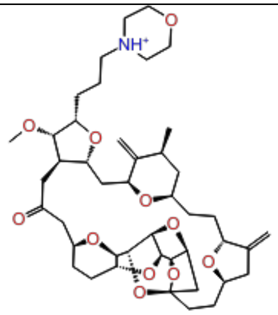 | 8.84 |
| 54. | CHEMBL1683551 | 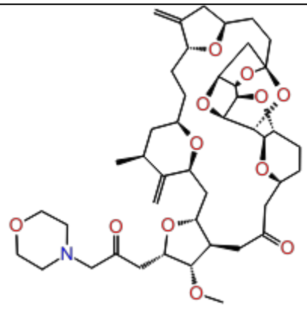 | 9.06 |
| 55. | CHEMBL1683545 | 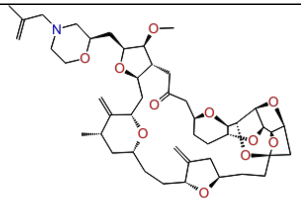 | 9.30 |
| 56. | CHEMBL1683555 | 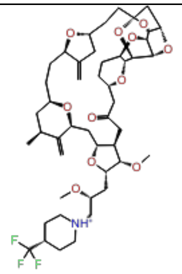 | 9.48 |
